# Exploring the value of thematic urban ecosystem accounts in Western Balkan Countries to inform urban greening actions

**DOI:** 10.64898/2026.01.02.697364

**Authors:** Javier Babí Almenar, Claudia Romelli, Saša Čegar, Kejt Dhrami, Saša Drezgić, Fiona Imami, Matjaz Hribar, Vesna Janković Milić, Sonja Jovanović, Vanja Krmelj, Milena Lipovina-Bozovic, Jelena Stanković, Žiga Turk, Simona Muratori, Renato Casagrandi

## Abstract

Urban ecological health is increasingly prioritized in Europe, as reflected in the EU Biodiversity Strategy 2030, the Nature Restoration Regulation, and the amendment to the Regulation on Environmental-Economic Accounts. Translating this ambition into greening actions that halt and reverse urban ecological degradation requires standardized, spatially explicit indicators. This study explores the utility of the System of Environmental-Economic Accounting–Ecosystem Accounting (SEEA-EA) framework for developing thematic urban ecosystem accounts as integrated metrics to inform urban greening in the Western Balkans. We constructed pilot accounts for the Western Balkans for 2018, with particular focus on Horizon-funded CROSS-REIS countries, where local accounts were also developed for representative municipalities (Kranj, Rijeka, Niš, Podgorica, Tirana). The accounts quantify ecosystem extent, four condition metrics (imperviousness, green space, tree cover, and PM₁₀ concentration), and the service air filtration using PM₁₀ deposition onto vegetation as a proxy. Results were benchmarked nationally against EU/EFTA/UK countries and locally against Italian municipalities of the National Biodiversity Future Centre project. The analysis reveals that urban ecosystems in the Western Balkans exist on a continuum with other European countries but exhibit a distinct, more homogeneous profile shaped by geography, climate, and policy legacies. The accounts identify ecological degradation hotspots (e.g., high soil sealing) and areas of high service-delivery capacity, supporting translation of spatial diagnosis into targeted greening priorities. Overall, the study shows that thematic urban ecosystem accounts offer a standardized, multi-scale evidence base to inform and monitor urban greening policies, supporting the Western Balkans’ environmental management needs on their path toward EU integration.

## 1. Introduction

Over the past decade, urban (re)greening has attracted renewed global attention, particularly within the European Union (EU) where many initiatives and applied actions have been developed (Varbova, 2022; Wild et al., 2020). This trend has been driven by increased research investment and the emergence of concepts like urban nature-based solutions, which promise net environmental, economic, and social benefits (Frantzeskaki et al., 2020; Grace et al., 2021). A growing body of evidence highlights the potential of these solutions to address specific urban challenges, such as those derived from climate change (e.g., exacerbation of the urban heat island effect), physical and mental health issues, and biodiversity loss (Cohen-Shacham et al., 2016; Eggermont et al., 2015; Keeler et al., 2019). In urban contexts, biodiversity loss is closely linked to land degradation and land-use change processes that drive the expansion of soil sealing and the loss of non-artificial land covers and their structural components (e.g., trees and shrubs), resulting in habitat loss and fragmentation (Fairbairn et al., 2024; Ferreira et al., 2018; Liu et al., 2016). While nature-based solutions are increasingly advocated as responses to the above challenges, several scholars caution against treating them as a panacea by assuming inherent benefits in all cases (Babí Almenar et al., 2021; Torres et al., 2023). These critiques have reinforced calls for more comprehensive, transparent, and (where feasible) standardized frameworks to assess and monitor urban nature-based solutions and greening actions in general (Alves Prado and Flauzino Pires, 2025; Pierce et al., 2024).

In parallel to research developments, EU policy on urban nature has undergone substantial evolution in recent years, reflecting a shift from general recognition of ecosystem benefits toward more actionable, and in some cases legally binding, targets (Perez-Soba et al., 2025). Early policy instruments acknowledged the role of urban nature in climate adaptation, e.g., EU White Paper Adapting to Climate Change (European Commission, 2009), and promoted green infrastructure approaches, e.g., EU Green Infrastructure Strategy 2012 (European Commission, 2013) and EU Biodiversity Strategy 2020 (European Commission, 2011). This trajectory has culminated in the recent adoption of the Nature Restoration Regulation (NRR), which explicitly seeks to reverse land degradation and restore ecosystems as a means to halt biodiversity loss (European Parliament, Council of the European Union, 2024a). Complementarily, the EU Biodiversity Strategy for 2030 mandates the development of “ambitious” urban greening plans for all municipalities with more than 20,000 inhabitants (European Commission, 2020). As a result, EU Member States are now required to translate these commitments into national and local policy instruments. Then, monitoring systems that are comprehensive, coherent, and comparable across jurisdictions are needed. This requirement is equally pertinent for countries in the Western Balkans, which are expected to progressively integrate into the EU (European Union External Action, 2025), and will therefore need to adopt compatible approaches to urban ecosystem monitoring and assessment.

Ecosystem accounting provides a promising decision-support framework for meeting these policy and monitoring demands by systematically recording ecosystem extent, condition, service flows, and monetary ecosystem asset values (Comte et al., 2022; Hein et al., 2015). The System of Environmental-Economic Accounting–Ecosystem Accounting (SEEA-EA), developed by the United Nations, constitutes the global statistical standard for this purpose (United Nations, 2021). Importantly, the SEEA-EA is applicable to both natural and anthropogenic ecosystems, including urban systems, which encompass settlements as well as surrounding peri-urban buffers of semi-natural and natural ecosystems (Vallecillo et al., 2022). In the context of accelerating land degradation and biodiversity loss, standardized accounting frameworks such as SEEA-EA provide a spatially explicit evidence foundation for linking policy goals to targeted actions. By tracking changes in ecosystem extent and condition, urban ecosystem accounts can directly capture certain dimensions of urban ecological degradation, such as the loss of green space and their heterogeneity, the expansion of impervious surfaces, and the reduction in biodiversity-supporting structures, e.g., tree cover. Accordingly, our paper proposes to use urban ecosystem accounting as a relevant tool for monitoring urban ecological health, and for informing urban greening policies aimed at mitigating and reversing it.

At the global level, an emerging ensemble of studies are testing the utility of urban ecosystem accounts for policy and planning purposes (e.g., Barton, 2023; Heris et al., 2021), with some explicitly addressing challenges related to standardization and comparability (e.g., Babí Almenar et al., 2026, 2023; Venter et al., 2024). As an example, Heris et al. (2021) conducted a research in the United States, with municipal-scale accounts developed for cities with populations exceeding 50.000, focusing on land cover extent, rainfall interception, and local climate regulation. In Australia, exploratory urban ecosystem accounting initiatives have been led by local and regional authorities, with early pilot projects documenting both opportunities and methodological challenges (Cryle et al., 2021). Within the EU, policy developments have also accelerated interest in urban ecosystem accounting. A recent regulation (European Parliament, Council of the European Union, 2024b) mandates the regular compilation of SEEA-EA-aligned accounts, including indicators such as green space share, air pollution (PM_10_, PM _2.5_) concentrations, and local climate regulation services for urban areas. The regulation also requires ecosystem service accounts at national level for services relevant to urban contexts, including global climate regulation and air filtration (European Parliament, Council of the European Union, 2024b). Reflecting this agenda, the European Commission’s Joint Research Centre recently published a pilot study on urban ecosystem accounts for EU and EEA countries (Babí Almenar et al., 2023). However, that work did not extend to prospective EU Member States, including those in the Western Balkans.

Existing pilot studies, together with ongoing methodological discussions, underscore the need for further testing and refinement of urban ecosystem accounts, particularly in regions where implementation has not yet occurred (Babí Almenar et al., 2023; La Notte and Zulian, 2021). Expanding the empirical base can reveal context-specific challenges, enhance policy relevance, support methodological harmonization, and foster communities of practice (Babí Almenar et al., 2026). Such efforts also may create opportunities for knowledge exchange among researchers, practitioners, and institutions across countries and governance levels. Consequently, additional exploratory applications of urban ecosystem accounting remain both necessary and timely.

Our study investigates the utility of pilot thematic urban ecosystem accounts for informing urban green actions in the Western Balkans that halt and reverse urban ecological degradation, with a particular attention to countries participating to the Horizon-funded project CROSS-REIS (http://www.crossreis.com): Albania, Croatia, Montenegro, Serbia and Slovenia. At the local level, one representative municipality per CROSS-REIS country (Tirana, Rijeka, Podgorica, Niš and Kranj) was selected as a pilot city where to deepen the analysis.

The aim of this study is pursued through four connected, yet independent objectives:

i. To develop the first SEEA-EA-aligned urban ecosystem accounts for Western Balkan countries, consistent with approaches applied in EU and EEA Member States.
ii. To benchmark the national ecosystem accounting results of Western Balkan countries against each other and against those of EU, EFTA, and UK countries.
iii. To benchmark ecosystem accounting outcomes for pilot CROSS-REIS municipalities, against similar Italian pilot municipalities of the project *National Biodiversity Future Centre* (http://www.nbfc.it).
iv. To discuss the usefulness of urban ecosystem accounts for tracking ecological degradation in urban ecosystems and for informing urban greening policies aimed at reverting it.

This study develops novel urban ecosystem accounts based exclusively on publicly available Earth observation data, aligned with the SEEA-EA framework. The accounts cover ecosystem extent, four ecosystem condition variables (green space, tree cover, impervious surfaces, and particulate matter concentration), and one ecosystem service (air filtration by vegetation) for 2018. It is important to note that national-level analysis includes all Western Balkan countries, not only the CROSS-REIS participants.

## 2. Methods

### 2.1. Definition of reporting units and the ecosystem accounting area

For the development of urban ecosystem accounts, we follow the recommendations from SEEA-EA thematic ecosystem accounts, a pilot urban ecosystem accounting exercise for the EU and EEA illustrated in Babí Almenar et al.(2023), and insights from a recent comprehensive review of challenges and lessons in urban ecosystem accounting (Babí Almenar et al., 2026). The SEEA-EA encourages the development of thematic accounts for policy-relevant environmental themes, such as biodiversity, oceans, and urban areas, which are not part of its core standards or recommendations (Edens et al., 2022; United Nations, 2021). Within SEEA-EA thematic accounts, urban areas are generally qualified as “urban ecosystems”. They are defined as heterogeneous fine-grained landscape mosaics of patches (ecosystem assets) with fuzzy boundaries that comprise multiple ecosystem types (Grimm et al., 2008; Wu, 2014). In other words, the diverse ecosystem assets within settlements, together with their semi-natural or natural buffers, collectively represent an urban ecosystem. Thus, single urban ecosystems serve as individual reporting units within the ecosystem accounting area.

Following the approach of the pilot EU/EEA urban ecosystem accounts (Babí Almenar et al., 2023) and relevant EU regulations, this study uses Local Administrative Units (LAUs) classified as ‘cities’ or ‘towns and suburbs’ by the Degree of Urbanization (Dijkstra et al., 2021), as its core reporting units, herein referred to as ‘urban LAUs’. In most European countries, LAUs correspond to municipalities, with exceptions such as Ireland and Portugal. This definition aligns with the methodology adopted by recent amendments to the Regulation on Environmental-Economic Accounts and the Nature Restoration Regulation for delineating urban ecosystems (European Parliament, Council of the European Union, 2024b, 2024a). Our study area’s LAU boundaries were compiled by merging the European Commission’s 2020 Geographic Information System (GISCO) dataset (https://ec.europa.eu/eurostat/web/gisco/geodata) with the LAUs data from the Global Administrative Areas (GADM) platform (https://gadm.org) for Bosnia and Herzegovina, Kosovo and Montenegro. The spatial analysis used the Joint Research Centre’s 2018 population grid (Batista e Silva et al., 2021) as its primary input, supplemented by the WorldPop dataset for parts of Cyprus not covered by the JRC grid. Only urban LAUs were retained for the final analysis.

The selection was further refined and urban LAUs were excluded if they had (i) fewer than 1,000 inhabitants or (ii) an overall population density below 10 inhabitants per km². This step ensured the removal of LAUs that, although classified as “urban” under the Degree of Urbanization, extend beyond the conceptual limits of urban ecosystems. For example, large LAUs containing a very small share of urban territory. Similar thresholds have also been applied in previous studies with analogous aims (Babí Almenar et al., 2023; Zulian et al., 2022). Figure 1 shows the resulting refined set of reporting units, i.e. the corresponding entire urban ecosystem accounting area, in the Western Balkans together with those of European counterparts.

**Figure 1.**
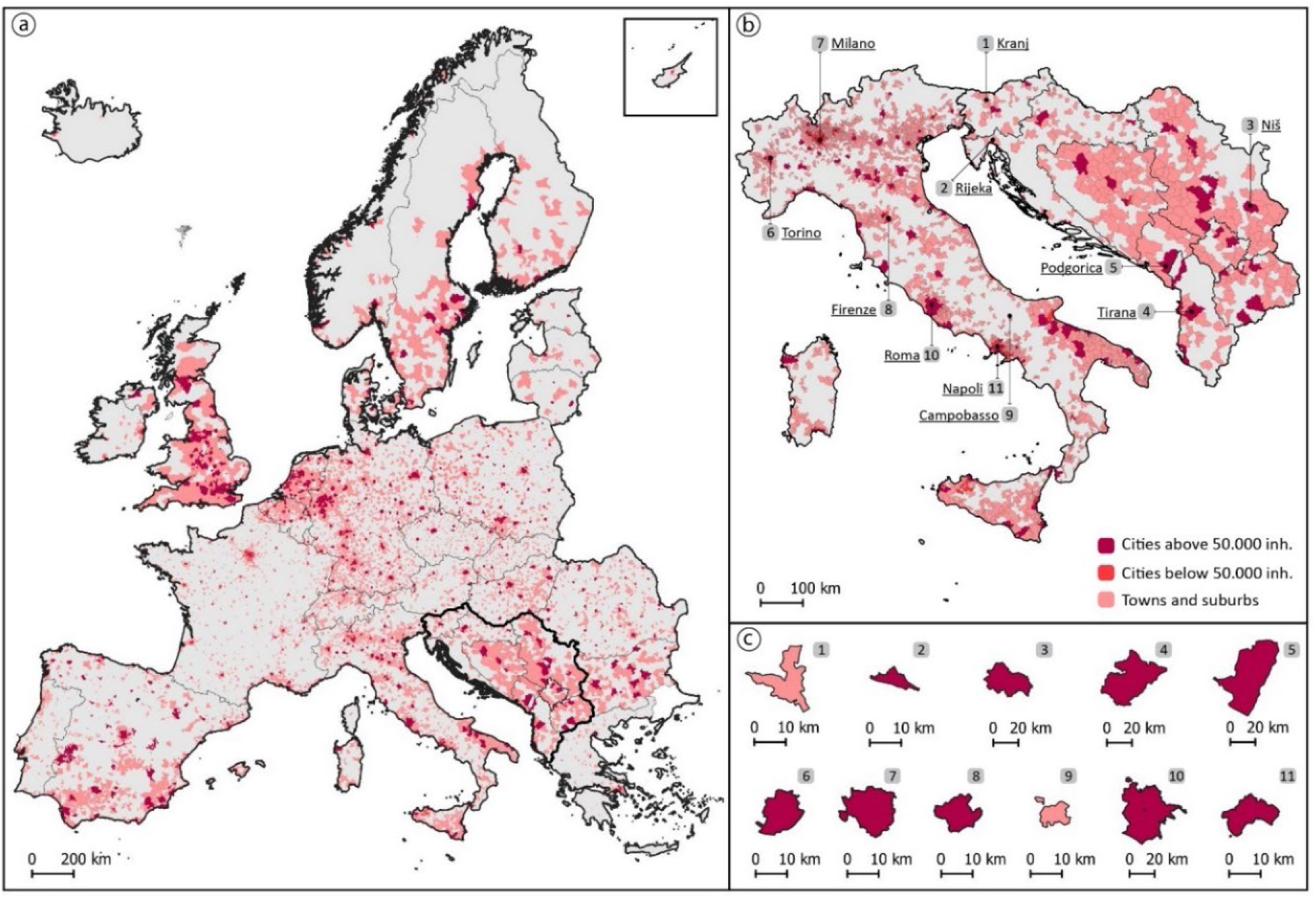
**Ecosystem accounting area for the thematic urban ecosystem accounts reported in this study and their reporting units* (a) Ecosystem accounting area at the continental-scale, full area of scope**; (b) Country-scale view of Italy and the Western Balkans, highlighting the pilot cities; (c) Detailed zoomed-in view of each pilot city.** *Local administrative areas classified as cities (above and below 50,000 inhabitants) and towns and suburbs, which represent urban ecosystems. **Area of scope: Urban ecosystems within the European Union, European Free Trade Association Countries, the UK and the Western Balkans. For representation purposes, Cyprus is shown within a rectangle in the upper right corner in (a).

### 2.2. Urban ecosystem extent accounts

In thematic urban ecosystem accounts, extent accounts measure the distribution of ecosystem assets across the different ecosystem types within each reporting unit. In this study, the classification of ecosystem types follows the EU Ecosystem Typology at Level 1 (see Table 1), as required by the amendment to the Regulation on Environmental-Economic Accounts (European Parliament, Council of the European Union, 2024b).

**Table 1.**
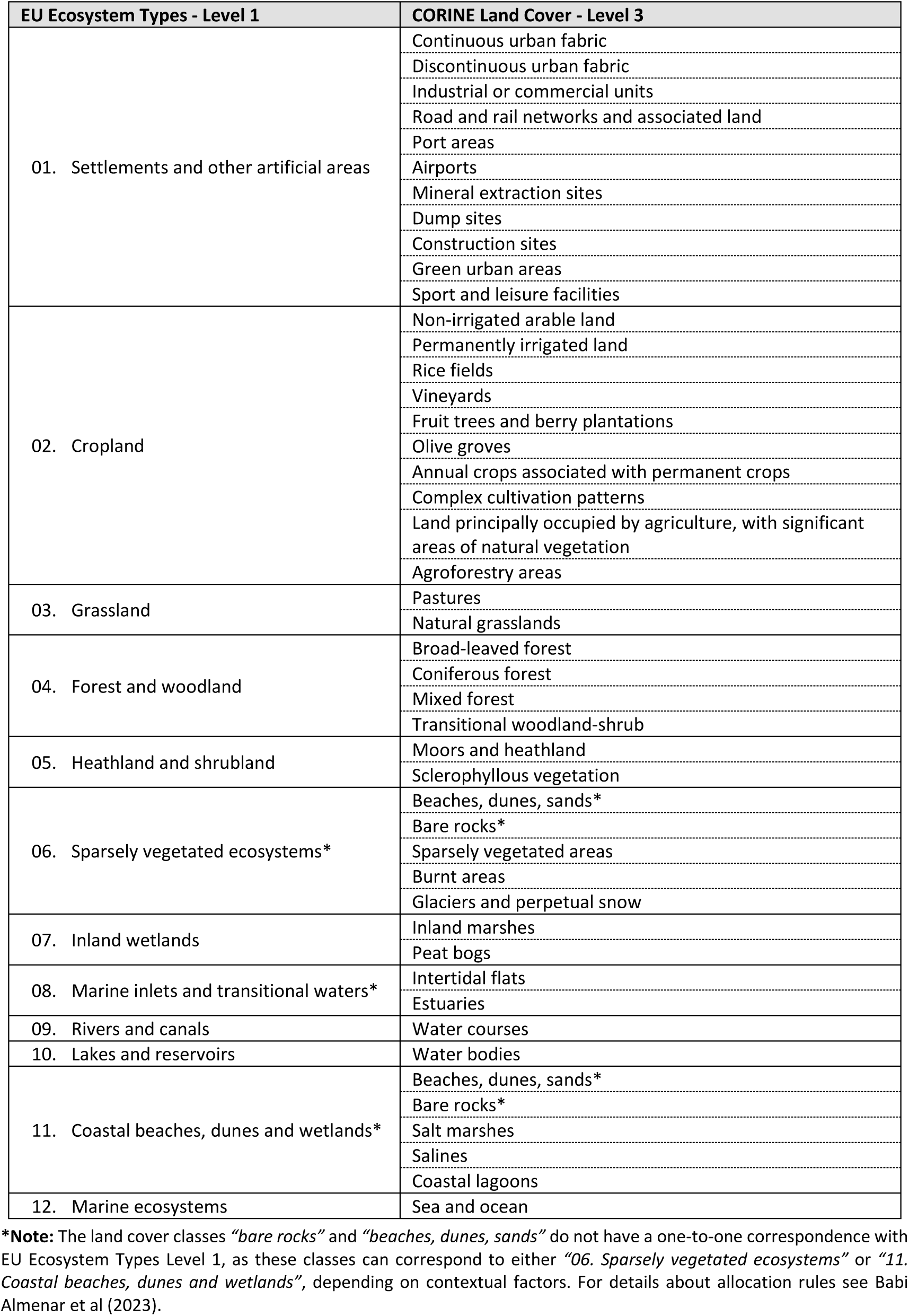
EU Ecosystem Typology Level 1 with the correspondence to CORINE land cover classes Level 3.

| EU Ecosystem Types - Level 1 | CORINE Land Cover - Level 3 |
| --- | --- |
| 01. Settlements and other artificial areas | Continuous urban fabric |
|  | Discontinuous urban fabric |
|  | Industrial or commercial units |
|  | Road and rail networks and associated land |
|  | Port areas |
|  | Airports |
|  | Mineral extraction sites |
|  | Dump sites |
|  | Construction sites |
|  | Green urban areas |
|  | Sport and leisure facilities |
| 02. Cropland | Non-irrigated arable land |
|  | Permanently irrigated land |
|  | Rice fields |
|  | Vineyards |
|  | Fruit trees and berry plantations |
|  | Olive groves |
|  | Annual crops associated with permanent crops |
|  | Complex cultivation patterns |
|  | Land principally occupied by agriculture, with significant areas of natural vegetation |
|  | Agroforestry areas |
| 03. Grassland | Pastures |
|  | Natural grasslands |
| 04. Forest and woodland | Broad-leaved forest |
|  | Coniferous forest |
|  | Mixed forest |
|  | Transitional woodland-shrub |
| 05. Heathland and shrubland | Moors and heathland |
|  | Sclerophyllous vegetation |
| 06. Sparsely vegetated ecosystems* | Beaches, dunes, sands* |
|  | Bare rocks* |
|  | Sparsely vegetated areas |
|  | Burnt areas |
|  | Glaciers and perpetual snow |
| 07. Inland wetlands | Inland marshes |
|  | Peat bogs |
| 08. Marine inlets and transitional waters* | Intertidal flats |
|  | Estuaries |
| 09. Rivers and canals | Water courses |
| 10. Lakes and reservoirs | Water bodies |
| 11. Coastal beaches, dunes and wetlands* | Beaches, dunes, sands* |
|  | Bare rocks* |
|  | Salt marshes |
|  | Salines |
|  | Coastal lagoons |
| 12. Marine ecosystems | Sea and ocean |

**\*Note:** The land cover classes “bare rocks” and “beaches, dunes, sands” do not have a one-to-one correspondence with EU Ecosystem Types Level 1, as these classes can correspond to either “06. Sparsely vegetated ecosystems” or “11. Coastal beaches, dunes and wetlands”, depending on contextual factors. For details about allocation rules see Babi Almenar et al (2023).

The typology used aggregates CORINE Land Cover classes into broader ecosystem types. The definitions of *sparsely vegetated ecosystems* and *coastal beaches, dunes and wetlands* were refined following Babi Almenar et al. (2023), where the interested reader can find more precise specifications.

Ecosystem extent accounts provide the fundamental spatial baseline for assessing habitat availability and land cover change, key processes in land degradation and biodiversity loss. By tracking the proportional shares of natural and semi-natural ecosystems (e.g., forests and woodlands, grasslands) versus artificial surfaces (like Settlements and Other Artificial Areas - SOAS) within urban boundaries, these accounts directly quantify the habitat matrix and landscape heterogeneity of urban ecosystems. This is a prerequisite for understanding biodiversity potential and monitoring the effects of land-use change.

### 2.3. Urban ecosystem condition accounts

Ecosystem condition accounts compile and aggregate data on biophysical characteristics of ecosystem assets, providing insights into the ecological condition of different ecosystem assets and types within a defined accounting area and year (United Nations, 2021). These characteristics are quantified through metrics, often referred to as parameter-proxies, which can also indicate how far a characteristic deviates from a reference (optimal) condition. The SEEA-EA defined an ecosystem condition typology that organizes these characteristics into three groups (and six associated classes): abiotic ecosystem characteristics (physical state, chemical state), biotic ecosystem characteristics (compositional state, structural state, functional state), and landscape level characteristics (landscape and seascape characteristics) (United Nations, 2021). For further details on this typology and the criteria for selecting suitable condition variables within each group and class, please see Czúcz et al. (2021) and Keith et al. (2020).

Building on the SEEA-EA typology, the EU-wide Methodology (Vallecillo et al., 2022) identified relevant ecosystem condition variables for several ecosystem types for which EU level data were available. Among these ecosystems, a set of relevant condition variables for urban ecosystems was defined. Here we develop ecosystem condition accounts for four of those variables: i. share of imperviousness (physical state class); ii. annual average air concentration of PM_10_ (particulate matter, chemical state class); iii. share of green spaces (structural state class); and iv) share of tree canopy cover (structural state class). The values for all the variables were derived from publicly available Earth observation data, specifically from the Copernicus program, which also provides coverage for the United Kingdom and the Western Balkans. Detailed descriptions of each variable, their data sources, and spatial and temporal coverage are presented in Supplementary Material (SM) 1.

These four condition variables were selected not only for their environmental and policy relevance, but because they serve as direct indicators of urban ecological degradation, with consequent impact on ecological function. Imperviousness is a primary measure of land degradation (habitat loss) and of soil sealing. PM_10_ concentration reflects air chemical pollution causing stress to ecosystems. On the contrary, green space and tree canopy cover are critical structural components that support urban biodiversity, preserve natural habitats, and mitigate urban challenges like the urban heat island effect. These variables align with the need for indicators to monitor the effects of land degradation on biodiversity, as well as on human well-being. Three of the variables (PM_10_, share of green spaces, and share of tree canopy cover) correspond to variables and targets that must be reported for urban ecosystems under the amendment to the Regulation on Environmental-Economic Accounts and/or the NRR (European Parliament, Council of the European Union, 2024a, 2024b). Although not specifically listed for urban ecosystems, imperviousness is considered a relevant condition variable in other ecosystem accounts (European Parliament, Council of the European Union, 2024b). In addition, imperviousness, together with tree canopy cover and green space, directly influences local climate regulation in urban areas, an ecosystem service which monitoring is also required under the amended Regulation on Environmental-Economic Accounts (European Parliament, Council of the European Union, 2024b).

### 2.4. Urban ecosystem service flow account: air filtration

Ecosystem service accounts record data on final ecosystem service flows provided by ecosystem assets, i.e., supply tables, and inform on the use of those services by economic sectors, i.e., use tables (United Nations, 2021). These flows are reported in biophysical and monetary terms, forming different but interconnected accounting tables. Monetary ecosystem service flow accounts are derived by applying exchange value-based monetary valuation to biophysical ecosystem service flows (United Nations, 2021).

This study develops biophysical ecosystem service accounts for air filtration by vegetation, using the dry deposition of PM_10_ onto vegetation as a proxy. Deposition is modelled making use of the electrical resistance analogy, a methodological approach previously applied in studies of air filtration (Manes et al., 2016; Marando et al., 2016). In this analogy, the components of a plant are modelled as resistors in an electrical circuit. Air filtration was selected because of its policy relevance, as it is included among the mandatory ecosystem service accounts required by the amendment to the Regulation on Environmental-Economic Accounts (European Parliament, Council of the European Union, 2024b).

The calculation of PM_10_ dry deposition requires as inputs the time series of leaf area index (LAI) and of air concentration of PM_10_ during a recording year. PM_10_ concentration data were obtained from the Copernicus Atmosphere Monitoring Service at an hourly temporal resolution with spatial resolution of 0.1° for continental Europe, which is around 7.9 kilometers in Northen Europe. LAI data were taken from the Copernicus Global Land Service, a temporal resolution of 10 days and spatial resolution of 300 meters, and combined with LAI data from the MODIS Terra and Aqua datasets, at a temporal resolution of 4 days and spatial resolution of 500 meters.

As reported in (Babí Almenar et al., 2023), the calculation of PM_10_ dry deposition followed five steps:

- Aggregation of LAI data. LAI values were aggregated to monthly intervals, with missing values in the Copernicus Global Land Service dataset corrected using MODIS data;
- Calculation of deposition velocity 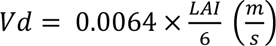 where 0.0064 m/s represents the average deposition velocity for PM_10_ at a reference LAI value of 6 (Hirabayashi et al., 2022; Manes et al., 2016);
- Calculation of PM_10_ flux: *F* = *Vd* × [*PM*10] × *timestep* (*Mass Unit*/*m*2) with time measured in seconds at one-hour time steps (3600 seconds).
- Calculation of *PM*_10_ deposition: *PM*_10_ *Dep* = *F* × *Ecosystem Asset Cell* (*Mass Unit*)
- Aggregation per reporting unit. PM_10_ deposition values were aggregated to the level of urban LAUs and expressed in tonnes.

The outcome is the total deposition of PM_10_ per reporting unit over the entire ecosystem accounting area during a recording year. However, to facilitate fair comparisons across reporting units of varying size and pollution levels, dry deposition is also expressed in terms of an efficiency metric. This metric is calculated by normalizing deposition values by the land area of each reporting unit and by their average annual PM_10_ concentration.

### 2.5. Representative municipalities of the Balkan Countries and Italy

As part of the CROSS-REIS project, one representative municipality from each Western Balkan country within the ecosystem accounting area was selected as a pilot city. In particular, the pilot municipalities are Kranj (SI), Rijeka (HR), Niš (RS), Podgorica (ME), and Tirana (AL), (Figure 1). For these municipalities, detailed insights into ecosystem extent, condition, and service accounts were gathered. They were chosen because collaborative urban greening initiatives, organized as Living Labs, are being planned within the CROSS-REIS project in each location. These initiatives bring together research institutions, local authorities, and citizens. By providing baseline information on the state of urban ecosystems, the analysis aims to support the design of specific urban greening actions.

Within the National Biodiversity Future Centre project, particularly Spoke 5, a set of six pilot municipalities in Italy (Torino (TO), Milano (MI), Firenze (FI), Roma (RO), Napoli (NA) and Campobasso (CA)) was selected. In these cities, Living Labs and various other activities are also being conducted, focusing on monitoring and planning urban green spaces and biodiversity. These efforts aim to enhance urban biodiversity and, by extension, improve urban ecological health.

Given their geographical proximity and the similarity of some objectives of both projects, comparing the Balkan pilot municipalities with the Italian was deemed valuable. This comparison serves to enrich the discussion on the potential utility of thematic urban ecosystem accounts for guiding urban greening initiatives and policies defined at a local governance level.

## 3. Results

### 3.1. Extent and composition of urban ecosystem accounts

National urban ecosystem accounting areas, comprising only cities and towns and suburbs, show high variability in their share of total land area (see the proportions of darker colors in the histogram of Fig. 2a) across the Western Balkans (from **∼**20% of HR to **∼**70% of ME), EU, EEA countries, and the UK, but far greater consistency in their share of national population despite differences of up to a 50% both in CR and in EU (Fig. 2b). For example, Iceland’s urban ecosystem area covers <2% of its land, yet it houses **∼**80% of its inhabitants. Within the Western Balkans, there is also considerable variability in the land area share of national accounting areas, and more homogeneous values for the population share covered. On average (see the rightmost bins of the histograms in Fig. 2), the Western Balkans (BLK) and CROSS-REIS (CR) country groups have a larger share of their territory within urban ecosystem areas than the EU-27/EEA/UK group (EUR), despite all three groups containing **∼**75% of their respective populations. Therefore, a common pattern emerges across the BLK, CR, and EUR groups and their constituent countries: the high variability in the land area covered is significantly reduced when measured by population share.

**Figure 2.**
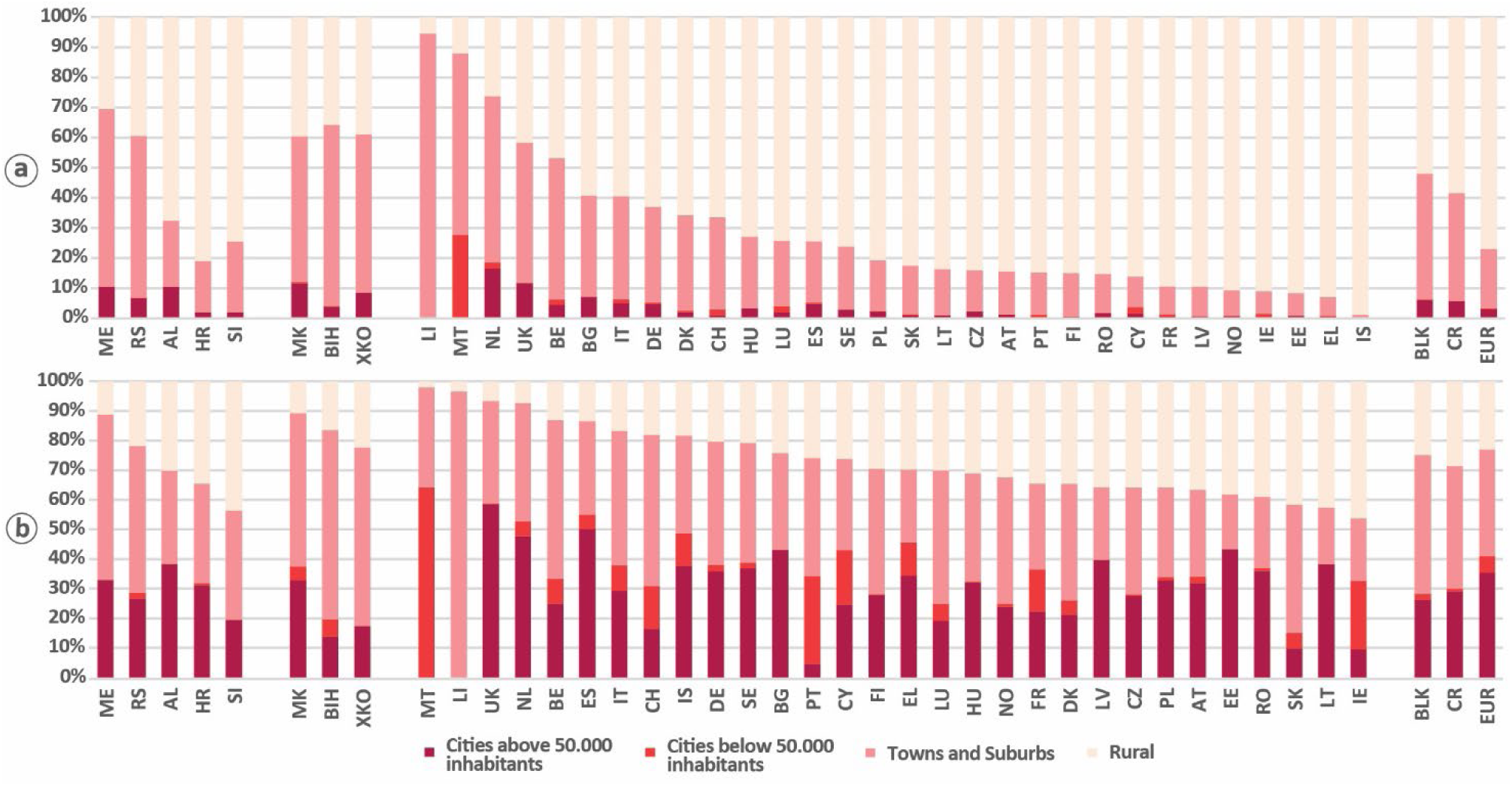
**Distribution of local administrative units (LAUs) across the urban-rural gradient. LAUs are classified as cities (above/below 50,000 inhabitants), towns and suburbs, or rural. National urban ecosystem accounting areas are composed only of LAUs classified as cities and towns/suburbs. (a) Share of total national land area in each class. (b) Share of total national population in each class.** From left to right, countries are grouped as: CROSS-REIS countries, other Western Balkan countries, and other European countries (the EU—excluding Croatia and Slovenia—EFTA, and the UK). The rightmost bars show weighted averages for the Western Balkans (BLK), CROSS-REIS countries (CR), and other European countries (EUR). European countries are ordered from most to least urban; Western Balkan countries and regional averages follow a fixed order. The full names for each country acronym are provided in the glossary.

The composition of accounting areas by urban class (cities above and below 50.000 inhabitants and towns and suburbs) also reveals an interesting, distinct land-population contrast. In terms of land area, towns and suburbs (pale pink portions of bins in Fig. 2a) form the largest and most homogeneous urban class within urban ecosystem accounting areas (darker pink portions). Spatially, cities (darkest pink) account for <5% of national land in most EUR countries, with notable exceptions (e.g., NL, BG, UK). Several BLK countries (e.g., ME, AL, MK) show higher shares, around 10%. For population share, however, the distribution across classes is more varied than for land share (Fig. 2b). While the share of total population within cities below 50.000 inhabitants is often low, denoting a concentration of people in large cities, the share in large cities varies more across countries, even within BLK. Nevertheless, the aggregated population shares for the BLK, CR, and EUR groups are quite similar, with EUR’s average total population share in cities being only about 10 percentage points higher than in BLK and CR. Detailed country-level land and population shares corresponding to these distributions are reported in SM 2.

### 3.2. National-level comparison of urban ecosystem accounts

National average values of urban ecosystem accounting areas for ecosystem extent, condition, and air filtration are shown for BLK, CR and EUR groups and their countries in Fig. 3-5. Detailed national-level ecosystem accounts for extent, condition and air filtration are also reported in SM 3.

**Figure 3.**
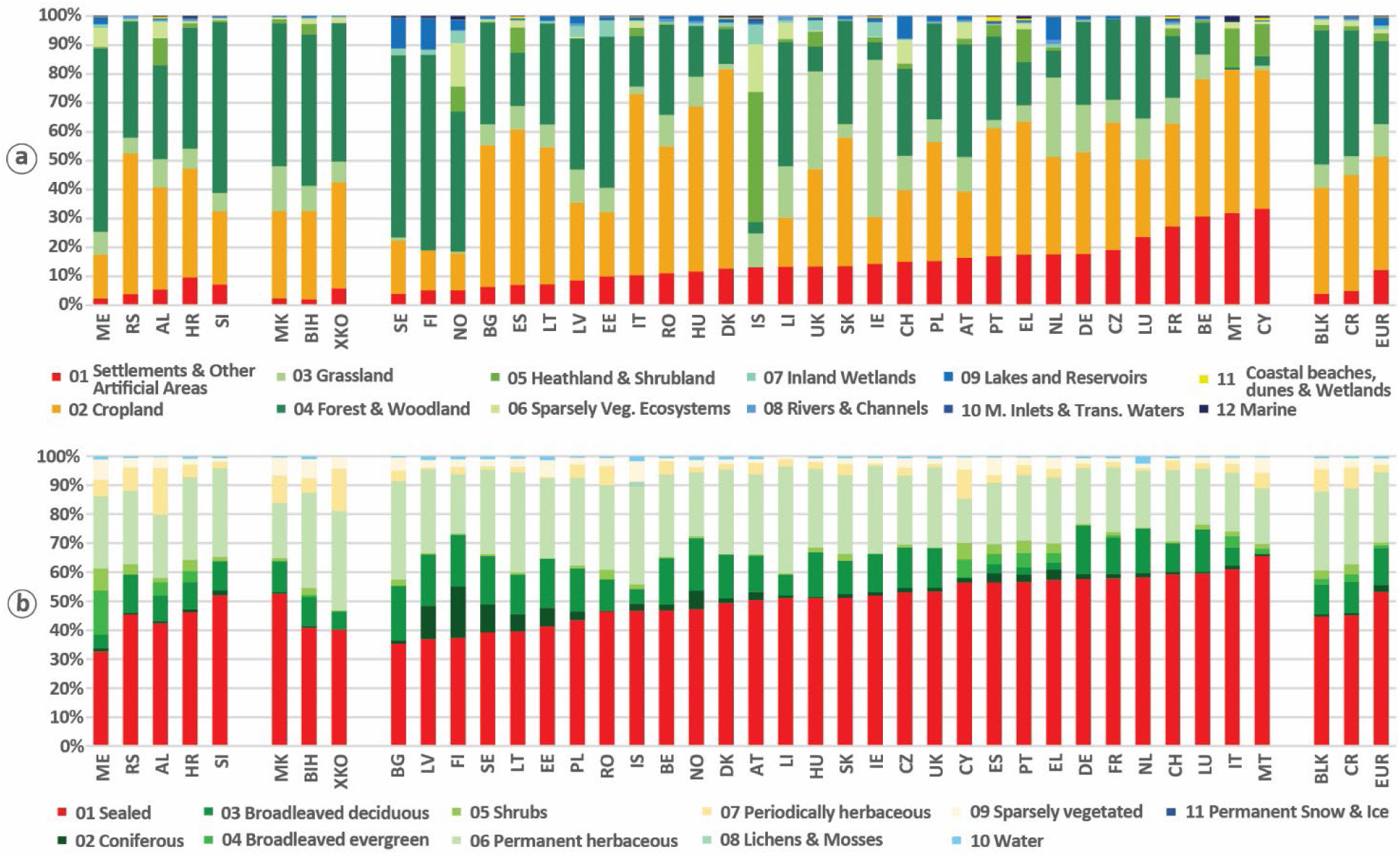
**Benchmarking of ecosystem extent in urban ecosystems of Western Balkan countries against other European countries. (a) Share of specific ecosystem types within urban ecosystems (i.e., urban local administrative units), and (b) Share of detailed land cover classes in *Settlements and Other Artificial Areas* within urban ecosystems (the red portion of each bin in panel a).** From left to right, countries are grouped as: CROSS-REIS countries, other Western Balkan countries, and other European countries (the EU—excluding Croatia and Slovenia—EFTA, and the UK). The rightmost bars show weighted averages for the Western Balkans (BLK), CROSS-REIS countries (CR), and other European countries (EUR). European countries are ordered from most to least urban; Western Balkan countries and regional averages follow a fixed order. The full names for each country acronym are provided in the glossary.

Regarding ecosystem extent (Fig. 3a), urban ecosystem composition in the Western Balkans is quite homogeneous, dominated by a few Ecosystem Types Level 1: Forests and Woodlands (40–60%), Croplands (**∼**30%), SOAS (below or **∼**5%), and Grasslands (**∼**8%). National values cluster closely, with few exceptions. For instance, Croatia (HR) has a higher SOAS share (**∼**10%), while Montenegro (ME) and Serbia (RS) are outliers for Croplands (around 15% and 48%, respectively). This regional homogeneity contrasts with high variability among other European countries, where the SOAS share alone ranges from 3% to 30%, a pattern of high variation that extends to all other ecosystem types. This result reflects in part the geographic-climatological and the economic-developmental differences between countries. Despite this national-level contrast, the aggregated regional values for the BLK, CR, and EUR groups are more similar. The key differences are a higher SOAS share in EUR (**∼**12%, three times larger than BLK/CR) and a correspondingly lower Forests and Woodlands share in EUR, approximately 29%, close to a 15 percentage points lower than in CR and BLK.

A detailed view within the SOAS in urban ecosystems reveals greater similarity between individual Western Balkan and other European countries (Fig. 3b). Simply said, SOAS are made by very similar fractions of land covers almost everywhere in geographical Europe. For sealed land cover, most countries fall within 40–60%, with the Western Balkans values typically being near 40% (exceptions: SI and MK **∼**50%; ME **∼**30%). Homogeneity is also seen in permanent herbaceous covers (20–30%) and broadleaved deciduous forests (10–15%), though some Nordic/Baltic countries (e.g., FI, SE, NO) and Bulgaria (BG) show forest shares slightly above 15%. At the aggregated level, BLK, CR, and EUR values remain similar, with EUR’s sealed cover share being just an 8 percentage points higher, compensated by its lower permanent and periodically herbaceous covers. This indicates a typical SOAS composition in urban ecosystems for most European countries within and outside the Western Balkans.

In terms of ecosystem condition (Fig. 4), values for imperviousness share (panel a), urban green share (b), tree cover share (c), and PM_10_ concentration (d) are relatively homogeneous across Western Balkan countries but highly variable across other European nations. However, aggregated values for the BLK, CR, and EUR groups are similar, with **∼**10% difference for imperviousness and tree cover, and **∼**15% for urban green.

**Figure 4.**
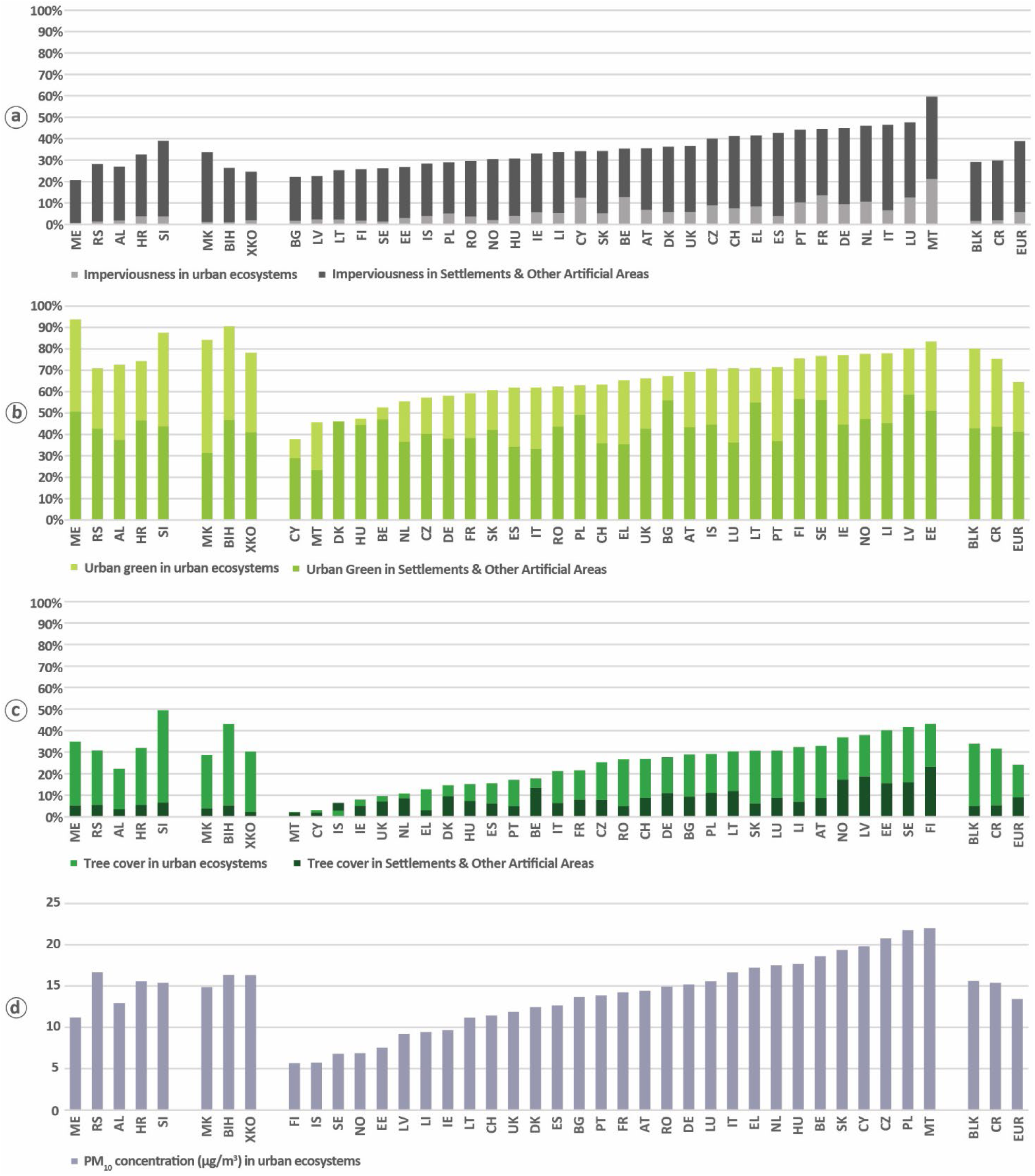
**Benchmarking of ecosystem condition in urban ecosystems of Western Balkan countries against other European countries. (a) Imperviousness share, (b) Urban green share, (c) Tree cover share, and (d) PM₁₀ concentration.** Values are shown for the urban ecosystem accounting area of each country (i.e., urban local administrative units) and for the *Settlements and Other Artificial Areas* within urban ecosystems. Bars for *Settlements and Other Artificial Areas and* for the entire urban ecosystems overlap but values are not cumulative and should be interpreted as absolute. From left to right, countries are grouped as: CROSS-REIS countries, other Western Balkan countries, and other European countries (the EU—excluding Croatia and Slovenia—EFTA, and the UK). The rightmost bars show weighted averages for the Western Balkans (BLK), CROSS-REIS countries (CR), and other European countries (EUR). European countries are ordered from most to least urban; Western Balkan countries and regional averages follow a fixed order. The full names for each country acronym are provided in the glossary.

For imperviousness (Fig. 4a), national averages in the Western Balkans are systematically low for total urban ecosystems (<5%) and around 20–30% for SOAS, except for Slovenia (**∼**40%). In contrast, other European countries show a wide range (20–60%), with many exceeding 30% for SOAS.

Urban green shares (Fig. 4b) are around 70–80% of urban ecosystems in most Western Balkan countries, and 40–50% within their SOAS. Two of the CR countries (ME and SI) and the other three Western Balkan countries (MK, BIH, XKO) have urban green shares higher than the highest value for EUR countries (that of EE). Our analysis thus reveals that urban settlements in the Western Balkan region are in the top 5 greenest cities/towns in geographical Europe. Other European countries exhibit lower and more variable shares overall, though SOAS values are more convergent, leading also to similar aggregated values for BLK, CR and EUR (**∼**40%).

Tree cover share (Fig. 4c) is the most variable condition metric in the Western Balkans and other European countries. Within SOAS, values for the Western Balkan countries are consistently low (≤5%), placing the BLK and CR with the lower range of European values. In contrast, the values of tree cover share in urban ecosystem are consistently high and on average higher than the average EUR country. As a result, the top two values of tree cover share within geographical Europe are registered in the Western Balkans, with Slovenia and Bosnia Herzegovina exceeding the top EUR country (Finland). Consequently, the aggregated tree cover share within SOAS is **∼**5% higher in EUR than in BLK or CR.

PM_10_ concentrations (Fig. 4d) average around 15 µg/m³ in most Western Balkan countries. Only about one-third of other European countries exceed this level, but higher values elsewhere compensate, resulting in only slightly elevated (by **∼**5 µg/m³) averages for BLK and CR compared to EUR. This is saying that, despite the good level of greening in the Western Balkans, the pollution generated by human activities remains high and far from being compensated by air filtration by vegetation.

Overall, the results shown in Fig. 4 reveal that urban condition values within Western Balkans are quite homogeneous, as per ecosystem extent values. This homogeneity is extended to condition variables within SOAS, not only just within the Western Balkans but across most of the other European countries.

Air filtration services provided by plants, measured as PM_10_ deposition, varies strongly across countries, both in terms of the share of the absolute deposition per land unit deposited in SOAS (g/m²), and of the deposition normalized by PM_10_ concentration (µg/m³) (Fig. 5). For Western Balkan countries, most PM_10_ deposition occurs outside SOAS (Fig. 5a), consistent with SOAS’s low share of urban ecosystem area in the region (see again Fig. 3a). Furthermore, the deposition share in SOAS is lower than its area share, indicating a lower per-unit deposition efficiency. This pattern is common to most European countries and may be explained by the lower amount of vegetation, compared to other ecosystem types, that drives the supply of this service. In terms of absolute deposition per land unit (Fig. 5b), Western Balkan countries perform better than most of Europe, both for total urban ecosystems and for SOAS. CROSS-REIS countries typically show the highest values. The normalized values (Fig. 5c) reveal that these high deposition rates in CR countries are not just an artifact of high PM10 concentrations but reflect higher effective leaf area, as their normalized deposition remains elevated. Despite high national-level heterogeneity, especially in European countries outside the Western Balkans, the aggregated BLK, CR, and EUR values for PM_10_ deposition, at least in SOAS, are notably similar.

**Figure 5.**
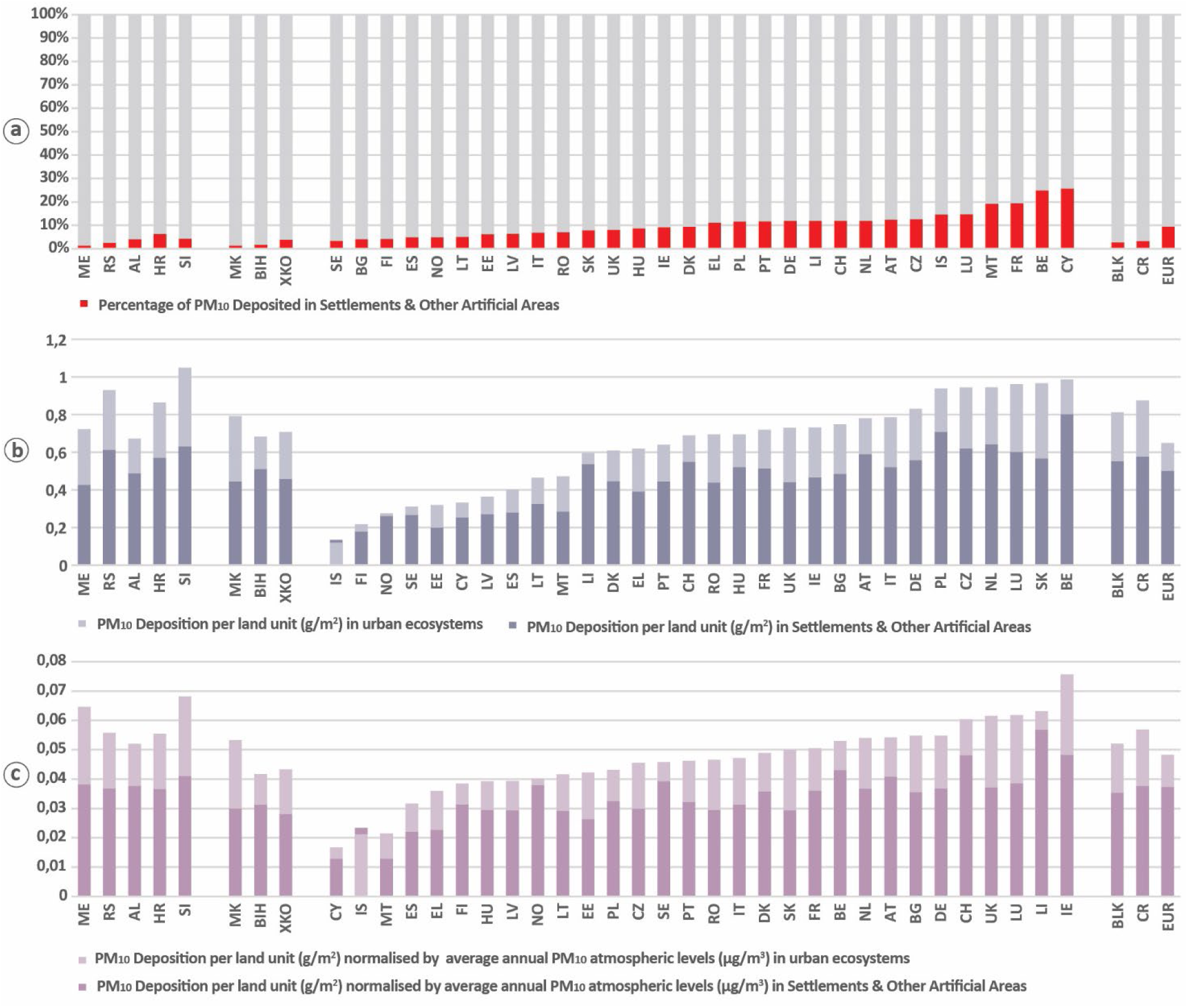
**Benchmarking of the ecosystem service air filtration (PM_10_ deposition) in urban ecosystems of Western Balkan countries against other European countries. (a) Share of PM_10_ deposited in *Settlements and Other Artificial Areas* of urban ecosystems. (b) PM_10_ deposition per land unit (g/m^2^) in *Settlements and Other Artificial Areas* of urban ecosystems and in the entire urban ecosystems. (c) PM_10_ deposition per land unit (g/m^2^) normalized by average annual atmospheric PM_10_ levels (µg/m^3^) in *Settlements and Other Artificial Areas* of urban ecosystems and in the entire urban ecosystems.** Bars for *Settlements and Other Artificial Areas and* for the entire urban ecosystems overlap but values are not cumulative and should be interpreted as absolute. From left to right, countries are grouped as: CROSS-REIS countries, other Western Balkan countries, and other European countries (the EU—excluding Croatia and Slovenia—EFTA, and the UK). The rightmost bars show weighted averages for the Western Balkans (BLK), CROSS-REIS countries (CR), and other European countries (EUR). European countries are ordered from most to least urban; Western Balkan countries and regional averages follow a fixed order. The full names for each country acronym are provided in the glossary.

### 3.3. Influence of geography, climate, and urban classes on urban ecosystem account values

Spatial and statistical visualizations of ecosystem condition and air filtration service values (Fig. 6-7) reveal that variation is partly associated with geographical and climatic factors, but not with urban classes. These visualizations identify potential relationships not apparent from national-level comparisons. They show correlations rather than causality, while still highlighting relevant factors when defining and applying urban ecosystem accounts, as elaborated in the Discussion.

**Figure 6.**
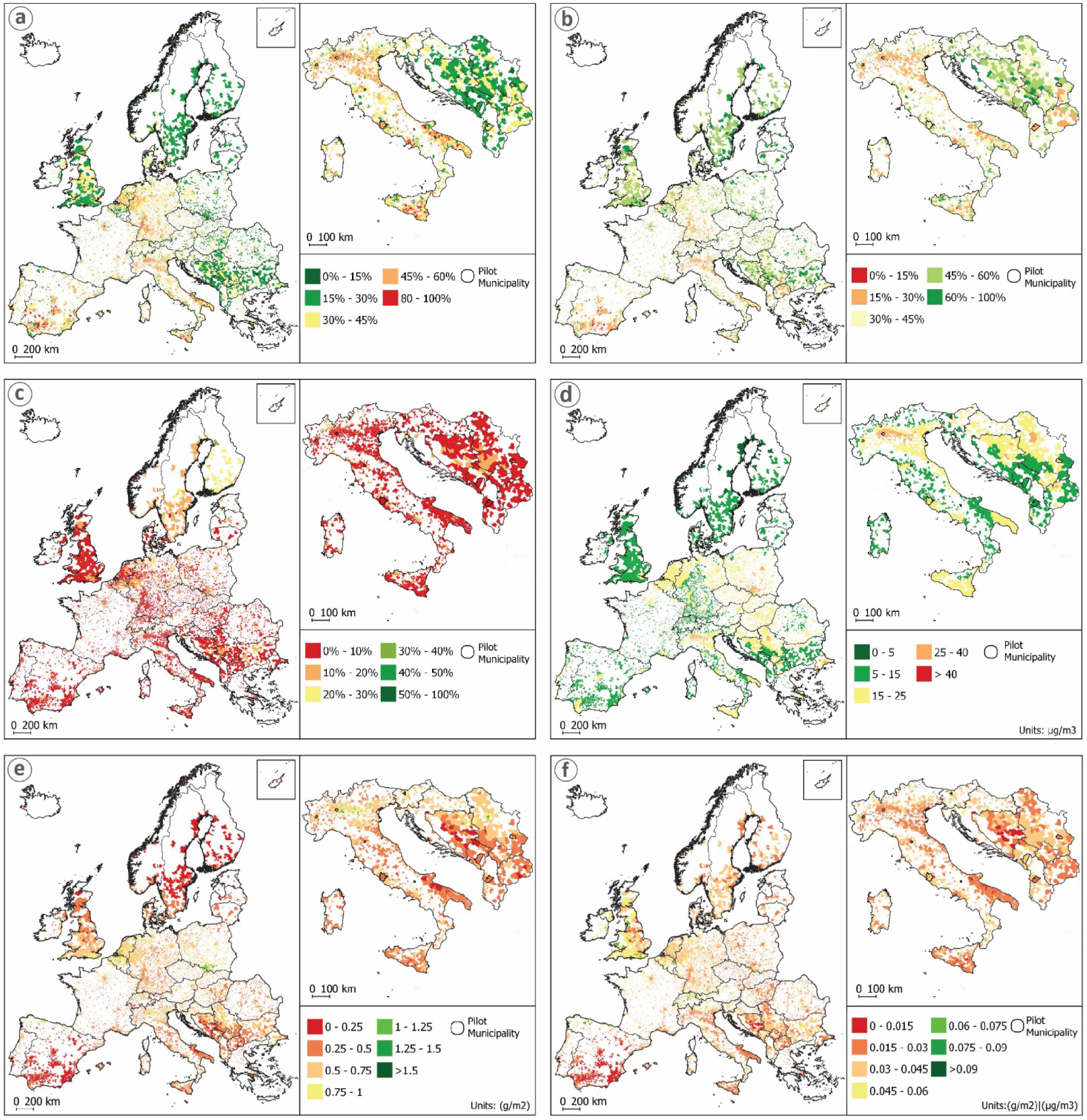
**Mapping of ecosystem condition and the ecosystem service air filtration in the *Settlements and Other Artificial Areas* of urban ecosystems in Europe, with a zoom-in on the Western Balkans and Italy. (a) Imperviousness share, (b) Urban green share, (c) Tree cover share, (d) PM₁₀ concentration (µg/m^3^), (e) PM_10_ deposition per land unit (g/m^2^), (f) PM_10_ deposition per land unit (g/m^2^) normalized by average annual atmospheric PM_10_ levels (µg/m^3^).** Values are presented per urban ecosystem (i.e., urban local administrative unit). In the Supplementary Material 4 each map is visualized at a larger scale, and corresponding maps for overall values for each urban ecosystem rather than only for its Settlements and Other Artificial Areas are also included.

**Figure 7.**
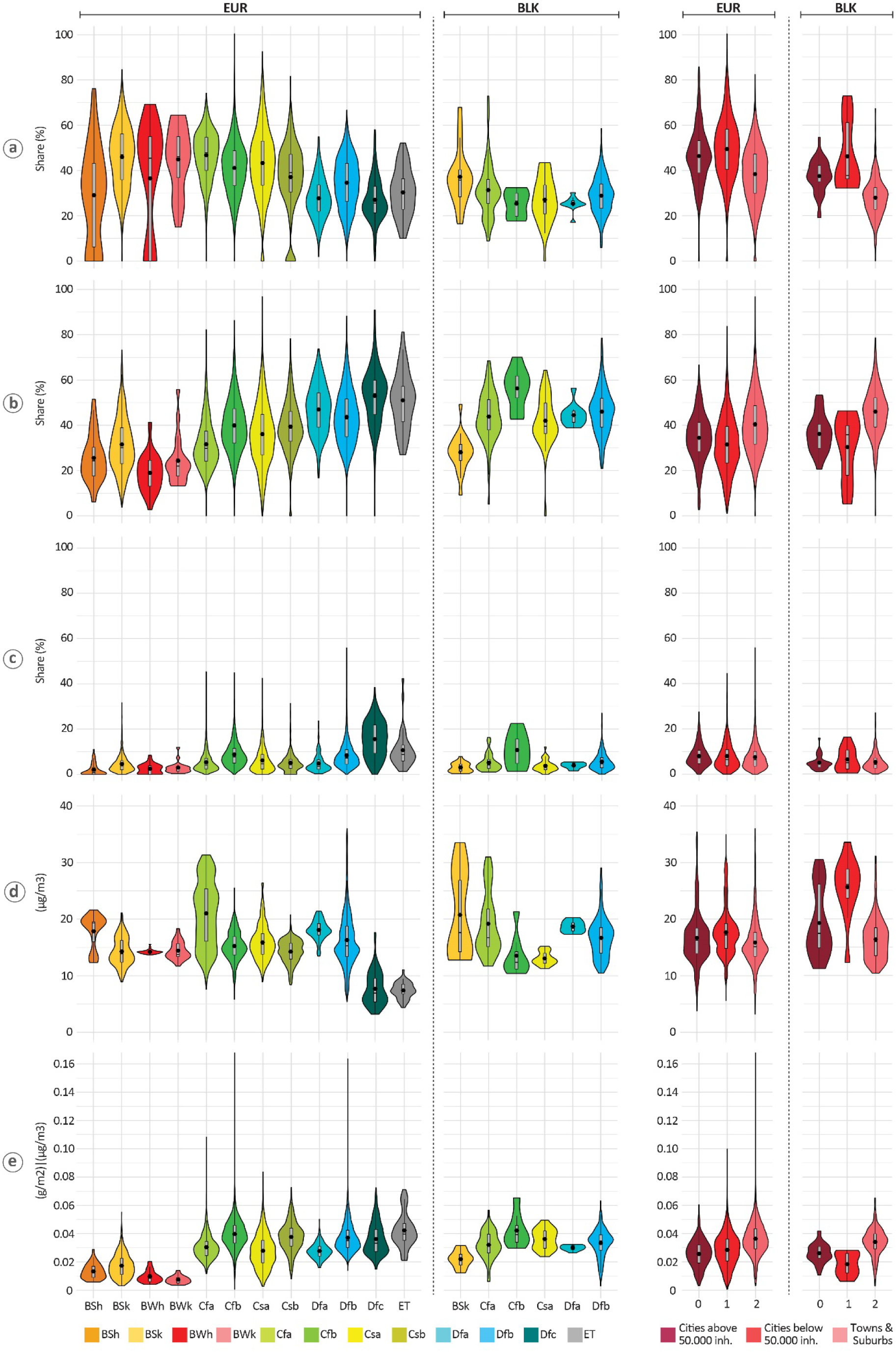
**Statistical distribution of ecosystem condition (a to c) and the ecosystem service air filtration values (d to e) in the *Settlements and Other Artificial Areas* of urban ecosystems, aggregated by Köppen-Geigger climate classes (left) and urban classes (right). Violin plots show values for all European countries (EUR) and the Western Balkans (BLK): (a) Imperviousness share, (b) Urban green share, (c) Tree cover share, (d) PM₁₀ concentration (µg/m³), and (e) PM_10_ deposition per land unit (g/m^2^) normalized by average annual atmospheric PM_10_ levels (µg/m^3^).** Supplementary Material 5 provides values for each urban ecosystem rather than only for its Settlements and Other Artificial Areas. The acronyms for the Köppen-Geigger climate class are in the Glossary.

For instance, imperviousness in SOAS (Fig. 6a) is notably lower in urban ecosystems of the Nordic, Baltic, Eastern-Central European, and most Western Balkan regions compared to Western and Southern Europe. Albania and North Macedonia display exceptionally lower values than the rest of Western Balkan countries. These geographical areas correspond well with the Dfb and Dfc climate classes, and therefore, their statistical distributions show also systematically lower values (Fig. 7a).

As expected from imperviousness values, the geographical pattern for urban green share is broadly inverse, though less consistent (Fig. 6b). This is also reflected in generally higher distributions for the Dfb and Dfc zones (Fig. 7b). Notably, within the Western Balkans, areas outside these climate zones also maintain high urban green shares. In contrast, the spatial distribution of tree cover in SOAS is distinct (Fig. 6c), with the highest shares (10–30%) concentrated in the Nordic countries (Norway, Sweden, and Finland), while most of the rest of Europe exhibits values below 10%. This pattern is also evident when tree cover is examined by climate class: the Dfc class shows consistently higher values than all others, with distributions from the lower quartile upward exceeding those of other classes. A similar pattern is visible for PM_10_ concentration, where Dfc and ET climate classes show much lower values across their entire distributions (Fig. 7d), though this is not clearly visible spatially (Fig. 6d). For air filtration (normalized by PM_10_ deposition), distributions clearly show that all arid climate classes (BSh, BSk, BWh, BWk) have systematically lower values, especially the desert classes (BWh, BWk) (Fig. 7e). This is explained by their lower urban green and tree cover, implying less annual leaf area. This latter pattern is not easily discernible from the spatial visualization.

Despite the overall lack of clear patterns related to urban classes, an exception exists within the Western Balkans. There, LAUs classified as towns and suburbs have clearly lower imperviousness and higher urban green shares, which may also explain their higher air filtration values (Fig. 7a-b, e).

### 3.4. Local-level comparison of urban ecosystem accounts across pilot cities

Local-level (i.e., individual LAU) values of ecosystem extent (Fig. 8), condition (Fig. 9), and air filtration (Fig. 10) are computed and presented for pilot cities in CROSS-REIS countries and for NBFC pilot cities in Italy, and are compared with national- and regional-level averages (BLK, CR, EUR). Detailed local-level ecosystem accounts for extent, condition, and air filtration are provided in SM 6.

**Figure 8.**
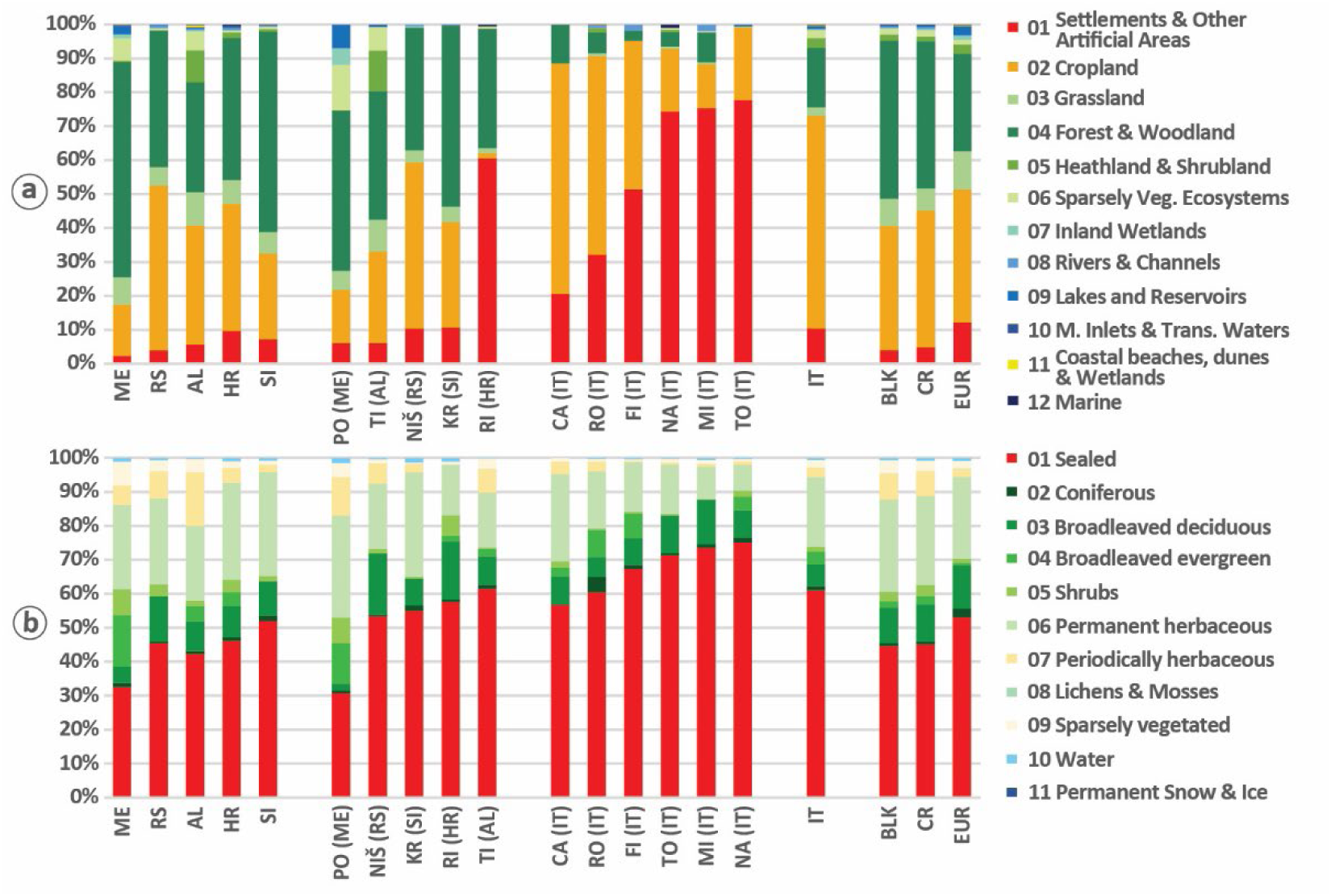
**Benchmarking of ecosystem extent in urban ecosystems of pilot municipalities* of CROSS-REIS countries and in NBFC pilot municipalities in Italy against average national and regional values. (a) Share of specific ecosystem types within urban ecosystems (i.e., urban local administrative units); (b) Share of detailed land cover classes in the ecosystem type *Settlements and Other Artificial Areas* within urban ecosystems (red in panel a).** In each graph, the group of bins computed for CROSS-REIS pilot municipalities are ordered from lowest to highest values and displayed to the right of the group of bins for their national corresponding reference value and to the left of the group of bins for the NBFC pilot cities in Italy, followed by the Italian average value. The regional averages for Western Balkans (BLK), CROSS-REIS countries (CR), and other European countries (EUR) are shown in a consistent fixed order. The full names for each country and municipality acronym are provided in the glossary.

**Figure 9.**
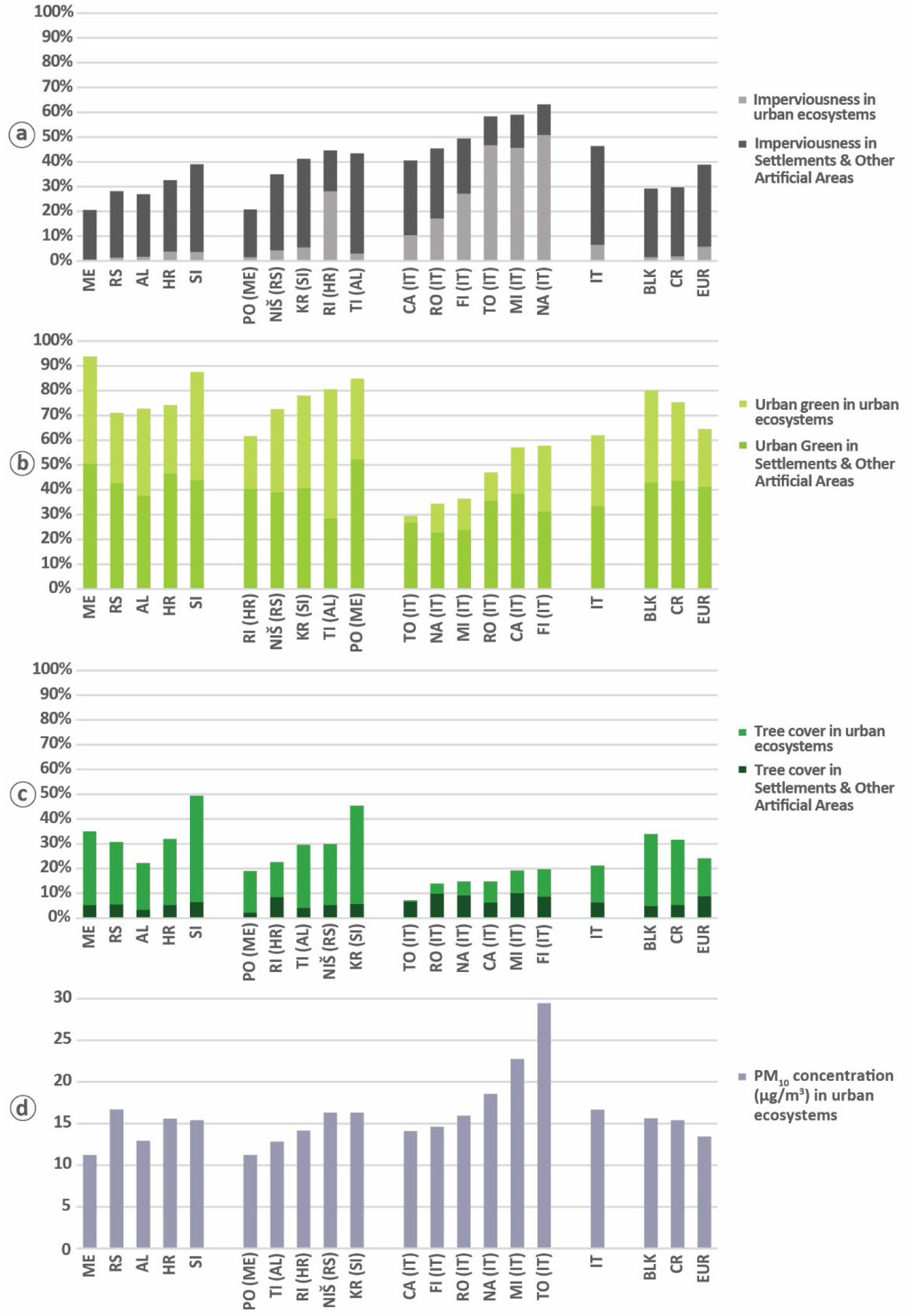
**Benchmarking of ecosystem condition in urban ecosystems of pilot municipalities* in CROSS-REIS countries and in NBFC pilot municipalities in in Italy against average national and regional values. (a) Imperviousness share, (b) Urban green share, (c) Tree cover share, and (d) PM₁₀ concentration.** Values are shown for the urban ecosystem accounting area and for the *Settlements and Other Artificial Areas* within urban ecosystems. Bars for *Settlements and Other Artificial Areas and* for the entire urban ecosystems overlap but values are not cumulative and should be interpreted as absolute. In each graph, the group of bins computed for CROSS-REIS pilot municipalities are ordered from lowest to highest values and displayed to the right of the group of bins for their national corresponding reference value and to the left of the group of bins for the NBFC pilot cities in Italy, followed by the Italian average value. The regional averages for Western Balkans (BLK), CROSS-REIS countries (CR), and other European countries (EUR) are shown in a consistent fixed order. The full names for each country and municipality acronym are provided in the glossary.

**Figure 10.**
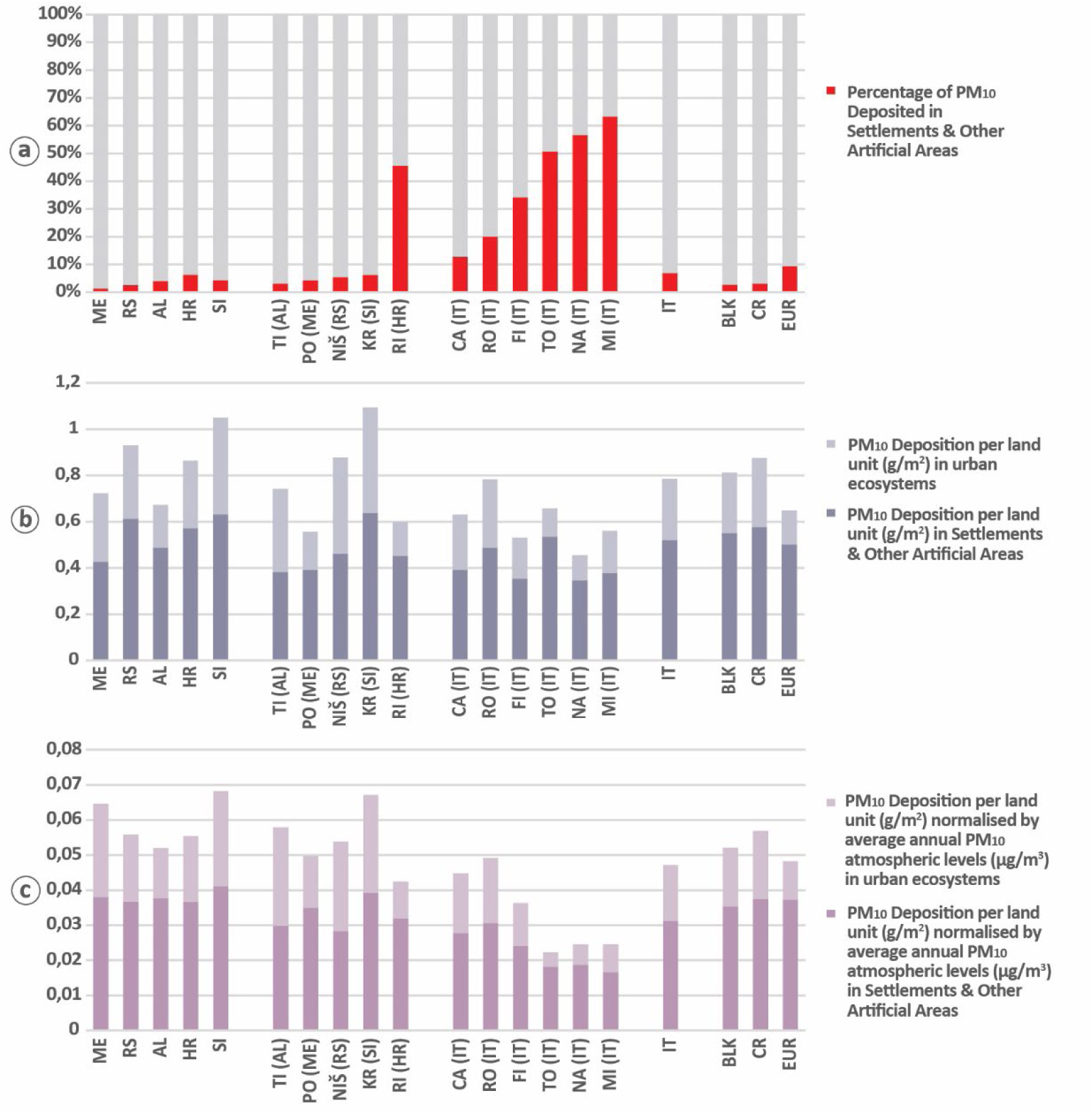
**Benchmarking of the ecosystem service air filtration (PM_10_ deposition) in urban ecosystems of pilot municipalities* in CROSS-REIS countries and in Italy against average national and regional values. (a) Share of PM_10_ deposited in *Settlements and Other Artificial Areas* of urban ecosystems. (b) PM_10_ deposition per land unit (g/m^2^) in *Settlements and Other Artificial Areas* of urban ecosystems and in the entire urban ecosystems. (c) PM_10_ deposition per land unit (g/m^2^) normalized by average annual atmospheric PM_10_ levels (µg/m^3^) in *Settlements and Other Artificial Areas* of urban ecosystems and in the entire urban ecosystems.** Values are shown for the urban ecosystem accounting area and for the *Settlements and Other Artificial Areas* within urban ecosystems. Bars for *Settlements and Other Artificial Areas and* for the entire urban ecosystems overlap but values are not cumulative and should be interpreted as absolute. In each graph, the group of bins computed for CROSS-REIS pilot municipalities are ordered from lowest to highest values and displayed to the right of the group of bins for their national corresponding reference value and to the left of the group of bins for the NBFC pilot cities in Italy, followed by the Italian average value. The regional averages for Western Balkans (BLK), CROSS-REIS countries (CR), and other European countries (EUR) are shown in a consistent fixed order. The full names for each country and municipality acronym are provided in the glossary.

In terms of ecosystem extent (Fig. 8a), CROSS-REIS pilot cities fit well the patterns observed at their corresponding national level. CROSS-REIS cities show high homogeneity in values, except for Rijeka, with values for dominant ecosystem types (Forests/Woodlands, Croplands, SOAS, Grasslands) closely matching their national and BLK/CR averages. In contrast, Italian pilot cities show high internal variability and frequently differ from the national Italian average, which is similar only to Campobasso. This aligns with Italy’s urban ecosystems being predominantly composed of towns and suburbs like Campobasso (see again Fig. 2a). Overall, Italian pilot cities have a larger share of SOAS and Croplands than CROSS-REIS cities, a difference not explained by urban class, suggesting other regional factors are at play, as both sets consist of comparable large cities and just one LAU classified as town and suburb.

When looking at extent within SOAS (Fig. 8b), values reveal greater similarity between Western Balkan and Italian cities. Both sets of pilot cities quite mirror their corresponding national patterns, but generally have higher shares of sealed cover,except for Podgorica and Campobasso, SOAS of Italian pilots clearly have higher shares of sealed cover than Western Balkan pilots. Despite differences in specific share values, the dominant land covers (i.e., sealed, broadleaved trees, permanent herbaceous) are nevertheless consistent across national and BLK/CR/EUR averages and pilot cities.

In the Western Balkans, ecosystem condition in pilot cities (Fig. 9) largely mirrors the patterns observed for ecosystem extent (Fig. 8a), showing values quite similar to national and BLK/CR averages. Although similarity is less pronounced for all variables, except for PM_10_ concentration (Fig. 9d). For imperviousness, Tirana and Rijeka are notable exceptions, with values much higher than their national averages, and similar to the values of Italian cities – which are systematically higher than those of their CR counterparts; in Rijeka, this is true for both SOAS and the total urban ecosystem (Fig. 9a). For urban green, Fig. 9b makes very clear that the CR pilot municipalities have significantly higher values than the Italian pilot cities. At a closer look, we can say that the overall values in most Western Balkan pilots are slightly below their national averages, though SOAS values align more closely. Tirana is an exception, with urban green in SOAS ∼10% lower than the national average (Fig. 9b). For tree cover (Fig. 9c), again the CROSS-REIS pilots overcome the NBFC pilots in a significant manner. Among Wester Balkan cities, relevant exceptions are Rijeka and Podgorica, which have lower total tree cover than their national averages. Interestingly, Rijeka’s tree cover in SOAS is higher than the Croatian national average for SOAS, and Tirana’s overall tree cover value is higher than its national average. An overall comparison between Italian and Western Balkan pilot sets reveals that Italian cities generally perform significantly worse across most condition metrics. They exhibit higher imperviousness and PM_10_ concentration, and lower urban green cover for both SOAS and total urban ecosystems. The sole exception is tree cover share within SOAS, where Italian values tend to be slightly higher.

Regarding air filtration service, as with national averages, most PM_10_ deposition occurs outside SOAS in pilot cities (Fig. 10a), which is consistent with the low areal share of SOAS in these ecosystems (see again Fig. 8a). In addition, the deposition share within SOAS is lower than its area share, indicating a lower per-unit deposition efficiency. This pattern also holds for Italian pilot cities and is reflected in national averages, for the same underlying reasons, i.e., a lower amount of vegetation compared to other ecosystem types. For absolute PM_10_ deposition per land unit (Fig. 10b), values vary across all pilots, though less within the Italian set. In contrast, deposition per unit within SOAS is more consistent across all cities, with values clustered in a range around 0.4–0.6 g/m². Normalized PM_10_ deposition, per unit of PM_10_ concentration, (Fig. 10c) shows that values within SOAS for Western Balkan pilots are very similar to, though often slightly below, their national averages. Furthermore, the Italian pilots frequently show lower normalized values than the Western Balkan set, both for total urban ecosystems and for their SOAS components. These normalized values indicate that pilots in the Western Balkans have a higher effective leaf area, which is consistent with their greater extent of non-artificial ecosystem types (Fig. 8a,b) and higher urban green cover (Fig. 9b).

## 4. Discussion

### 4.1. Interpreting the spatial patterns and disparities across urban ecosystems

The pilot urban ecosystem accounts in the Western Balkan region, specifically in the CROSS-REIS countries, reveal both contrasts and commonalities. At the aggregate European level, urban ecosystems in the Western Balkans are not ecological outliers; their key metrics for extent, condition, and services fall within the continental range. For instance, the average share of green space within urban settlements (SOAS) remains comparable (∼40-50%), despite cities in the Western Balkans have often a higher overall greenness than most other European countries. This regional coherence likely stems from shared biogeographic contexts, visible in the clustering of condition values by climate zone (see again Fig. 7) and the spatialization of outcomes (Fig.6). Common recent historical pathways, including post-socialist transitions and, for several countries, a legacy of Yugoslav-era spatial planning may also play a role in the homogeneity of outputs within the Western Balkans. One of the critical findings of our study is that Western Balkan urban ecosystems display characteristics that are on a continuum with the rest of Europe, validating the direct relevance and transferability of the SEEA-EA accounting framework for the region’s EU integration agenda.

Beneath this regional coherence, national and local drivers create distinct ecological profiles, which the accounts help to visibilize. These patterns can be broadly explained by understanding the interplay of policy frameworks, economic structure, and geography within each country and pilot municipality.

In Western Balkan countries with established planning systems and strong environmental governance, such as Slovenia and Croatia, which are already integrated into the EU, values of thematic urban ecosystem accounts nonetheless reflect a tension between regulated urban form and functional ecological deficits.

Slovenia’s urban ecosystems profile is dominated by extensive peri-urban forests, a result of extensive Natura 2000 protection areas - over 37% of national territory (Ministry of Natural Resources and Spatial Planning of Slovenia, 2024) - agricultural land abandonment, and conservative forest management (Čater and Železnik, 2021). The pilot municipality of Kranj is a national outlier and exhibits exceptionally high forest cover reinforced by topography, water protection zones, and strict spatial plans with strong emphasis on nature conservation (Mestna občina Kranj, 2014). Yet, as in national average values, its urban core is highly sealed, with limited tree canopy, thus weakening local ecological functions (e.g. shading and evapotranspiration) where most people need them. This is also reflected in air quality patterns: despite extensive surrounding green cover, PM₁₀ concentrations in Kranj are strongly influenced by household wood heating, traffic emissions, basin topography, and limited atmospheric dispersion, resulting in persistently high concentration levels.

Croatia’s hierarchically coordinated spatial planning has constrained sprawl and maintained low overall imperviousness at the city scale (<10% of urban LAU area), yet topographic constraints along the Adriatic coast concentrate development into dense and highly sealed urban cores (imperviousness >30% in SOAS). Indeed, Rijeka in Croatia exemplifies the compact, port-city morphology that leads to high imperviousness in its core (∼40% in SOAS) despite the strategic greening ambitions of the municipality. Rijeka, high imperviousness is a direct function of its constrained geography, a narrow coastal strip between the sea and mountains, and its industrial port legacy. While deindustrialization has reduced heavy industrial emissions, traffic and maritime activity sustain moderate PM_10_ levels (Alebić-Juretić and Mifka, 2022), illustrating how compact urban form, geography, and economic restructuring jointly condition local ecological outcomes despite increasingly ambitious greening policies.

Conversely, in contexts of rapid transformation, such as in Albania or in Serbia, the values of the thematic urban ecosystem accounts highlight development pressures and data gaps.

In Albania, rapid urbanization driven by migration toward major cities, combined with a planning tradition focused on construction and weak enforcement of forestry and air-quality policies (Aliaj and Rossi, 2015; Instituti i Statistikave, 2024), have led to intense soil sealing urban cores and declining settlement-level green cover, as condition values showcased. The 2015 territorial reform substantially enlarged municipalities by amalgamating surrounding rural areas into local administrative units. As a result, ecosystem condition values are inflated when interpreted for the entire urban ecosystem area rather than for SOAS alone, thereby masking intensive soil sealing and sparse tree cover in urban cores. This effect is clearly illustrated by the local-level results for Tirana, which appear to reflect a highly green peri-urban area but in fact represent enlarged LAUs that may not accurately capture a real urban ecosystem. Although national strategies increasingly acknowledge green infrastructure and air-quality integration, implementation remains uneven, while construction activity, transport, biomass burning, and seasonal wildfires continue to drive PM_10_ pollution.

Serbia presents a structurally different but equally challenging urban ecosystem profile. Despite a land base dominated by cropland and forests, as showcased in Fig. 3a, historical spatial planning and urban development strategies have under-prioritized urban greenery (Malinić et al., 2025), resulting in one of the lowest shares of green urban space among the CROSS-REIS countries (as illustrated in Fig 4b and 9b). At the same time, Serbia records moderate-severe PM_10_ concentrations, which is explained by coal-based energy production, ageing transportation fleets, industrial and agricultural emissions, and topographic confinement in basins like the Pannonian Plain (North of Serbia) and the Niš valley (in the South, and more locally around Niš). While recent drafts of spatial and climate policies explicitly recognize the loss and fragmentation of urban green spaces and promote green and blue infrastructure delayed implementation and persistent energy and mobility structures continue to constrain improvements in urban ecological performance (Matić, et al., 2025).

Montenegro presents a case where proactive policy alignment and favorable natural assets coincide, factors that result in comparatively high urban green shares and lower PM_10_ levels (as illustrated in Fig. 4b,d). The country’s early constitutional designation as an “ecological state” (Parliament of Montenegro, 2007), combined with early adoption of EU environmental *acquis* has supported stronger frameworks for nature protection, air quality management, and environmental spatial planning. The lack of heavy industry, limited reliance on polluting energy sources, and slower urbanization further reduce pressure on urban ecosystems. The capital, Podgorica, benefits from this orientation and its riverside location, though sustaining these conditions will require robust planning as development pressures increase. However, much of Montenegro’s favorable performance still reflects structural conditions—abundant natural landscapes, compact cities, and restrained development—rather than fully institutionalized urban ecological management. As urbanization accelerates, particularly in Podgorica and coastal areas, sustaining these outcomes will depend on the effective implementation and enforcement of emerging planning and environmental policies.

Ultimately, the accounts performed in this study underscore a fundamental lesson: the scale of analysis is critical. National aggregates and even city-wide (LAU) values can obscure the conditions where people live and ecosystems are most stressed—within the settled urban core (SOAS). This scale-dependency is not a methodological artefact but a crucial diagnostic insight. It confirms that effective policy must be informed by data at multiple scales, connecting national strategic frameworks with the granular, site-specific realities that determine ecological outcomes and human well-being in cities.

### 4.2. The role of urban ecosystem accounts to inform urban greening policy and actions

By providing a standardized and spatially explicit evidence base, thematic urban ecosystem accounts can help bridge the gap between high-level regulatory ambitions (such as those embedded in the EU NRR), and actionable, locally relevant planning and management decisions. Our study offers initial indications that thematic urban ecosystem accounts may constitute an easily transferable tool for operationalizing urban greening policy in the Western Balkans, and not just EU and EFTA countries where implementation is already partly in progress with the regulation on the amendment on environmental economic accounts (European Parliament, Council of the European Union, 2024b).

First, urban ecosystem accounts can support the establishment of quantitative baselines necessary for setting, calibrating, and tracking meaningful policy targets. Across Europe, including in Western Balkan countries, a major barrier to effective urban greening policy is the lack of harmonized spatial data against which ecosystem condition reference values, such as minimum green space provision or maximum soil sealing, can be defined and assessed. In Albania and Montenegro, where no formal accounts exist and environmental statistics remain fragmented, applying urban ecosystem accounting based on publicly and free available data would provide a first comprehensive picture of urban ecological conditions and support priority identification and target setting. For example, the 2018 accounts presented here give a simple foundational benchmark for future monitoring that can be used for both comparison of performance within the Western Balkans or/and across Europe. More importantly, the accounts can help to translate broad and somehow ideal policy concepts into specific, measurable indicators. For example, imperviousness can serve as an effective, direct proxy for soil sealing and habitat degradation; green space and tree cover can act as indicators of structural habitat capacity in urban ecosystems. Instead, PM_10_ concentrations can be reliable measures of air pollution pressure. All the above indicators help reveal to policy makers priority locations for some strategies of specific greening actions, such as imperviousness exceeding 40% in the settled cores of Tirana and Rijeka, or persistently low tree cover (below 5%) across the SOAS component of most urban ecosystems. This diagnostic capability can guide major policy shifts from somehow generic or vague “greening” advocacy to operational, targeted investment in areas of acute ecosystemic deficit. It can directly support national and local strategies, such as Serbia’s Spatial Plan 2035, which aims to prioritize the realization of green infrastructure (Ministarstvo građevinarstva, saobraćaja i infrastrukture, 2021), or Montenegro’s operationalization of the constitutionally normed “ecological state” principle toward measurable targets for green cover in Podgorica (Parliament of Montenegro, 2007).

Second, the multi-scale applicability of urban ecosystem accounts ensures that evidence aligns across governance levels. The results consistently show that condition values aggregated at the LAU level can obscure pronounced contrasts within urban cores (SOAS). For example, Kranj’s LAU is largely forested, while its urban core is highly sealed. Tirana’s enlarged municipal boundary masks ecological deficits in the urban core. Recognizing this scale dependency is essential for coherent policy design. National governments can rely on aggregated accounts and national average values to report progress toward international and EU commitments, such as targets under the Nature Restoration Law, and to strategically allocate funding to specific municipalities. Municipal authorities, by contrast, require finer spatial resolution. At least, SOAS level information can support the design and justification of specific interventions, such as de-sealing programs in highly impervious urban cores or tree-planting initiatives in areas with minimal canopy cover. Thematic local accounts may need even finer granularity than the one used here, linking national strategies with local drivers such as Niš’ topography, Rijeka’s constrained morphology, or Tirana’s rapid construction, which ultimately shape outcomes. Effective urban greening therefore depends on ecosystem accounts that operate across scales. Beyond their use for national reporting, cities such as Kranj or Ljubljana could apply accounts at finer-than-LAU resolution to inform zoning, target-setting (e.g., green roof ordinances), and the prioritization of investments in neighborhoods with low tree cover or pronounced heat island effect.

Third, urban ecosystem accounts provide a standardized framework for monitoring and evaluation, addressing a critical gap in many existing greening strategies. By establishing repeatable protocols for measuring ecosystem extent and condition, the accounts shift urban greening from a collection of isolated and unrelated projects to a trackable, long-term policy program. This enables outcome-oriented evaluation and adaptive management. In Croatia and Serbia, existing datasets—like the Green Infrastructure Register (Ministry of Physical Planning, Construction and State Assets of Croatia, n.d.), the Pollutant Release and

Transfer Register (Ministry of Environmental Protection and Green Transition of the Republic of Croatia, n.d.), or other national environmental statistics—offer building blocks of primary importance, but risk to remain fragmented and non-communicating with each other. Structured accounts, such as the 2018 baseline developed here, would allow evaluation of whether interventions—park rehabilitation program in Podgorica, green renewal in Croatian cities, or municipal greening in Serbia— translate into measurable improvements in ecosystem condition and services, including air filtration and access to green space.

Finally, the standardized nature of the accounts enables comparative benchmarking, fostering policy learning and knowledge exchange. The finding that Western Balkan urban ecosystems lie along a continuum of condition shared with other European cities is particularly empowering, because it facilitates the process of integration of policies and practices between EU member states and candidate states. It allows cities such as Niš, which nowadays face moderate to severe PM_10_ pollution, to benchmark their performance against cities with similar topographic constraints elsewhere in Europe and/or to identify relevant mitigation strategies applied elsewhere in similar urban ecosystems. At the regional level, the accounts establish a common language for exchanging best practices, demystifying pathways toward EU environmental standards and accelerating the Western Balkans’ green transition through a transparent, data-driven roadmap.

In sum, thematic urban ecosystem accounts may offer more than a statistical snapshot. They could function as a structured decision-support system and may help translating regulatory expectations into context-sensitive actions, by identifying degradation hotspots, and guiding targeted restoration, measuring progress toward halting urban ecological degradation.

### 4.3. Methodological value of urban ecosystem accounts and pathways for implementation

The pilot accounts demonstrate the value of urban ecosystem accounting in establishing a standardised, spatially explicit baseline of urban ecosystems, quantifying extent, condition, and service provision (e.g., air filtration). While this study presents a snapshot for 2018, the framework’s power for temporal tracking is also clear: compiling accounts over multiple years enables direct measurement of trends and provides concrete evidence of ecosystem degradation or recovery.

Standardization and multi-scale comparability are central to the methodological relevance of thematic urban ecosystem accounts. They enable comparative assessments of ecosystem extent and condition across diverse contexts, supporting the identification of widespread threat patterns, the exchange of best practices, and the calibration of policy responses to the severity of diagnosed degradation. The homogeneity observed in the Western Balkans, for example, provides a valuable baseline. Future account iterations will be critical to assess whether the region follows sustainable or divergent development trajectories, particularly in the context of further EU integration, and to inform targeted efforts to mitigate land degradation (e.g., loss of non-artificial ecosystems) and its impacts on ecological condition, biodiversity, and ecosystem services.

At the same time, the very standardization that underpins the accounts’ methodological value also raises concerns. Thematic urban ecosystem accounts are not yet formally adopted within the SEEA-EA statistical standard or its core principles and recommendations (Edens et al., 2022; United Nations, 2021). Although thematic accounts were conceived to address specific policy-relevant issues requiring the integration of multiple accounting frameworks (e.g., SEEA-EA, SEEA Central Framework, Statistical National Accounts), they remain underdeveloped as a standard. This creates a risk of divergent and incompatible approaches, potentially undermining comparability, a challenge already noted in early implementations of urban ecosystem accounts in Australia (Cryle et al., 2021).

A recent review highlights additional unresolved challenges for thematic urban ecosystem accounts that are also evident in our pilot results (Babí Almenar et al., 2026). One relates to the delineation of accounting areas. Using LAUs classified as urban under the Degree of Urbanization ensures governance relevance but introduces comparability issues due to large cross-country differences in size and urban character. For example, urban LAUs of Northern Italy are much smaller than in Serbia. Or as another example, the territorial reform that aggregated LAUs in Albania for a more economically efficient governance of municipalities distorted values for entire urban ecosystems when assumed equivalent to urban LAUs. As a result, not all territory within an “urban LAU is functionally urban, which may impede fair comparison. This also helps explain why condition values within the more homogeneous SOAS component showed greater similarity across Europe than values for entire urban ecosystems, which encompass more heterogeneous land uses.

Another challenge concerns the definition of reference levels for condition variables, a prerequisite in the SEEA-EA for creating normalized indicators (United Nations, 2021). While our results indicate that the Western Balkans often perform better in ecosystem condition variables (e.g., urban green share), the ecological meaning of these values still remain unclear. What constitutes a “good” or desirable condition (reference level) in human-dominated landscapes? This is neither clearly defined in the SEEA-EA, and only future values to be defined are anticipated in the EU NRR (European Parliament, Council of the European Union, 2024a). Furthermore, in our study the clustering of values by climate and urban class (Fig. 7) suggests that universal reference levels for the entire Europe may be inappropriate. For instance, tree cover naturally varied by climate zones, with systematically higher values in the Dfc zone (see again Fig. 7c), while air pollution levels are influenced by regional topography and atmospheric dynamics, not just emissions. For example, high concentrations in the Po Valley and Pannonian Plain (shown in Fig. 6d) are also due to regional topography and atmospheric dynamics as amplifying factors. Thus, developing meaningful, regionally contextualized, and achievable, values for reference levels remains a challenge for implementing robust, ecologically informative urban ecosystem accounts.

A final issue relevant to using thematic urban ecosystem accounts to track urban ecological degradation has been highlighted in another recent review (Stucchi et al., 2025). The authors note that urban ecosystem assessments, and by extension, the accounts built upon their methodological advances, predominantly rely on a few set of condition variables that are easy to measure. These typically belong to the *physical*, *chemical*, and *structural* state groups within the SEEA-EA condition typology. This is the case of imperviousness, PM_10_ concentration, green space, and tree cover, the condition variables justified and used in this study due to their policy relevance and data availability. While these already well-established metrics are logical candidates for regulatory monitoring, they provide only a partial picture of ecosystem and biodiversity status. Expanding methodological efforts to include compositional (e.g., species diversity), functional (e.g., evapotranspiration), and landscape-level (e.g., patch richness) metrics is essential to ensure that urban ecosystem accounts robustly inform on ecological degradation, biodiversity loss, and their implications for ecosystem service provision.

## 5. Conclusion

Our study illustrates the potential value of SEEA-EA thematic urban ecosystem accounts, already being tested in EU and EFTA countries as a tool for systematically monitoring ecosystem extent, condition and services in urban ecosystems, can easily be applied to candidate EU members so as to facilitate the application of policies and practices or to evaluate distance from operational targets. A critical first step for understanding and mitigating (or even reverting) urban ecological degradation and associated biodiversity loss in human-dominated landscapes through informed greening policies.

By establishing a spatially explicit 2018 baseline for the Western Balkans, and benchmarking it against other European countries, we demonstrate the value of thematic urban ecosystem accounts in providing a coherent, multi-scale base of evidence. These accounts enable the identification of degradation hotspots (e.g., cities with exceptionally high soil sealing) and regions of relative ecological capacity for specific services (e.g., higher normalized air filtration values due to greater effective leaf area), thereby directly informing priorities for conservation and restoration actions in urban ecosystems.

The findings within this study confirm that urban ecosystems in the Western Balkans exist on a continuum with the rest of Europe, exhibiting comparable ranges in ecosystem extent, condition, and air filtration values. This validates the transferability of the accounting framework and underscores its utility for future EU integration processes, where monitoring ecological degradation will be paramount. The accounts can help to operationalize policy goals, such as those in the EU Nature Restoration Regulation, into measurable accounts of variables, enabling targeted interventions and the tracking of progress toward halting biodiversity loss.

Looking forward, the full potential of thematic urban ecosystem accounts hinges on addressing key operational and conceptual challenges such as formalizing its standardization, like for general SEEA-EA accounts, developing ecologically meaningful and regionally contextualized reference levels, and integrating a broader suite of condition metrics (e.g., from compositional and functional state groups). Overcoming these hurdles is essential to ensure thematic urban ecosystem accounts become robust tools for informing on urban ecological degradation and its consequences for the supply of ecosystem services. As urban and peri-urban areas remain epicenters of land-use change, thematic urban ecosystem accounting is positioned as a relevant tool for evidence-based governance needed to foster resilient, biodiverse urban systems.

## 6. Supplementary Information

**Supplementary Information 1.** Detailed descriptions of each condition variable, their data sources, and spatial and temporal coverage.

**Supplementary Information 2.** Values for the distribution of local administrative units (LAUs) as cities (above and below 50.000 inhabitants) and towns and suburbs across the full study area (EUR/EFTA/UK and the Western Balkans).

**Supplementary Information 3.** National-level ecosystem extent, condition and air filtration service accounts for the urban ecosystems across the full study area (EUR/EFTA/UK and the Western Balkans). **Supplementary Information 4.** Maps of average ecosystem condition and the ecosystem service air filtration values for entire urban ecosystems and Settlements and Other Artificial Areas within urban ecosystems across the full study area (EUR/EFTA/UK and the Western Balkans).

**Supplementary Information 5.** Statistical distribution of ecosystem condition and the ecosystem service air filtration values aggregated by Köppen-Geigger climate classes (left) and urban classes for entire urban ecosystems and Settlements and Other Artificial Areas within urban ecosystems across the EUR/EFTA/UK and across the Western Balkans.

**Supplementary Information 6**. Local-level ecosystem extent, condition and air filtration service accounts for the urban ecosystems of the CROSS-REIS and NBFC pilot cities.

## 7. Acknowledgements

All authors acknowledge support from the Horizon Europe Widening project CROSS-REIS (Grant Agreement No. 101136834). JBA and RC also acknowledge support from the National Biodiversity Future Centre (NBFC) project, funded by the European Union’s Next Generation EU programme under the National Recovery and Resilience Plan (NRRP) (Project Code CN00000033; CUP D43C22001250001).

## Glossary

Countries are referred using commonly recognized acronyms, including in tables and figures, of ISO 3166-1 country codes (Alpha-2 and Alpha-3), which are provided in the glossary for the reference of readers. For clarity, UK is used as the acronym for the United Kingdom, instead of GB, in line with common policy and technical usage.

**Country Acronyms (ISO Alpha-2 and Alpha3 Codes)**

AL: Albania
AT: Austria
BE: Belgium
BIH: Bosnia and Herzegovina
BG: Bulgaria
HR: Croatia
CY: Cyprus
CZ: Czechia
DK: Denmark
EE: Estonia
FI: Finland
FR: France
DE: Germany
GR: Greece
HU: Hungary
IS: Iceland
IE: Ireland
IT: Italy
XKO: Kosovo
LV: Latvia
LI: Liechtenstein
LT: Lithuania
LU: Luxembourg
MT: Malta
ME: Montenegro
NL: Netherlands
MK: North Macedonia
NO: Norway
PL: Poland
PT: Portugal
RO: Romania
RS: Serbia
SK: Slovakia
SI: Slovenia
ES: Spain
SE: Sweden
CH: Switzerland
UK: United Kingdom

**Acronyms for Regional Grouping of Countries**

BLK: Western Balkan Countries
CR: Cross-Reis Countries
EFTA: European Free Trade Association
EU: European Union
EUR: Other European Countries (EFTA countries, UK and European Union Countries, except Croatia and Slovenia)

**Acronyms for pilot municipalities**

CA: Campobasso
KR: Kranj
FI: Firenze
MI: Milano
NA: Napoli
PO: Podgorica
RO: Roma
RI: Rijeka
TI: Tirana
TO: Torino

**Acronyms of Köppen–Geiger climate classes**

BSh: Hot semi-arid (steppe)
BSk: Cold semi-arid (steppe)
BWh: Hot desert
BWk: Cold desert
Cfa: Humid subtropical
Cfb: Oceanic (marine west coast)
Csa: Hot-summer Mediterranean
Csb: Warm-summer Mediterranean
Dfa: Humid continental hot summer
Dfb: Humid continental warm summer
Dfc: Subarctic with cool short summer
ET: Tundra

**Other acronyms**

NRR: Nature Restoration Regulation
SEEA-EA: System of Environmental-Economic Accounting–Ecosystem Accounting
SOAS: Settlements and Other Artificial Areas

