## Supplementary Material 1 for "Exploring the value of thematic urban ecosystem accounts in Western Balkan Countries to inform urban greening actions"

Supplementary Information 1. Detailed descriptions of each condition variable, their data sources, and spatial and temporal coverage.

| Ecosystem condition variable | Description and data sources. | Spatial Resolution | Temporal Resolution |
|---|---|---|---|
| Imperviousness share | Imperviousness share measures the sealing density of an entire urban ecosystem area or a specific Ecosystem Type Level 1 within that area in the range of 0 to 100%.<br><br>This condition variable is estimated using the High-Resolution Layer Imperviousnes (Imperviousness Density) for 2018 of Copernicus Land Monitoring Service.<br>DOI: https://doi.org/10.2909/3bf542bd-eebd-4d73-b53c-a0243f2ed862 | 10 m | 3-yearly |
| Urban green spaces share | Urban green spaces correspond to any piece of urban land covered with mosses and lichens, herbs, shrubs, or trees, which do not correspond to arable land. Urban green spaces can be of public, semi-private, or private ownership and be present in land of different built-up character (e.g., ground open space, building rooftop, building façade). Therefore, urban green share is defined as the proportion (ranging from 0 to 100%) of an entire urban ecosystem area, or of a specific Ecosystem Type Level 1 within that area, that is covered by vegetated surfaces, regardless of ownership type or the degree of surrounding built-up structures.<br><br>This condition variable is estimated using the CLC plus Backbone dataset for 2018 of Copernicus Land Monitoring Service.<br>DOI: https://doi.org/10.2909/cd534ebf-f553-42f0-9ac1-62c1dc36d32c | 10 m | 3-yearly |
| Urban tree canopy cover share | Urban tree canopy cover share is defined as the proportion (ranging from 0 to 100%) of an entire urban ecosystem area, or of a specific Ecosystem Type Level 1 withing that area, that is covered with tree canopy (i.e., the vertical projection of tree crowns to a horizontal earth's surface).<br><br>This condition variable is estimated using the High-Resolution Layer Tree Cover Density of 2018 of Copernicus Land Monitoring Service.<br>DOI: https://doi.org/10.2909/e677441e-fb94-431c-b4f9-304f10e4dfd8 | 10 m | yearly |
| Annual average air concentration of $PM_{10}$ | Annual average air concentration of $PM_{10}$ is the mean concentration of particulate matter with an aerodynamic diameter smaller than 10 µm measured over a one-year period at an hourly temporal resolution.<br><br>This condition variable is estimated using the Copernicus Atmosphere Monitoring Service - European air quality reanalyses – Dataset of Particulate matter < 10 µm (PM10)<br>DOI: 10.24381/7cc0465a | 0.1° | hourly |
