## Supplementary Material 2 for "Exploring the value of thematic urban ecosystem accounts in Western Balkan Countries to inform urban greening actions"

Supplementary Information 2. Values for the distribution of local administrative units (LAUs) as cities (above and below 50.000 inhabitants) and towns and suburbs across the full study area (EUR/EFTA/UK and the Western Balkans).

Units: Land share

| | Country | Cities above 50.000 inh. (Urban Class 0) | Cities below 50.000 inh. (Urban Class 1) | Towns and Suburbs (Urban Class 2) | Rural |
|---|---|---|---|---|---|
| | **ME** | 10,5% | 0,0% | 59,0% | 30,5% |
| | **RS** | 6,9% | 0,0% | 53,7% | 39,4% |
| | **AL** | 10,5% | 0,0% | 21,9% | 67,5% |
| | **HR** | 2,0% | 0,1% | 16,9% | 81,0% |
| | **SI** | 2,1% | 0,0% | 23,5% | 74,4% |
| | **MK** | 11,7% | 0,3% | 48,5% | 39,5% |
| | **BIH** | 3,9% | 0,3% | 60,2% | 35,6% |
| | **XKO** | 8,5% | 0,0% | 52,6% | 38,9% |
| | **LI** | 0,0% | 0,0% | 94,7% | 5,3% |
| | **MT** | 0,0% | 27,7% | 60,3% | 12,0% |
| | **NL** | 16,5% | 2,0% | 55,3% | 26,2% |
| | **UK** | 11,7% | 0,0% | 46,7% | 41,6% |
| | **BE** | 4,5% | 1,7% | 47,0% | 46,8% |
| | **BG** | 7,2% | 0,0% | 33,6% | 59,2% |
| | **IT** | 5,0% | 1,3% | 34,3% | 59,4% |
| | **DE** | 4,7% | 0,5% | 31,9% | 62,9% |
| | **DK** | 2,1% | 0,5% | 31,6% | 65,8% |
| | **CH** | 1,1% | 1,9% | 30,6% | 66,4% |
| | **HU** | 3,3% | 0,1% | 23,9% | 72,8% |
| | **LU** | 2,0% | 2,1% | 21,7% | 74,1% |
| | **ES** | 4,9% | 0,4% | 20,4% | 74,3% |
| | **SE** | 2,9% | 0,0% | 21,0% | 76,1% |
| | **PL** | 2,2% | 0,2% | 16,8% | 80,8% |
| | **SK** | 1,0% | 0,4% | 16,1% | 82,5% |
| | **LT** | 1,1% | 0,0% | 15,4% | 83,6% |
| | **CZ** | 2,3% | 0,1% | 13,7% | 84,0% |
| | **AT** | 1,2% | 0,2% | 14,2% | 84,4% |
| | **PT** | 0,2% | 1,1% | 14,1% | 84,7% |
| | **FI** | 0,5% | 0,0% | 14,5% | 85,0% |
| | **RO** | 1,7% | 0,2% | 12,9% | 85,3% |
| | **CY** | 1,6% | 2,1% | 10,2% | 86,1% |
| | **FR** | 0,7% | 0,8% | 9,2% | 89,3% |
| | **LV** | 0,7% | 0,0% | 10,0% | 89,4% |
| | **NO** | 0,8% | 0,0% | 8,5% | 90,7% |
| | **IE** | 0,2% | 1,3% | 7,5% | 91,0% |
| | **EE** | 0,9% | 0,0% | 7,5% | 91,6% |
| | **EL** | 0,5% | 0,3% | 6,3% | 92,9% |
| | **IS** | 0,2% | 0,1% | 0,8% | 98,8% |
| **Weighted Average** | **All-BLK** | 6,1% | 0,1% | 41,9% | 51,9% |
| | **All-C-R** | 5,8% | 0,0% | 35,8% | 58,4% |
| | **All-Eur** | 3,0% | 0,4% | 19,7% | 76,9% |

Units: Population share

| | Country | Cities above 50.000 inh. (Urban Class 0) | Cities below 50.000 inh. (Urban Class 1) | Towns and Suburbs (Urban Class 2) | Rural |
|---|---|---|---|---|---|
| | ME | 33,0% | 0,0% | 55,9% | 11,1% |
| | RS | 26,6% | 1,9% | 49,7% | 21,8% |
| | AL | 38,3% | 0,0% | 31,5% | 30,1% |
| | HR | 31,2% | 0,6% | 33,7% | 34,5% |
| | SI | 19,5% | 0,0% | 36,9% | 43,6% |
| | MK | 32,8% | 4,7% | 51,8% | 10,8% |
| | BIH | 13,9% | 5,8% | 63,9% | 16,4% |
| | XKO | 17,3% | 0,0% | 60,4% | 22,3% |
| | MT | 0,0% | 64,2% | 33,8% | 2,0% |
| | LI | 0,0% | 0,0% | 96,6% | 3,4% |
| | UK | 58,6% | 0,0% | 34,8% | 6,5% |
| | NL | 47,7% | 5,0% | 40,1% | 7,2% |
| | BE | 24,8% | 8,5% | 53,7% | 13,0% |
| | ES | 50,1% | 5,0% | 31,6% | 13,4% |
| | IT | 29,3% | 8,6% | 45,5% | 16,6% |
| | CH | 16,4% | 14,4% | 51,2% | 18,0% |
| | IS | 37,8% | 11,0% | 33,0% | 18,2% |
| | DE | 35,8% | 2,2% | 41,6% | 20,3% |
| | SE | 36,8% | 2,0% | 40,4% | 20,8% |
| | BG | 43,0% | 0,0% | 32,7% | 24,3% |
| | PT | 4,6% | 29,7% | 39,9% | 25,8% |
| | CY | 24,6% | 18,3% | 30,8% | 26,2% |
| | FI | 27,9% | 0,2% | 42,4% | 29,4% |
| | EL | 34,3% | 11,2% | 24,7% | 29,8% |
| | LU | 19,1% | 5,7% | 45,1% | 30,0% |
| | HU | 32,1% | 0,3% | 36,4% | 31,1% |
| | NO | 23,7% | 1,1% | 42,8% | 32,3% |
| | FR | 22,3% | 14,2% | 29,1% | 34,4% |
| | DK | 21,3% | 4,8% | 39,4% | 34,5% |
| | LV | 39,7% | 0,0% | 24,5% | 35,7% |
| | CZ | 27,4% | 0,7% | 36,0% | 35,9% |
| | PL | 32,7% | 1,1% | 30,3% | 35,9% |
| | AT | 31,9% | 2,2% | 29,2% | 36,7% |
| | EE | 43,3% | 0,0% | 18,5% | 38,2% |
| | RO | 36,1% | 0,9% | 24,0% | 39,0% |
| | SK | 9,8% | 5,1% | 43,5% | 41,6% |
| | LT | 38,2% | 0,0% | 19,1% | 42,6% |
| | IE | 9,5% | 23,1% | 21,1% | 46,3% |
| Weighted Average | All-BLK | 26,3% | 1,9% | 46,9% | 24,8% |
| | All-C-R | 29,1% | 1,0% | 41,4% | 28,6% |
| | All-Eur | 35,3% | 5,6% | 36,2% | 22,9% |
