## Supplementary Material 4 for "Exploring the value of thematic urban ecosystem accounts in Western Balkan Countries to inform urban greening actions"

**Supplementary Information 4.** Maps of average ecosystem condition and the ecosystem service air filtration values for entire urban ecosystems and Settlements and Other Artificial Areas within urban ecosystems across the full study area (EUR/EFTA/UK and the Western Balkans).

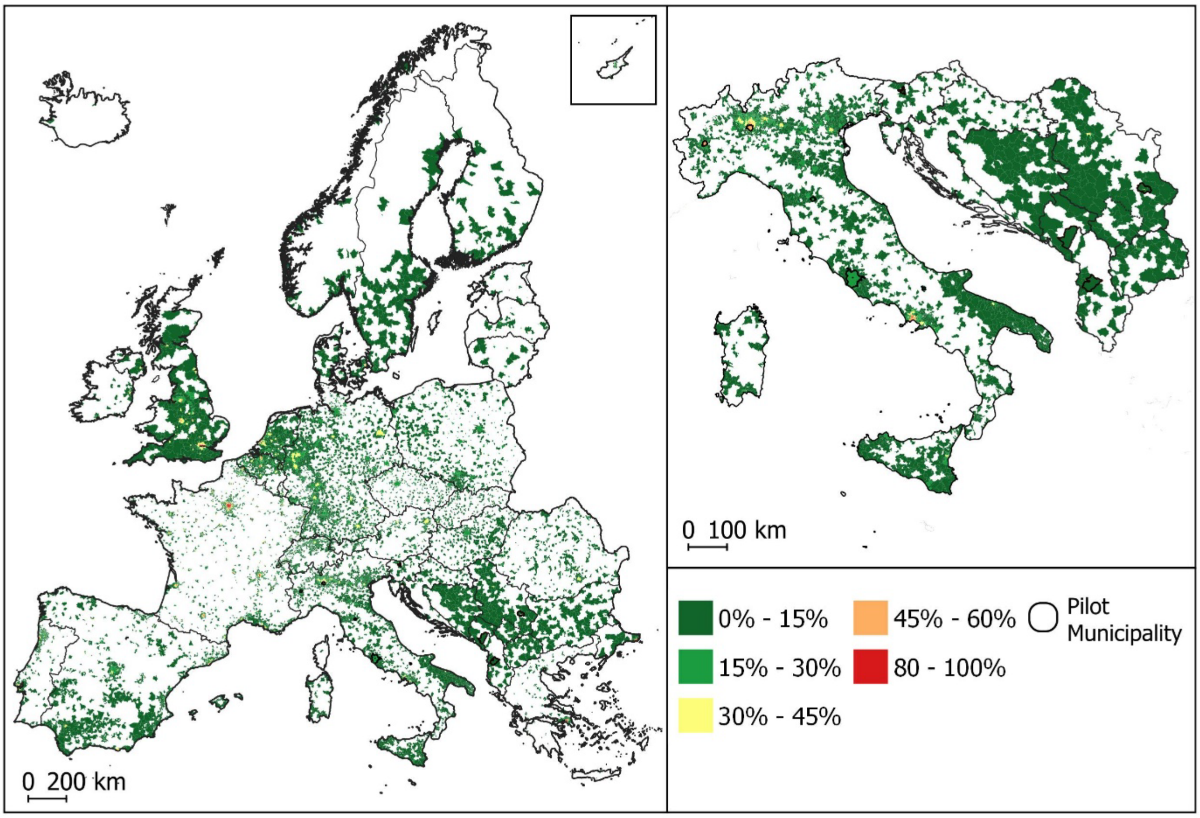


**Figure SM4.1. Mapping of the ecosystem condition variable imperviousness share for urban ecosystems for the entire area of scope, with a zoom-in on the Western Balkans and Italy.** Values are presented per urban ecosystem (i.e., urban local administrative unit).

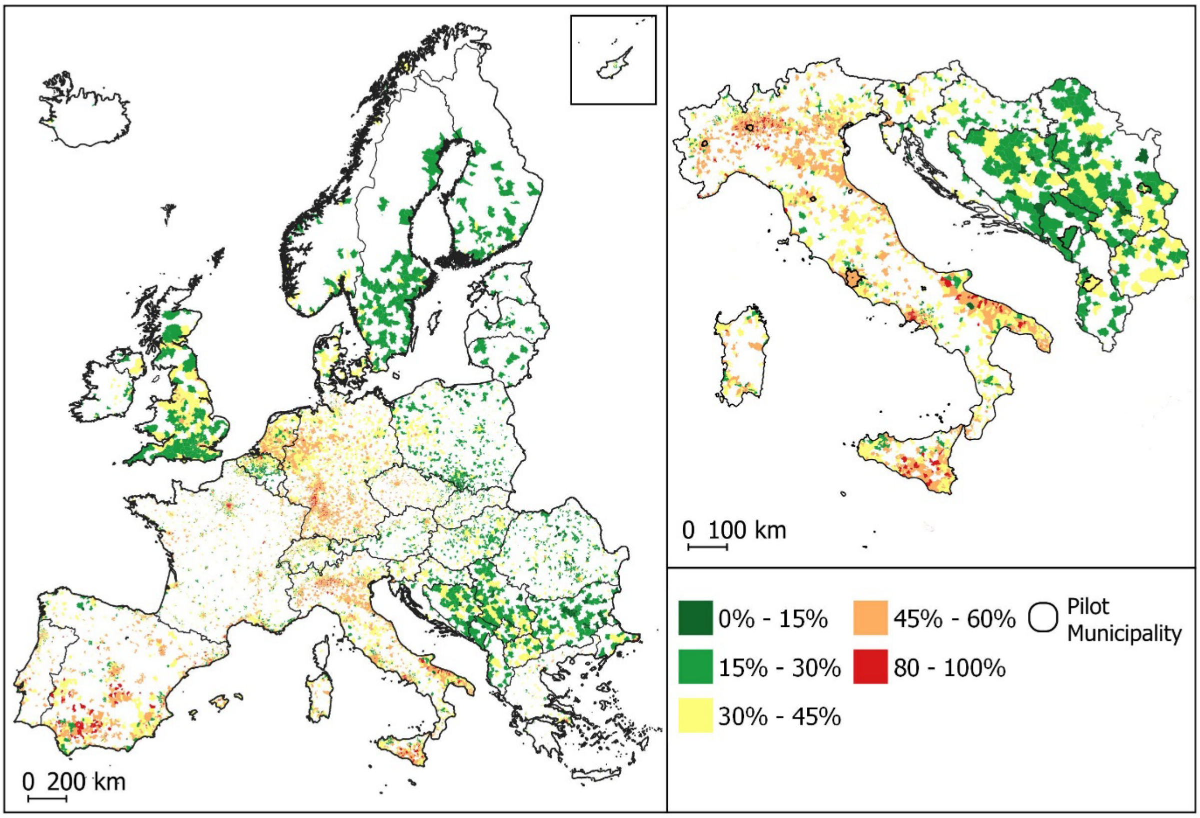


**Figure SM4.2. Mapping of the ecosystem condition variable imperviousness share in the *Settlements and Other Artificial Areas* of urban ecosystems for the entire area of scope, with a zoom-in on the Western Balkans and Italy.** Values are presented per urban ecosystem (i.e., urban local administrative unit).

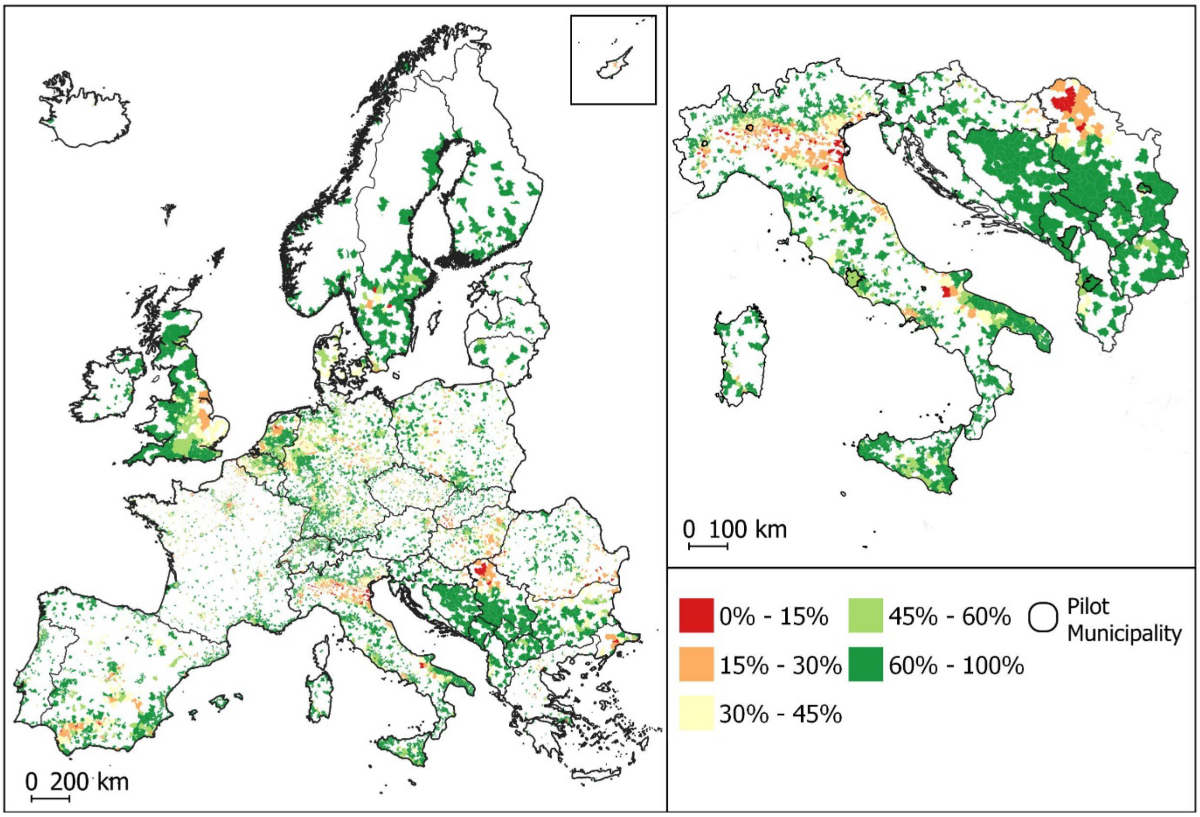


**Figure SM4.3. Mapping of the ecosystem condition variable urban green share for urban ecosystems for the entire area of scope, with a zoom-in on the Western Balkans and Italy.** Values are presented per urban ecosystem (i.e., urban local administrative unit).

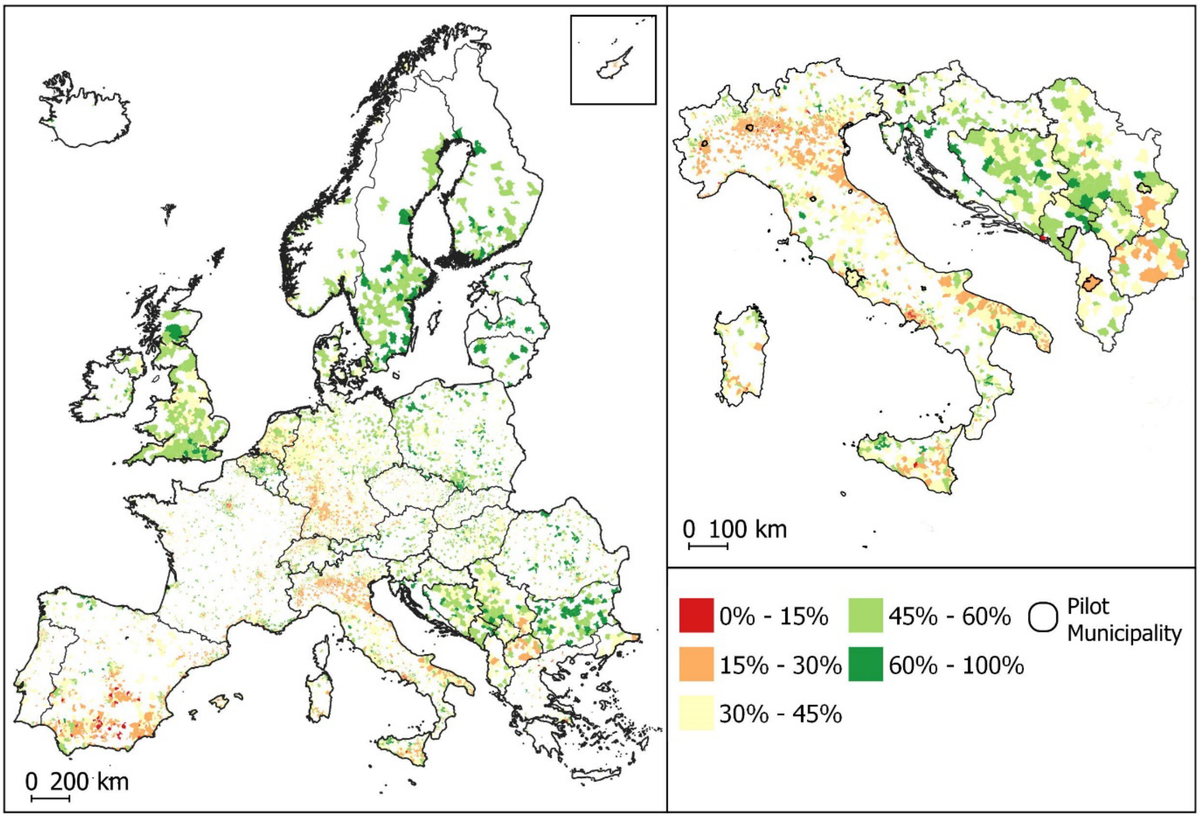


**Figure SM4.4. Mapping of the ecosystem condition variable urban green share in the *Settlements and Other Artificial Areas* of urban ecosystems for the entire area of scope, with a zoom-in on the Western Balkans and Italy.** Values are presented per urban ecosystem (i.e., urban local administrative unit).

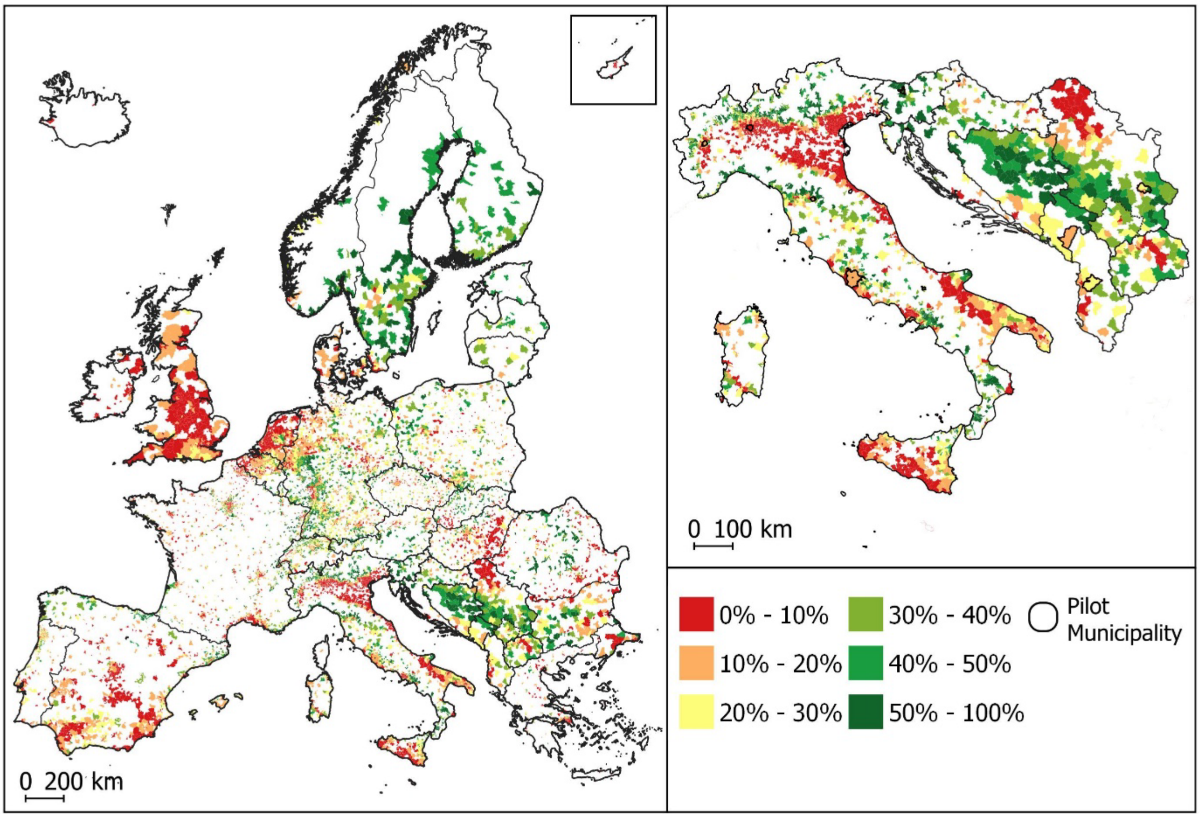


**Figure SM4.5. Mapping of the ecosystem condition variable tree cover share for urban ecosystems for the entire area of scope, with a zoom-in on the Western Balkans and Italy.** Values are presented per urban ecosystem (i.e., urban local administrative unit).

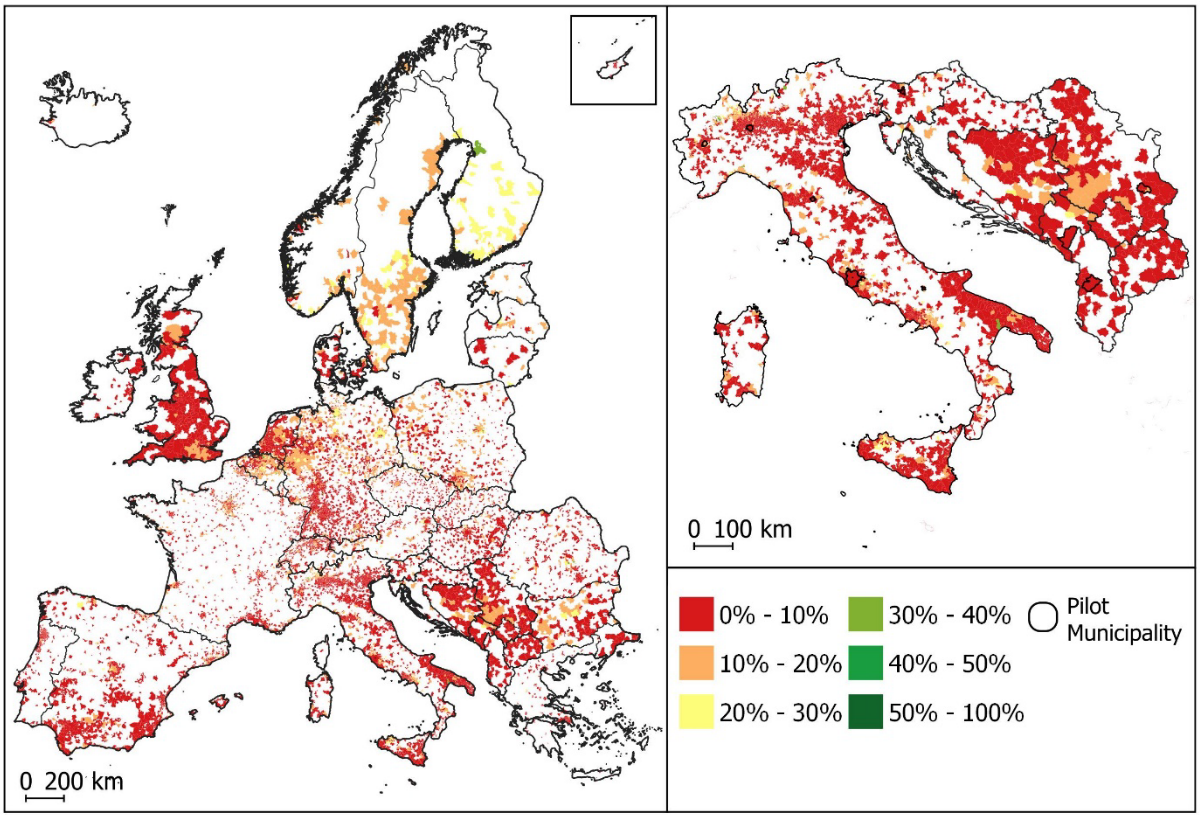


**Figure SM4.6. Mapping of the ecosystem condition variable tree cover share in the *Settlements and Other Artificial Areas* of urban ecosystems for the entire area of scope, with a zoom-in on the Western Balkans and Italy.** Values are presented per urban ecosystem (i.e., urban local administrative unit).

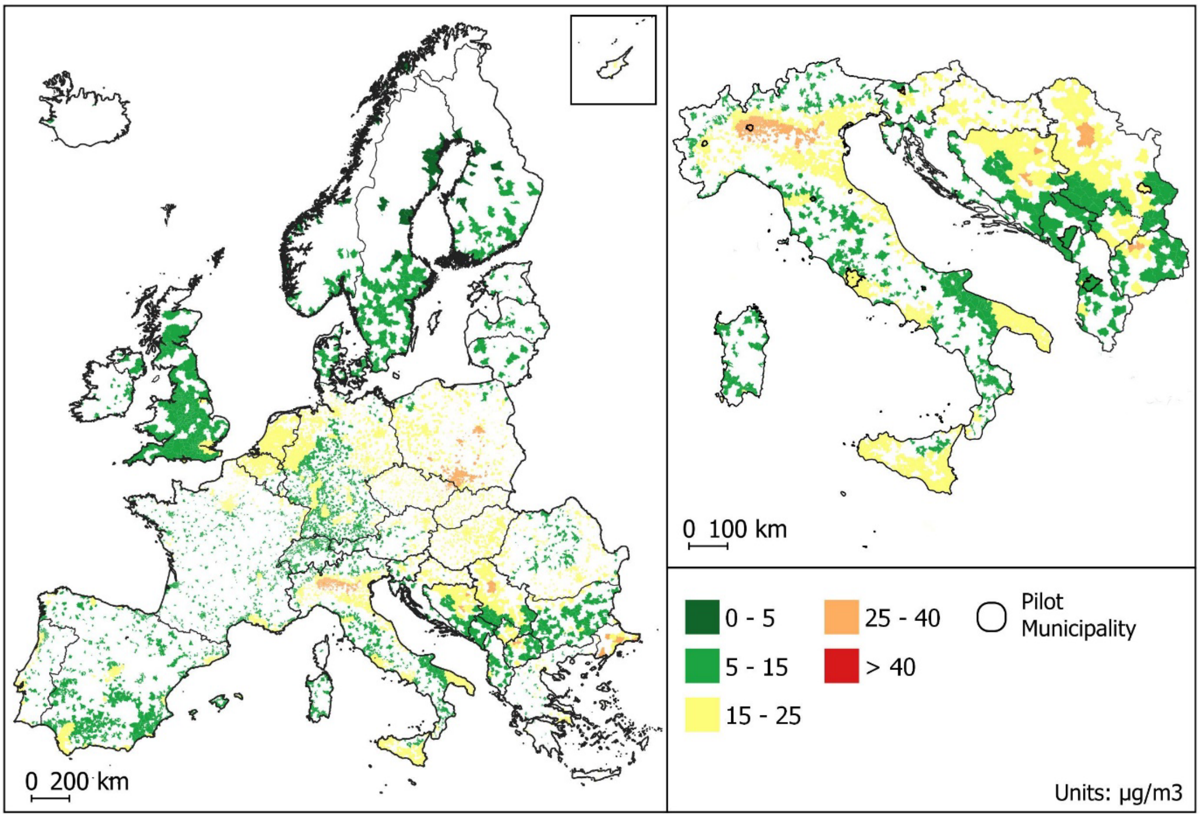


**Figure SM4.7. Mapping of the ecosystem condition variable $PM_{10}$ atmospheric concentration for urban ecosystems for the entire area of scope, with a zoom-in on the Western Balkans and Italy.** Values are presented per urban ecosystem (i.e., urban local administrative unit).

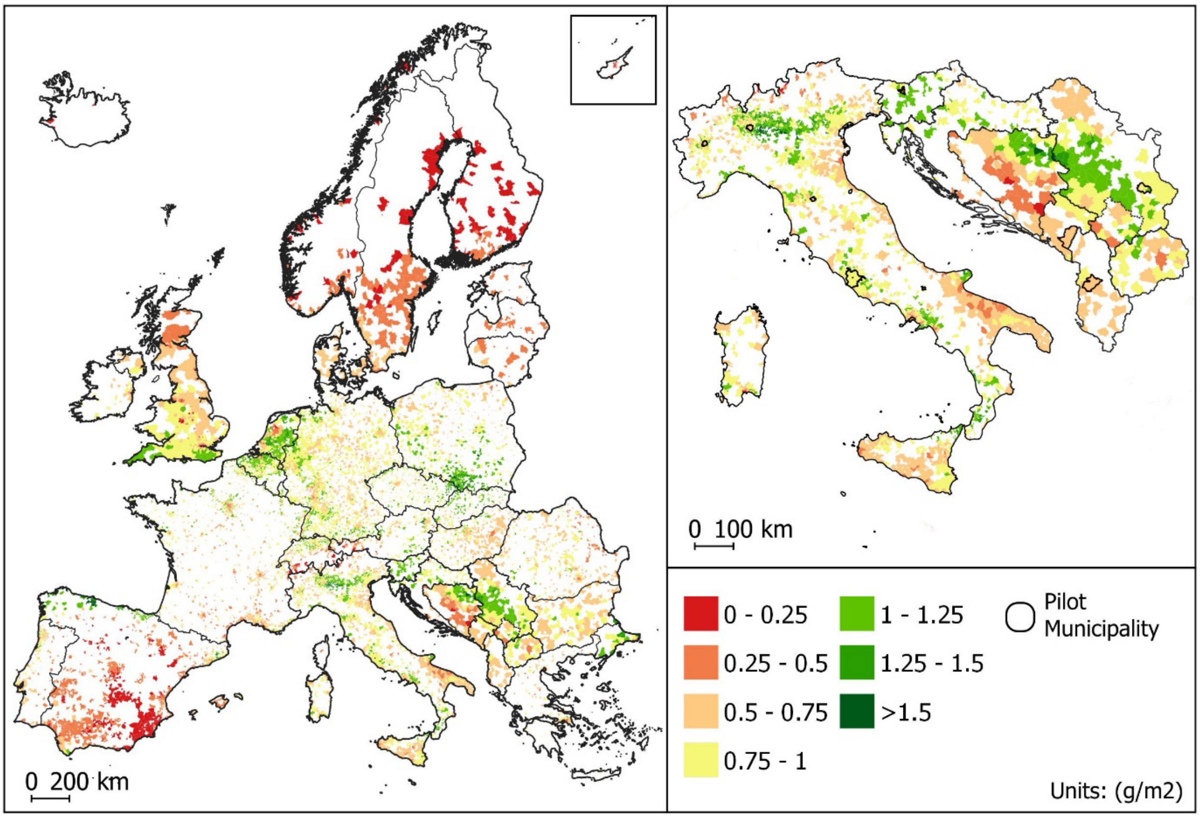


**Figure SM4.8. Mapping of the ecosystem condition variable $PM_{10}$ deposition per land unit (g/m$^2$) for urban ecosystems for the entire area of scope, with a zoom-in on the Western Balkans and Italy.** Values are presented per urban ecosystem (i.e., urban local administrative unit).

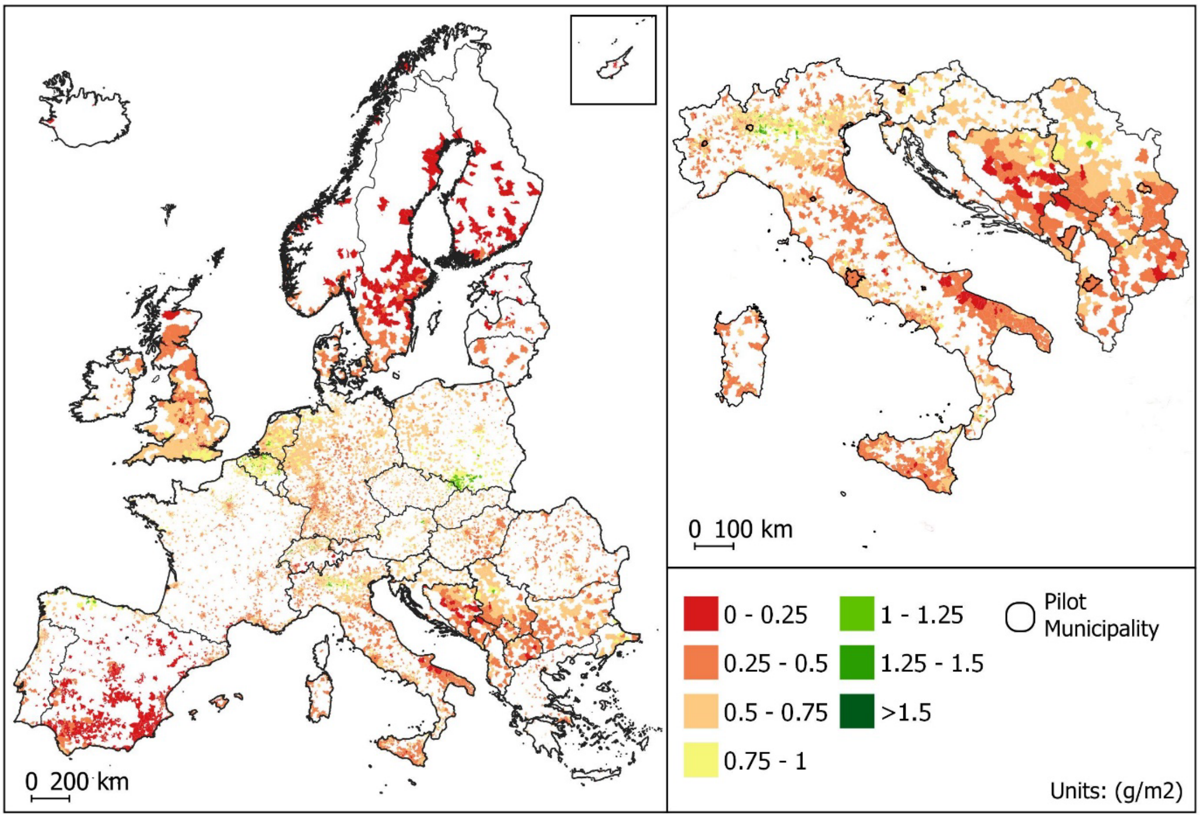


**Figure SM4.9. Mapping of the ecosystem condition variable $PM_{10}$ deposition per land unit (g/m$^2$) in the *Settlements and Other Artificial Areas* of urban ecosystems for the entire area of scope, with a zoom-in on the Western Balkans and Italy.** Values are presented per urban ecosystem (i.e., urban local administrative unit).

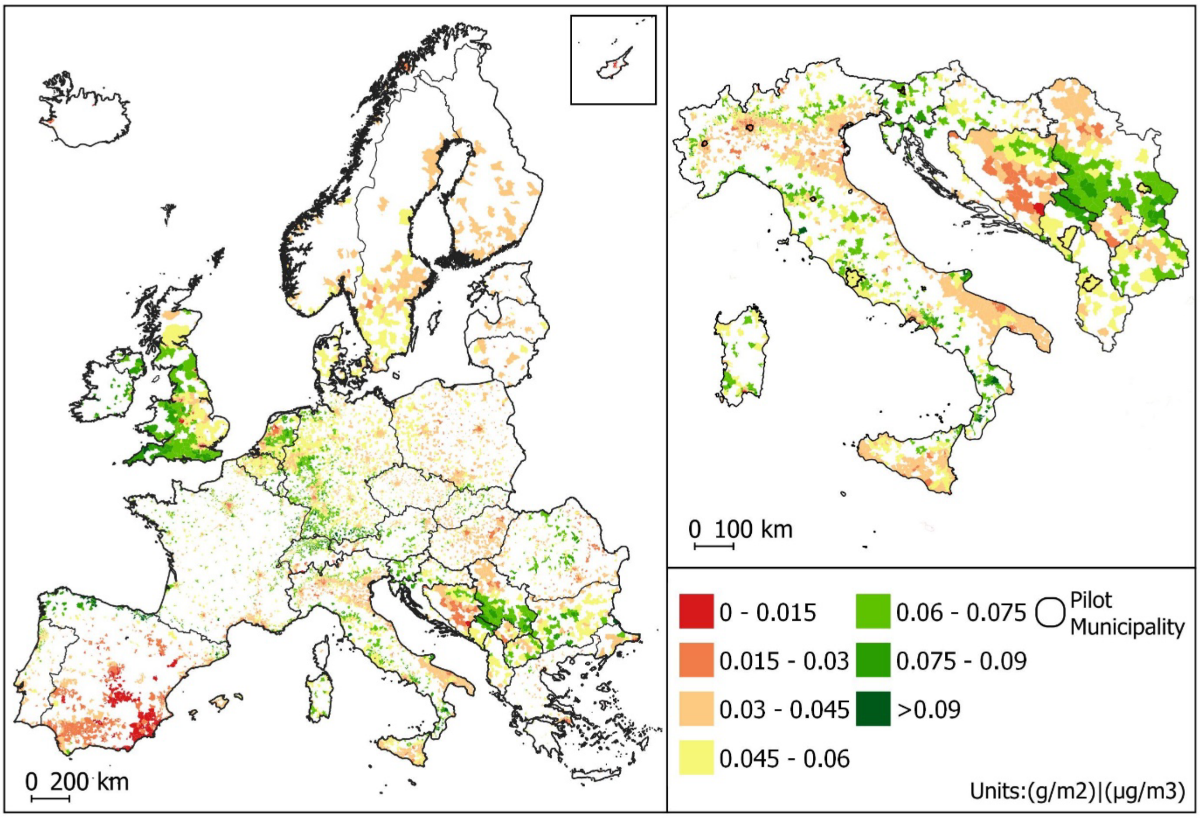


**Figure SM4.10. Mapping of the ecosystem condition variable $PM_{10}$ deposition per land unit ($g/m^2$) normalized by average annual atmospheric $PM_{10}$ levels ($\mu g/m^3$) for urban ecosystems for the entire area of scope, with a zoom-in on the Western Balkans and Italy.** Values are presented per urban ecosystem (i.e., urban local administrative unit).

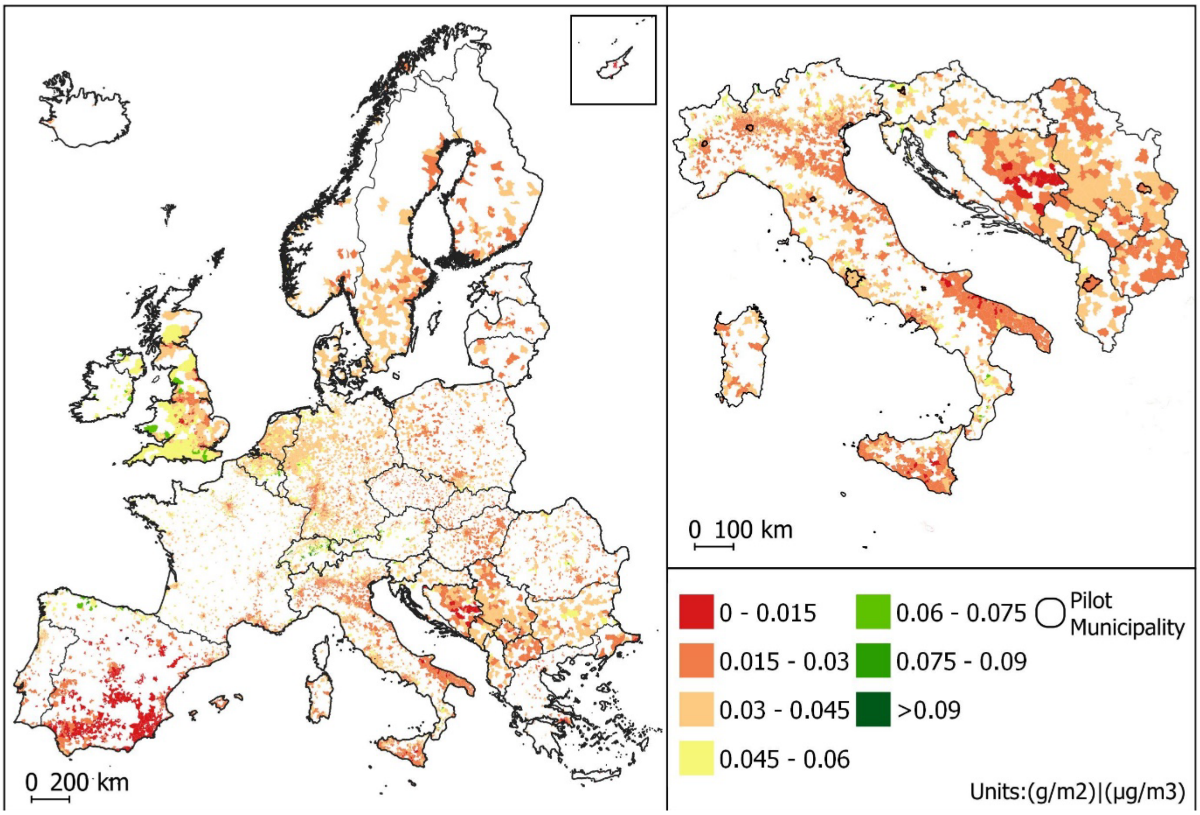


**Figure SM4.11. Mapping of the ecosystem condition variable $PM_{10}$ deposition per land unit ($g/m^2$) normalized by average annual atmospheric $PM_{10}$ levels ($\mu g/m^3$) in the *Settlements and Other Artificial Areas* of urban ecosystems for the entire area of scope, with a zoom-in on the Western Balkans and Italy.** Values are presented per urban ecosystem (i.e., urban local administrative unit).
