## Supplementary Material 5 for "Exploring the value of thematic urban ecosystem accounts in Western Balkan Countries to inform urban greening actions"

**Supplementary Information 5.** Statistical distribution of ecosystem condition and the ecosystem service air filtration values aggregated by Köppen-Geigger climate classes and urban classes for entire urban ecosystems and Settlements and Other Artificial Areas within urban ecosystems across the Western Balkans and EUR/EFTA/UK countries.

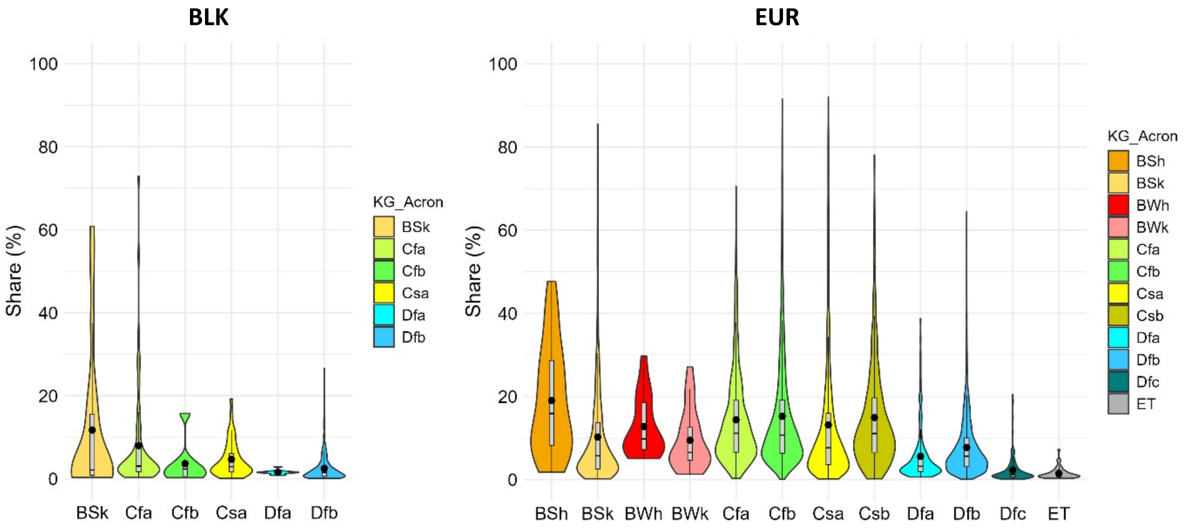


**Figure SM5.1. Statistical distribution of the ecosystem condition variable Imperviousness share in urban ecosystems, aggregated by Köppen-Geigger climate classes. Violin plots show values for the Western Balkans (BLK)and all European countries (EUR).**

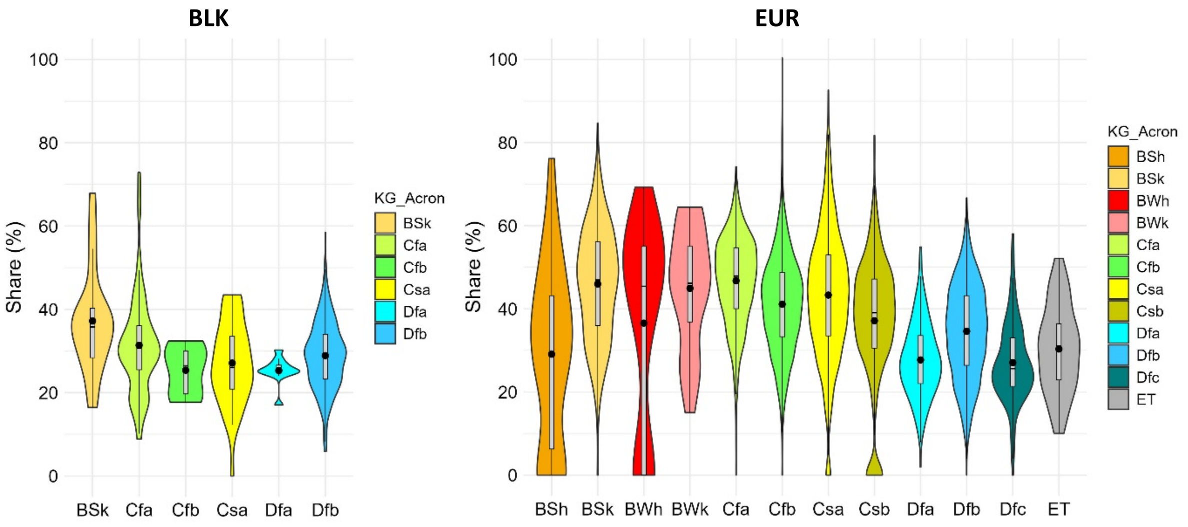


**Figure SM5.2. Statistical distribution of the ecosystem condition variable Imperviousness share in *Settlements and Other Artificial Areas* of urban ecosystems, aggregated by Köppen-Geigger climate classes. Violin plots show values for the Western Balkans (BLK)and all European countries (EUR).**

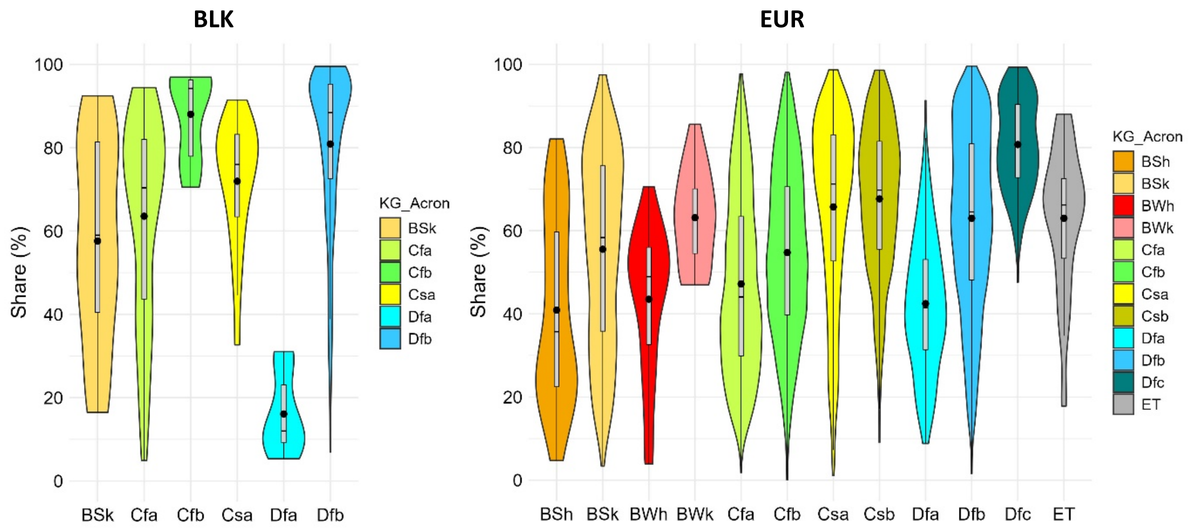


**Figure SM5.3. Statistical distribution of the ecosystem condition variable urban green share in urban ecosystems, aggregated by Köppen-Geigger climate classes. Violin plots show values for the Western Balkans (BLK)and all European countries (EUR).**

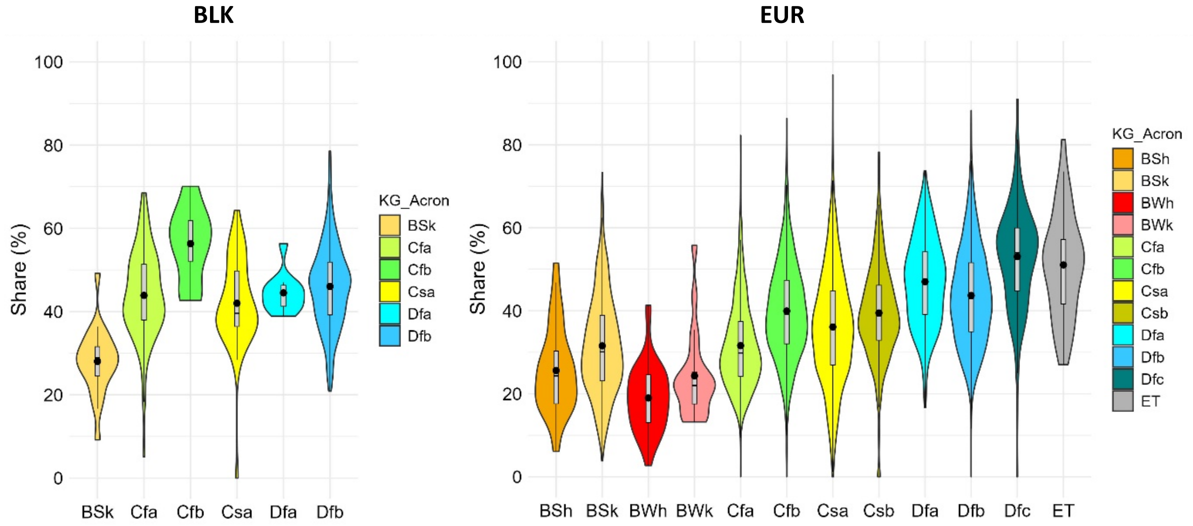


**Figure SM5.4. Statistical distribution of the ecosystem condition variable urban green share in *Settlements and Other Artificial Areas* of urban ecosystems, aggregated by Köppen-Geigger climate classes. Violin plots show values for the Western Balkans (BLK)and all European countries (EUR).**

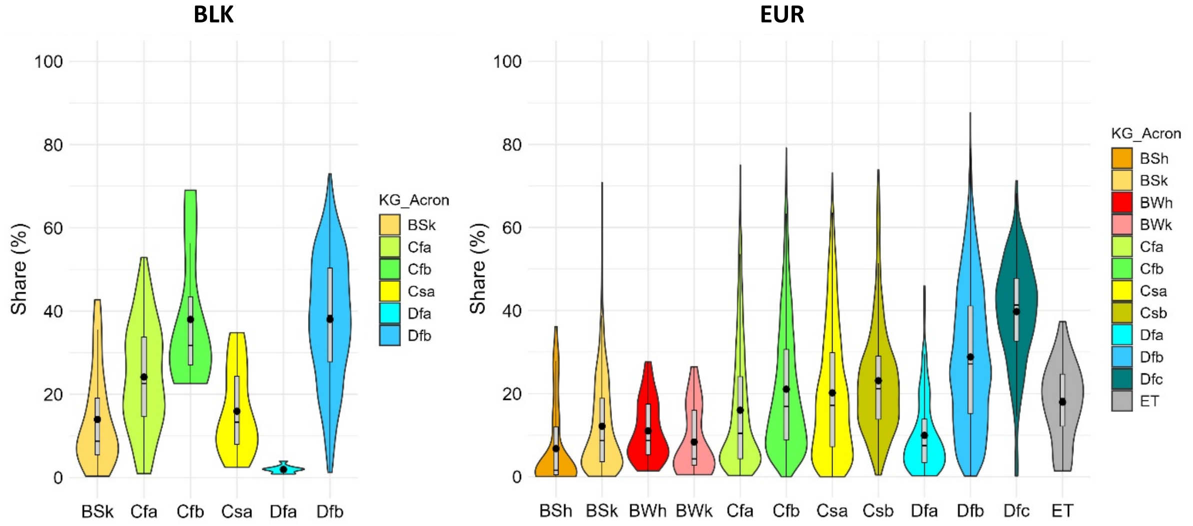


**Figure SM5.5. Statistical distribution of the ecosystem condition variable urban tree cover share in urban ecosystems, aggregated by Köppen-Geigger climate classes. Violin plots show values for the Western Balkans (BLK)and all European countries (EUR).**

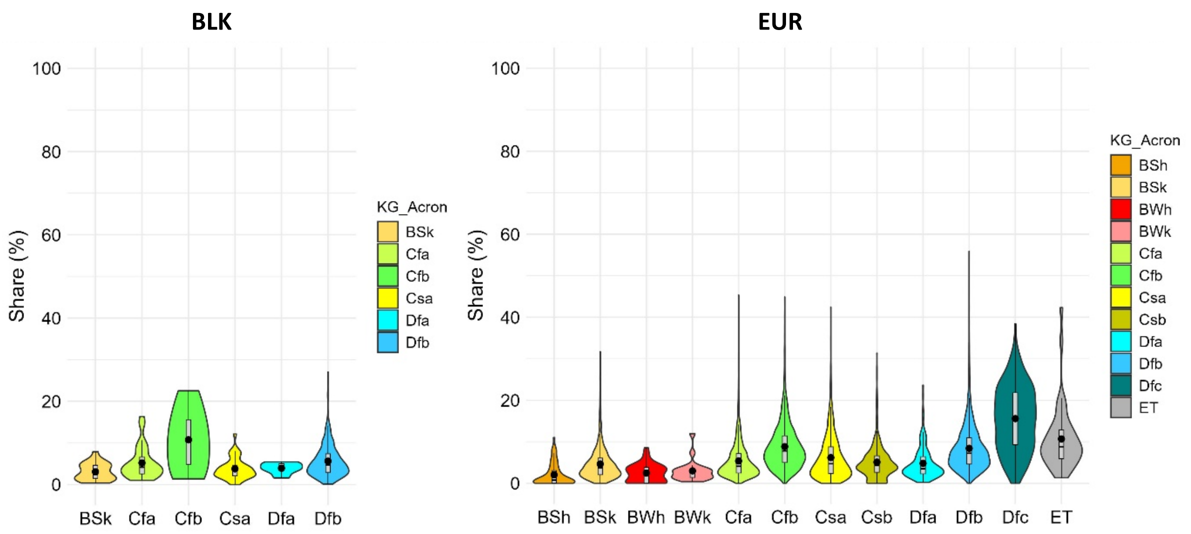


**Figure SM5.6. Statistical distribution of the ecosystem condition variable urban tree cover share in *Settlements and Other Artificial Areas* of urban ecosystems, aggregated by Köppen-Geigger climate classes. Violin plots show values for the Western Balkans (BLK)and all European countries (EUR).**

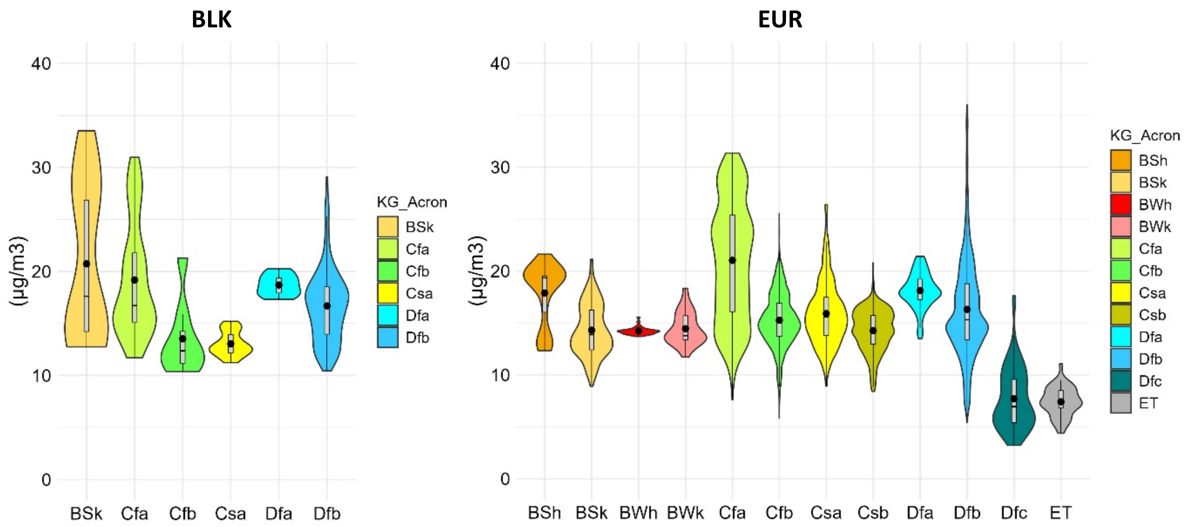


**Figure SM5.7. Statistical distribution of the ecosystem condition variable average annual $PM_{10}$ atmospheric concentration in urban ecosystems, aggregated by Köppen-Geigger climate classes. Violin plots show values for the Western Balkans (BLK)and all European countries (EUR).**

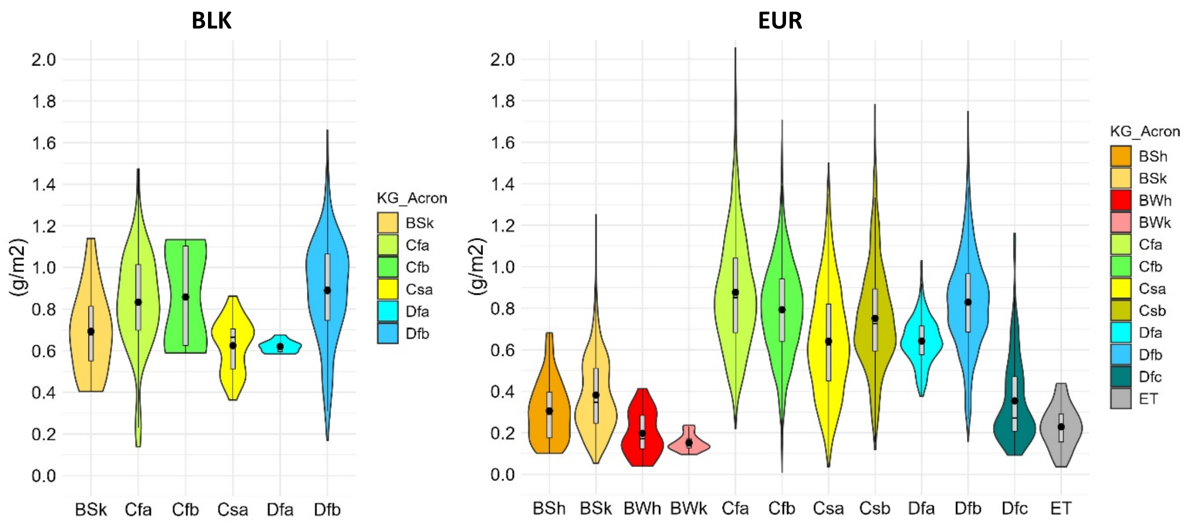


**Figure SM5.8. Statistical distribution of air filtration service as $PM_{10}$ deposition per land unit (g/m²) in urban ecosystems, aggregated by Köppen-Geigger climate classes. Violin plots show values for the Western Balkans (BLK)and all European countries (EUR).**

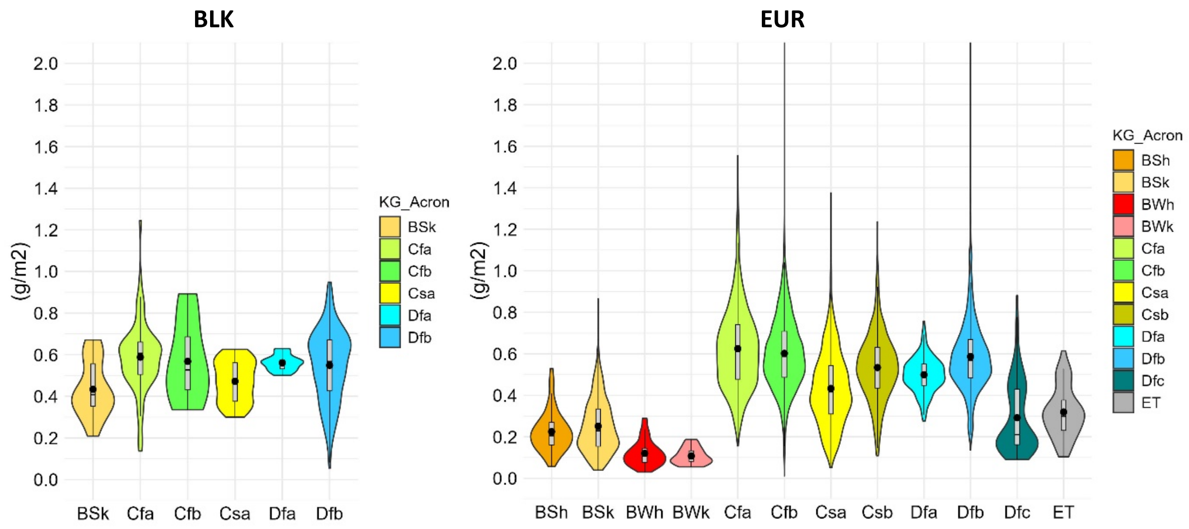


**Figure SM5.9. Statistical distribution of air filtration service as $PM_{10}$ deposition per land unit ($g/m^2$) in *Settlements and Other Artificial Areas* of urban ecosystems, aggregated by Köppen-Geigger climate classes. Violin plots show values for the Western Balkans (BLK)and all European countries (EUR).**

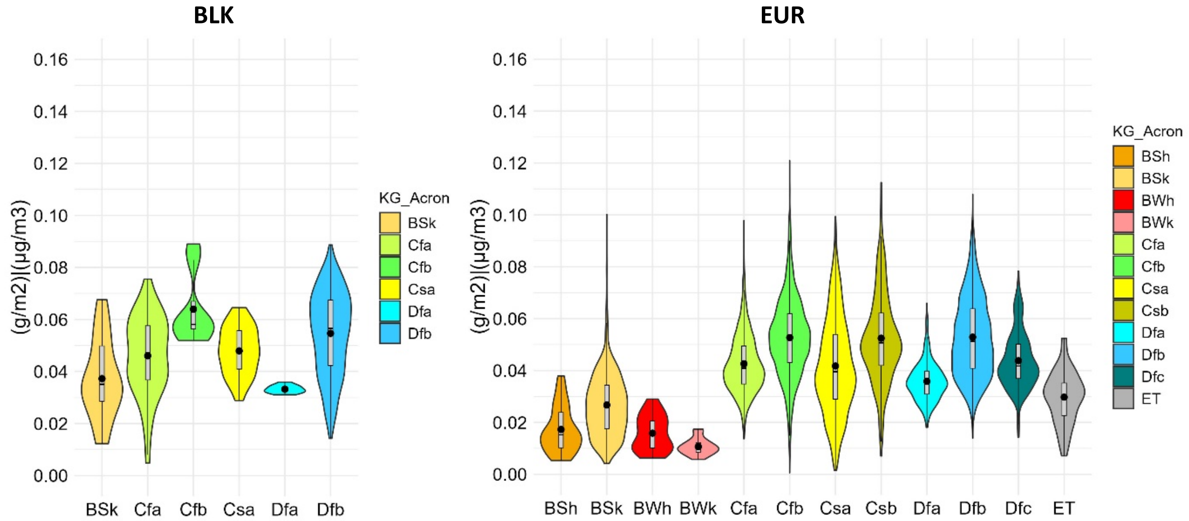


**Figure SM5.10. Statistical distribution of air filtration service as $PM_{10}$ deposition per land unit ($g/m^2$) normalized by average annual atmospheric $PM_{10}$ levels ($\mu g/m^3$) in urban ecosystems, aggregated by Köppen-Geigger climate classes. Violin plots show values for the Western Balkans (BLK)and all European countries (EUR).**

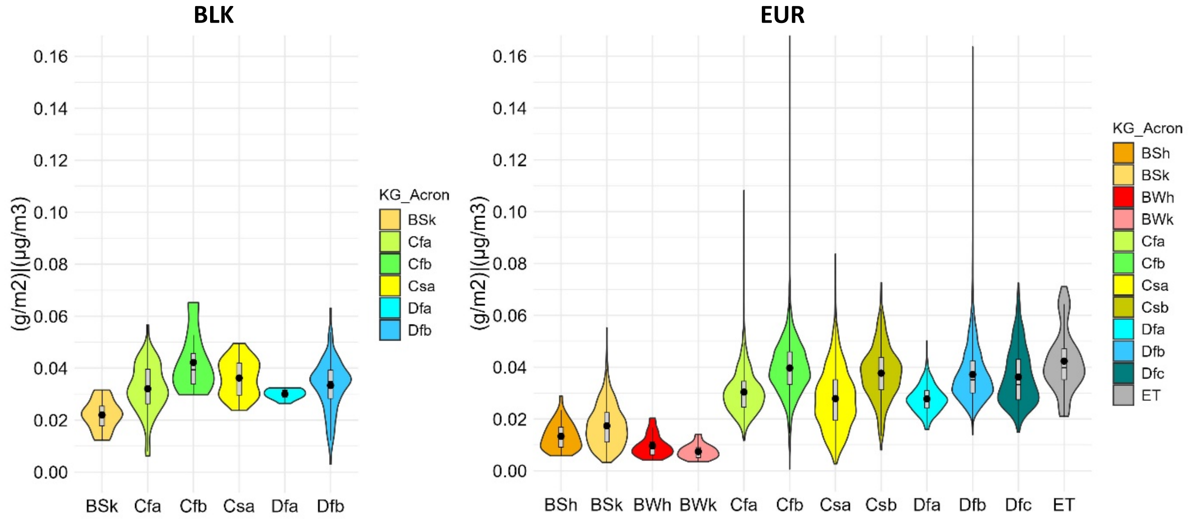


**Figure SM5.11. Statistical distribution of air filtration service as $PM_{10}$ deposition per land unit ($g/m^2$) normalized by average annual atmospheric $PM_{10}$ levels ($\mu g/m^3$) in *Settlements and Other Artificial Areas* of urban ecosystems, aggregated by Köppen-Geigger climate classes. Violin plots show values for the Western Balkans (BLK)and all European countries (EUR).**

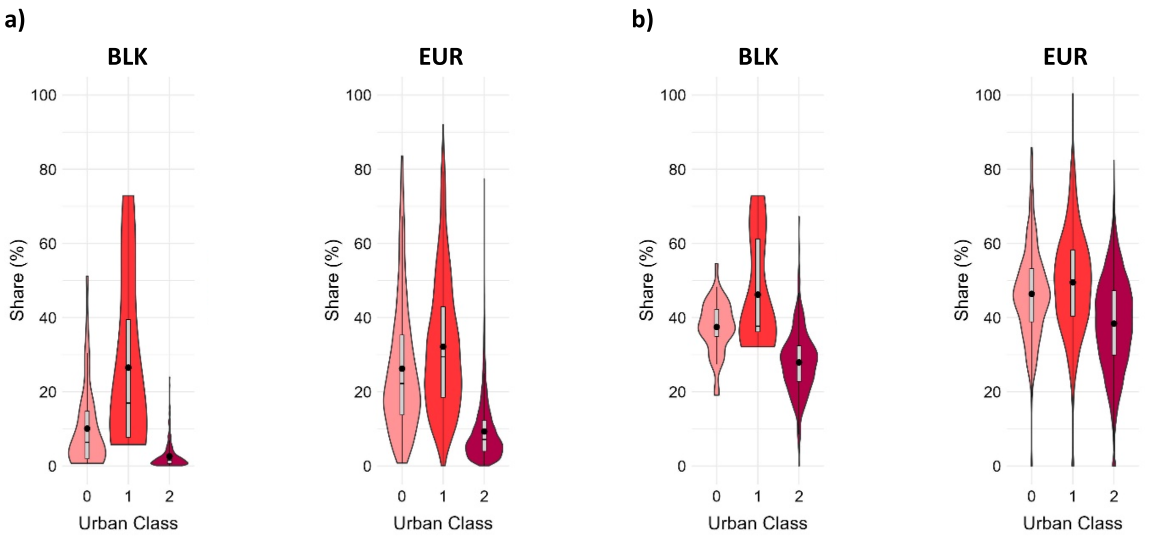


**Figure SM5.12. Statistical distribution of the ecosystem condition variable imperviousness share in urban ecosystems (a) and in *Settlements and Other Artificial Areas* of urban ecosystems (b) aggregated by urban classes. Violin plots show values for the Western Balkans (BLK)and all European countries (EUR).** Class 0 = local administrative units classified as cities above or equal to 50.000 inhabitants; Class 1 = local administrative units classified as cities below 50.000 inhabitants; Class 2 = local administrative units classified as towns and suburbs.

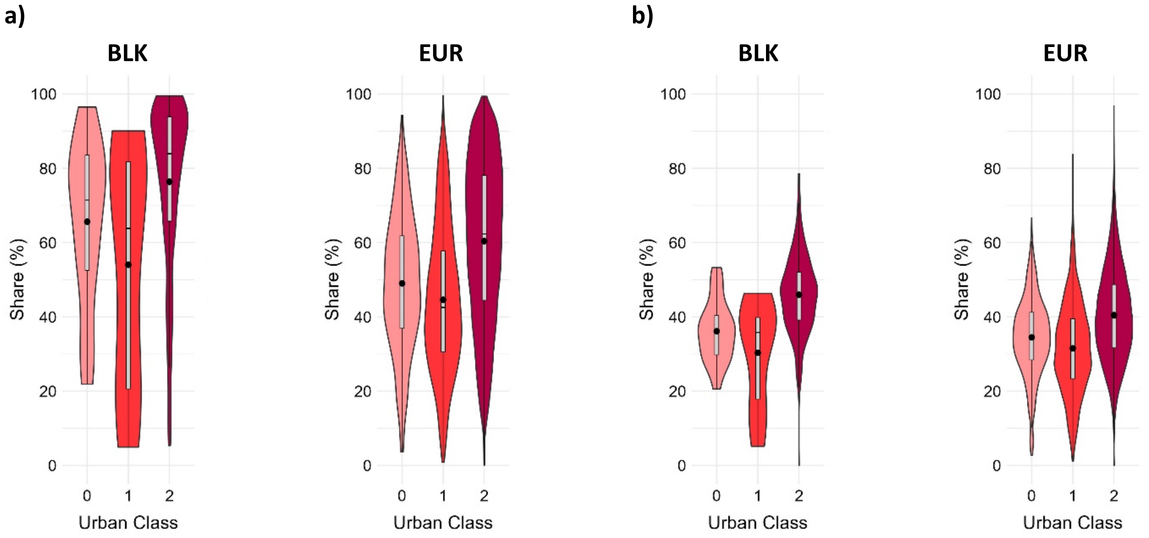


**Figure SM5.13. Statistical distribution of the ecosystem condition variable urban green share in urban ecosystems (a) and in *Settlements and Other Artificial Areas* of urban ecosystems (b) aggregated by urban classes. Violin plots show values for the Western Balkans (BLK)and all European countries (EUR).** Class 0 = local administrative units classified as cities above or equal to 50.000 inhabitants; Class 1 = local administrative units classified as cities below 50.000 inhabitants; Class 2 = local administrative units classified as towns and suburbs.

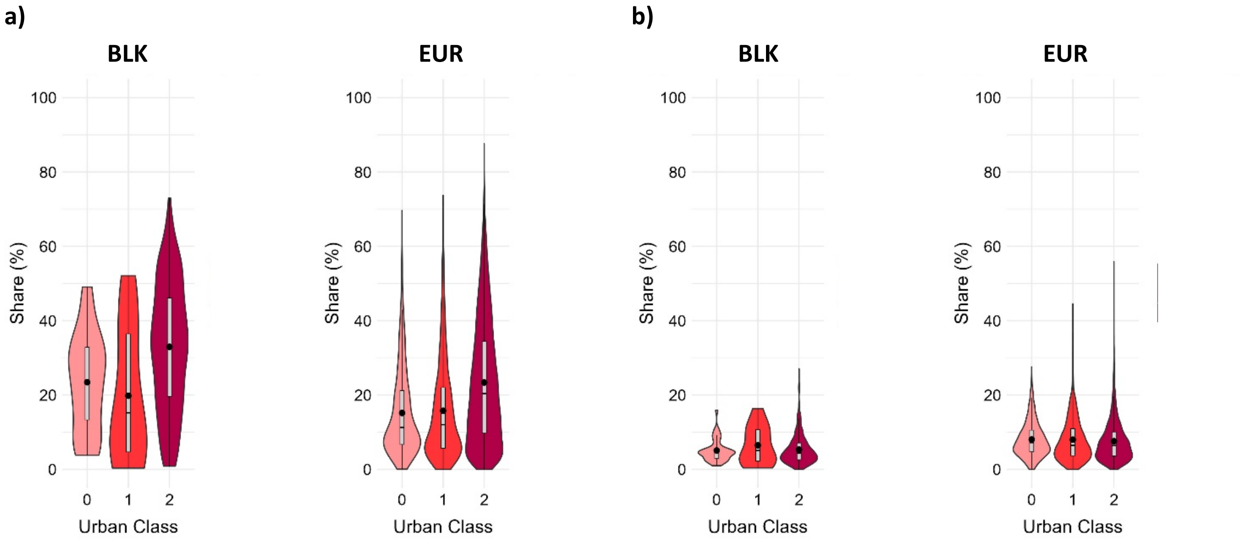


**Figure SM5.14. Statistical distribution of the ecosystem condition variable tree cover share in urban ecosystems (a) and in *Settlements and Other Artificial Areas* of urban ecosystems (b) aggregated by urban classes. Violin plots show values for the Western Balkans (BLK)and all European countries (EUR).** Class 0 = local administrative units classified as cities above or equal to 50.000 inhabitants; Class 1 = local administrative units classified as cities below 50.000 inhabitants; Class 2 = local administrative units classified as towns and suburbs.

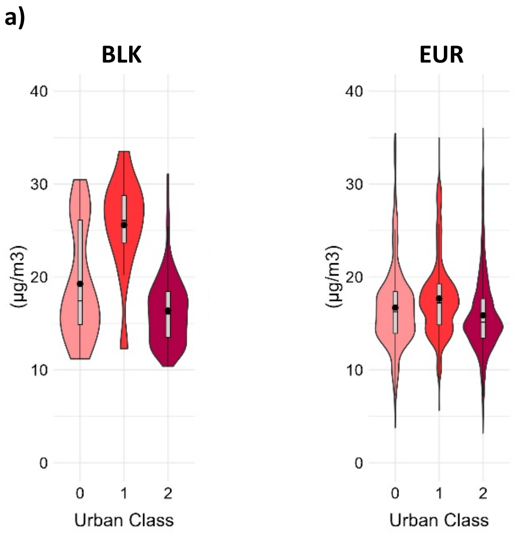


**Figure SM5.15. Statistical distribution of the ecosystem condition variable average annual $PM_{10}$ atmospheric concentration in urban ecosystems aggregated by urban classes. Violin plots show values for the Western Balkans (BLK)and all European countries (EUR).** Class 0 = local administrative units classified as cities above or equal to 50.000 inhabitants; Class 1 = local administrative units classified as cities below 50.000 inhabitants; Class 2 = local administrative units classified as towns and suburbs.

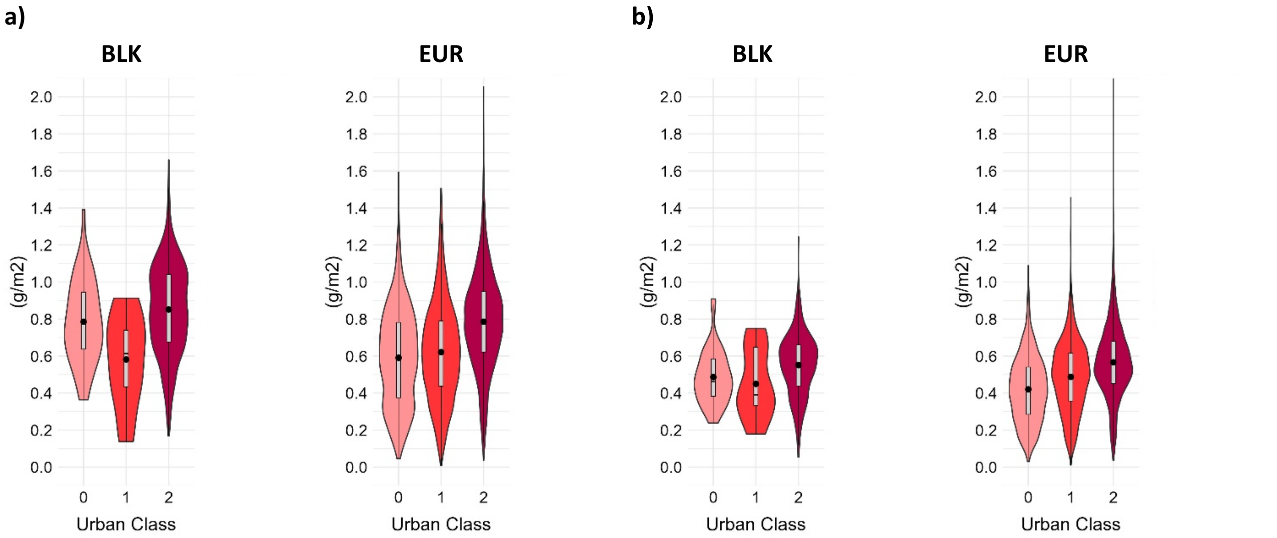


**Figure SM5.16. Statistical distribution of the air filtration service as $PM_{10}$ deposition per land unit (g/m²) in urban ecosystems (a) and in *Settlements and Other Artificial Areas* of urban ecosystems (b) aggregated by urban classes. Violin plots show values for the Western Balkans (BLK)and all European countries (EUR).** Class 0 = local administrative units classified as cities above or equal to 50.000 inhabitants; Class 1 = local administrative units classified as cities below 50.000 inhabitants; Class 2 = local administrative units classified as towns and suburbs.

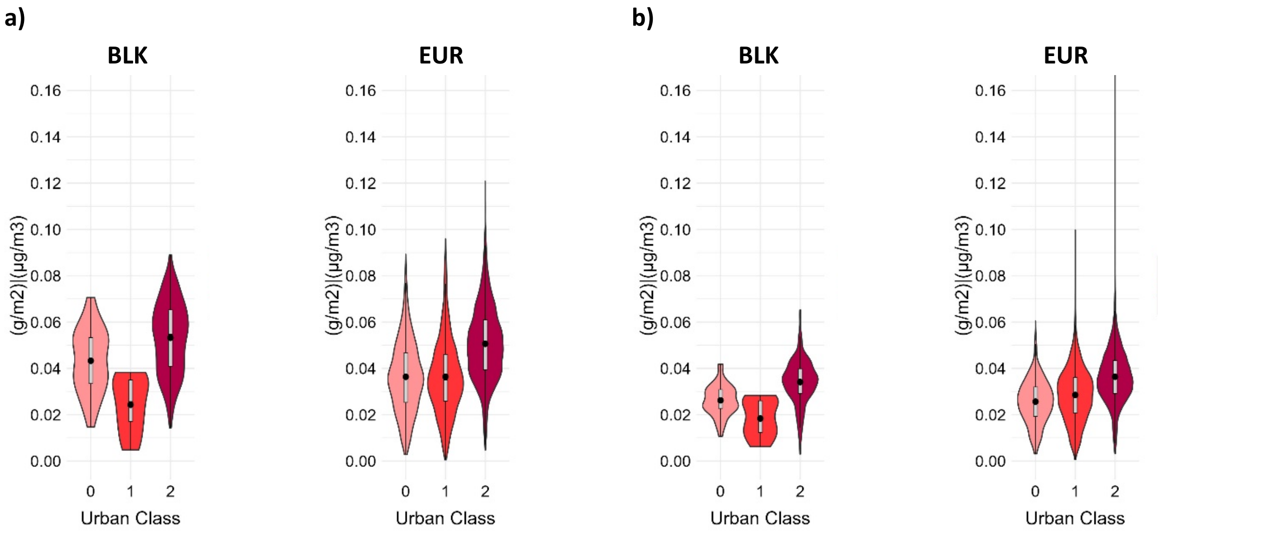


**Figure SM5.17. Statistical distribution of the air filtration service as $PM_{10}$ deposition per land unit (g/m²) normalized by average annual atmospheric $PM_{10}$ levels (µg/m³) in urban ecosystems (a) and in *Settlements and Other Artificial Areas* of urban ecosystems (b) aggregated by urban classes. Violin plots show values for the Western Balkans (BLK)and all European countries (EUR).** Class 0 = local administrative units classified as cities above or equal to 50.000 inhabitants; Class 1 = local administrative units classified as cities below 50.000 inhabitants; Class 2 = local administrative units classified as towns and suburbs.
