## Supplementary Material 6 for "Exploring the value of thematic urban ecosystem accounts in Western Balkan Countries to inform urban greening actions"

**Supplementary Information 6.** Local-level ecosystem extent, condition and air filtration service accounts for the urban ecosystems of the CROSS-REIS and NBFC pilot cities.

**SM6.1 Thematic urban ecosystem extent account in each pilot municipality for the year 2018.** Values are provided in km2. Values are rounded to whole numbers. National-level values and regional aggregated values are also provided as a reference.

| | Settlements | Cropland | Grassland | Forest & Woodland | Heathland | Sp. Veget. | Inl. Wetland | Rivers & Ch. | Lake & Res. | Mar. Inllets | Coastal B. | Marine |
|---|---|---|---|---|---|---|---|---|---|---|---|---|
| **AL** | 512 | 3252 | 896 | 2990 | 867 | 517 | 26 | 39 | 37 | 7 | 54 | 17 |
| **HR** | 1031 | 4033 | 749 | 4494 | 160 | 66 | 16 | 67 | 54 | 0,3 | 4 | 51 |
| **ME** | 228 | 1463 | 769 | 6117 | 43 | 642 | 109 | 6 | 241 | 1 | 18 | 8 |
| **RS** | 1850 | 22834 | 2558 | 18914 | 11 | 187 | 195 | 286 | 131 | - | - | - |
| **SI** | 376 | 1317 | 326 | 3071 | 32 | 39 | 2 | 13 | 12 | - | 7 | 1 |
| **TI (AL)** | 68,30 | 299,02 | 103,68 | 421,42 | 132,14 | 77,78 | - | - | 7,24 | - | - | - |
| **PO (ME)** | 89,13 | 227,74 | 82,45 | 691,34 | 0,16 | 194,80 | 70,48 | 1,99 | 101,12 | - | - | - |
| **NIŠ (RS)** | 61,85 | 291,98 | 21,92 | 215,28 | - | - | 2,64 | 3,03 | - | - | - | - |
| **KR (SI)** | 16,19 | 46,94 | 6,97 | 80,59 | - | 0,08 | - | - | 0,42 | - | - | - |
| **RI (HR)** | 26,14 | 0,68 | 0,60 | 15,19 | 0,01 | 0,32 | - | - | - | - | - | 0,21 |
| **CA (IT)** | 11,47 | 38,08 | 0,02 | 6,36 | - | - | - | - | - | - | - | - |
| **RO (IT)** | 414,19 | 755,76 | 9,70 | 78,14 | 16,49 | 1,58 | - | 8,19 | 1,49 | - | 2,20 | 0,67 |
| **FI (IT)** | 52,58 | 44,80 | - | 3,10 | - | - | - | 1,84 | - | - | - | - |
| **TO (IT)** | 98,46 | 16,94 | 0,62 | 11,43 | - | 0,55 | - | 2,57 | - | - | - | - |
| **NA (IT)** | 88,69 | 22,02 | 0,86 | 4,92 | 0,94 | 0,77 | - | - | - | - | - | 1,06 |
| **MI (IT)** | 141,59 | 38,91 | 0,35 | 0,41 | - | - | - | - | 0,72 | - | - | - |
| **IT** | 12714 | 76576 | 3065 | 21327 | 3589 | 2883 | 123 | 228 | 772 | 308 | 295 | 234 |
| **BLK** | 5466 | 49805 | 10915 | 63208 | 2542 | 2338 | 371 | 524 | 640 | 9 | 83 | 77 |
| **CR** | 3997 | 32900 | 5298 | 35585 | 1112 | 1450 | 348 | 411 | 475 | 9 | 83 | 77 |
| **EUR** | 136467 | 438228 | 125894 | 320412 | 30985 | 14852 | 12458 | 3614 | 28212 | 1553 | 2480 | 1950 |

Note: Complete names of ecosystem types 1 = Settlement and Other Artificial Areas; 2 = Cropland; 3 = Grassland; 4 = Forest and Woodland; 5 = Heathland and Shrubland; 6. = Sparsely Vegetated Ecosystems; 7. = Inland Wetlands; 8 = Rivers and Channels; 9 = Lakes and Reservoirs; 10 = Marine Inlets and Transitional Waters; 11 = Coastal Beaches, Dunes and Wetlands; 12 = Marine.

**SM6.2 Thematic urban ecosystem extent account in each pilot municipality for the year 2018.** Values are provided as share of the national urban ecosystem accounting area occupied by each Ecosystem Type Level 1. Values are rounded to the first decimal.

| | **Settlements** | **Cropland** | **Grassland** | **Forest & Woodland** | **Heathland** | **Sp. Veget.** | **Inl. Wetland** | **Rivers & Ch.** | **Lake & Res.** | **Mar. Inlets** | **Coastal B.** | **Marine** |
|---|---|---|---|---|---|---|---|---|---|---|---|---|
| **AL** | 5,6% | 35,3% | 9,7% | 32,4% | 9,4% | 5,6% | 0,3% | 0,4% | 0,4% | 0,1% | 0,6% | 0,2% |
| **HR** | 9,6% | 37,6% | 7,0% | 41,9% | 1,5% | 0,6% | 0,1% | 0,6% | 0,5% | 0,0% | 0,0% | 0,5% |
| **ME** | 2,4% | 15,2% | 8,0% | 63,4% | 0,4% | 6,7% | 1,1% | 0,1% | 2,5% | 0,0% | 0,2% | 0,1% |
| **RS** | 3,9% | 48,6% | 5,4% | 40,3% | 0,0% | 0,4% | 0,4% | 0,6% | 0,3% | 0,0% | 0,0% | 0,0% |
| **SI** | 7,2% | 25,3% | 6,3% | 59,1% | 0,6% | 0,7% | 0,0% | 0,2% | 0,2% | 0,0% | 0,1% | 0,0% |
| **TI (AL)** | 6,16% | 26,95% | 9,34% | 37,98% | 11,91% | 7,01% | - | - | 0,65% | - | - | - |
| **PO (ME)** | 6,11% | 15,61% | 5,65% | 47,38% | 0,01% | 13,35% | 4,83% | 0,14% | 6,93% | - | - | - |
| **NIŠ (RS)** | 10,37% | 48,93% | 3,67% | 36,08% | - | - | 0,44% | 0,51% | - | - | - | - |
| **KR (SI)** | 10,71% | 31,05% | 4,61% | 53,31% | - | 0,05% | - | - | 0,27% | - | - | - |
| **RI (HR)** | 60,59% | 1,58% | 1,39% | 35,21% | 0,01% | 0,73% | - | - | - | - | - | 0,48% |
| **CA (IT)** | 20,50% | 68,08% | 0,04% | 11,38% | - | - | - | - | - | - | - | - |
| **RO (IT)** | 32,15% | 58,66% | 0,75% | 6,07% | 1,28% | 0,12% | - | 0,64% | 0,12% | - | 0,17% | 0,05% |
| **FI (IT)** | 51,39% | 43,79% | - | 3,03% | - | - | - | 1,80% | - | - | - | - |
| **TO (IT)** | 77,81% | 21,38% | 0,19% | 0,23% | - | - | - | - | 0,40% | - | - | - |
| **NA (IT)** | 74,37% | 18,46% | 0,72% | 4,13% | 0,79% | 0,65% | - | - | - | - | - | 0,89% |
| **MI (IT)** | 75,40% | 12,97% | 0,48% | 8,76% | - | 0,42% | - | 1,97% | - | - | - | - |
| **IT** | 10,41% | 62,71% | 2,51% | 17,46% | 2,94% | 2,36% | 0,10% | 0,19% | 0,63% | 0,25% | 0,24% | 0,19% |
| **BLK** | 4,02% | 36,63% | 8,03% | 46,48% | 1,87% | 1,72% | 0,27% | 0,39% | 0,47% | 0,01% | 0,06% | 0,06% |
| **CR** | 4,89% | 40,25% | 6,48% | 43,53% | 1,36% | 1,77% | 0,43% | 0,50% | 0,58% | 0,01% | 0,10% | 0,09% |
| **EUR** | 12,22% | 39,23% | 11,27% | 28,68% | 2,77% | 1,33% | 1,12% | 0,32% | 2,53% | 0,14% | 0,22% | 0,17% |

Note: Complete names of ecosystem types 1 = Settlement and Other Artificial Areas; 2 = Cropland; 3 = Grassland; 4 = Forest and Woodland; 5 = Heathland and Shrubland; 6. = Sparsely Vegetated Ecosystems; 7. = Inland Wetlands; 8 = Rivers and Channels; 9 = Lakes and Reservoirs; 10 = Marine Inlets and Transitional Waters; 11 = Coastal Beaches, Dunes and Wetlands; 12 = Marine.

**SM6.3 Detailed ecosystem extent account for *Settlements and Other Artificial Areas* within urban ecosystem areas in each pilot municipality for the year 2018.** Values are provided in km2. Values are rounded to two decimals.

| | Sealed | Woody – needleleaved trees (Coniferous) | Woody - Broadleaved deciduous | Woody - Broadleaved evergreen | Low growing woody plants (Shrubs) | Permanent Herbaceous | Periodically Herbaceous | Lichens & Mosses | Sparsely Vegetated | Water | Permanent Snow & Ice |
|---|---|---|---|---|---|---|---|---|---|---|---|
| **AL** | 217 | 4 | 44 | 24 | 8 | 112 | 81 | - | 20 | 1 | - |
| **HR** | 476 | 10 | 95 | 41 | 39 | 295 | 45 | - | 21 | 9 | - |
| **ME** | 74 | 3 | 11 | 35 | 17 | 57 | 13 | - | 16 | 2 | - |
| **RS** | 841 | 10 | 246 | - | 65 | 468 | 148 | - | 59 | 12 | - |
| **SI** | 195 | 6 | 38 | 1 | 5 | 115 | 9 | - | 5 | 3 | - |
| **TI (AL)** | 42,01 | 0,81 | 5,73 | 1,53 | 0,18 | 11,13 | 4,71 | - | 2,11 | 0,09 | - |
| **PO (ME)** | 27,54 | 0,55 | 1,85 | 10,54 | 6,87 | 26,73 | 10,14 | - | 3,62 | 1,30 | - |
| **NIŠ (RS)** | 33,04 | 0,25 | 11,21 | - | 0,76 | 11,97 | 3,74 | - | 0,63 | 0,27 | - |
| **KR (SI)** | 8,91 | 0,25 | 1,29 | - | 0,08 | 4,98 | 0,42 | - | 0,05 | 0,21 | - |
| **RI (HR)** | 15,07 | 0,17 | 4,48 | 0,44 | 1,59 | 3,89 | 0,00 | - | 0,23 | 0,27 | - |
| **CA (IT)** | 6,51 | 0,02 | 0,93 | 0,31 | 0,20 | 2,95 | 0,42 | - | 0,12 | - | - |
| **RO (IT)** | 250,80 | 18,18 | 24,58 | 33,07 | 1,61 | 69,97 | 11,46 | - | 3,39 | 1,12 | - |
| **FI (IT)** | 35,44 | 0,57 | 4,19 | 3,83 | 0,24 | 7,68 | 0,22 | - | 0,23 | 0,18 | - |
| **TO (IT)** | 72,51 | 0,93 | 12,89 | 0,17 | 0,12 | 9,34 | 0,81 | - | 1,15 | 0,54 | - |
| **NA (IT)** | 66,63 | 1,31 | 7,14 | 3,53 | 1,62 | 6,71 | 0,95 | - | 0,50 | 0,29 | - |
| **MI (IT)** | 101,03 | 0,95 | 15,53 | 0,22 | 0,51 | 20,70 | 0,82 | - | 1,56 | 0,27 | - |
| **IT** | 7760 | 145 | 813 | 496 | 191 | 2598 | 374 | - | 268 | 68 | - |
| **BLK** | 2443 | 41 | 569 | 106 | 156 | 1486 | 422 | - | 206 | 37 | - |
| **CR** | 1804 | 33 | 434 | 100 | 134 | 1047 | 296 | - | 120 | 28 | - |
| **EUR** | 72619 | 3237 | 17407 | 1172 | 1575 | 32968 | 3348 | 3 | 3089 | 1049 | - |

**SM6.4 Detailed ecosystem extent account for *Settlements and Other Artificial Areas* within urban ecosystem areas in each pilot municipality for the year 2018.** Values are provided as share of the Settlements and Other Artificial Areas occupied by Corine Land Cover Plus Classes. Values are rounded to the first decimal.

| | Sealed | Woody – needleleaved trees (Coniferous) | Woody - Broadleaved deciduous | Woody - Broadleaved evergreen | Low growing woody plants (Shrubs) | Permanent Herbaceous | Periodically Herbaceous | Lichens & Mosses | Sparsely Vegetated | Water | Permanent Snow & Ice |
|---|---|---|---|---|---|---|---|---|---|---|---|
| **AL** | 42,40% | 0,80% | 8,68% | 4,61% | 1,53% | 21,91% | 15,90% | 0,00% | 3,89% | 0,28% | - |
| **HR** | 46,18% | 1,00% | 9,21% | 4,02% | 3,74% | 28,63% | 4,35% | 0,00% | 2,00% | 0,86% | - |
| **ME** | 32,60% | 1,23% | 4,68% | 15,24% | 7,65% | 24,85% | 5,72% | 0,00% | 6,94% | 1,09% | - |
| **RS** | 45,44% | 0,53% | 13,31% | 0,00% | 3,51% | 25,30% | 8,02% | 0,00% | 3,20% | 0,67% | - |
| **SI** | 52,01% | 1,62% | 10,08% | 0,17% | 1,39% | 30,55% | 2,28% | 0,00% | 1,21% | 0,69% | - |
| **TI (AL)** | 61,50% | 1,19% | 8,38% | 2,24% | 0,27% | 16,30% | 6,89% | - | 3,09% | 0,14% | - |
| **PO (ME)** | 30,90% | 0,62% | 2,07% | 11,82% | 7,71% | 29,99% | 11,38% | - | 4,06% | 1,46% | - |
| **NIŠ (RS)** | 53,42% | 0,41% | 18,12% | - | 1,22% | 19,35% | 6,04% | - | 1,02% | 0,43% | - |
| **KR (SI)** | 55,08% | 1,52% | 7,96% | - | 0,50% | 30,74% | 2,57% | - | 0,31% | 1,31% | - |
| **RI (HR)** | 57,65% | 0,64% | 17,13% | 1,68% | 6,08% | 14,89% | 0,01% | - | 0,88% | 1,03% | - |
| **CA (IT)** | 56,79% | 0,19% | 8,07% | 2,74% | 1,75% | 25,75% | 3,69% | - | 1,02% | - | - |
| **RO (IT)** | 60,55% | 4,39% | 5,94% | 7,98% | 0,39% | 16,89% | 2,77% | - | 0,82% | 0,27% | - |
| **FI (IT)** | 67,41% | 1,08% | 7,96% | 7,28% | 0,46% | 14,60% | 0,41% | - | 0,44% | 0,35% | - |
| **TO (IT)** | 71,35% | 0,67% | 10,97% | 0,15% | 0,36% | 14,62% | 0,58% | - | 1,10% | 0,19% | - |
| **NA (IT)** | 75,12% | 1,48% | 8,05% | 3,98% | 1,83% | 7,57% | 1,08% | - | 0,57% | 0,33% | - |
| **MI (IT)** | 73,65% | 0,94% | 13,09% | 0,18% | 0,12% | 9,49% | 0,82% | - | 1,17% | 0,54% | - |
| **IT** | 61,03% | 1,14% | 6,40% | 3,90% | 1,50% | 20,43% | 2,94% | - | 2,11% | 0,54% | 0,00% |
| **BLK** | 44,69% | 0,74% | 10,41% | 1,95% | 2,85% | 27,18% | 7,72% | - | 3,78% | 0,68% | - |
| **CR** | 45,13% | 0,83% | 10,87% | 2,51% | 3,35% | 26,19% | 7,41% | - | 3,01% | 0,70% | - |
| **EUR** | 53,21% | 2,37% | 12,76% | 0,86% | 1,15% | 24,16% | 2,45% | 0,00% | 2,26% | 0,77% | 0,00% |

**SM6.5 Thematic urban ecosystem condition account in each pilot municipality for the year 2018.** Values are only provided for the entire national urban ecosystem accounting area, independently of the Ecosystem Type Level 1, and for the Ecosystem Type Settlements and Other Artificial Areas within the national urban ecosystem areas. Values are provided as shares, except for PM10 atmospheric concentration, where values are provided in µg/m$^3$.

| | Imperviousness | | Urban Green | | Urban Tree Cover | | Average annual $PM_{10}$ atmospheric concentration |
|---|---|---|---|---|---|---|---|
| | Overall Urban Ecosystem | Settlements and Other Artificial Areas | Overall Urban Ecosystem | Settlements and Other Artificial Areas | Overall Urban Ecosystem | Settlements and Other Artificial Areas | Overall Urban Ecosystem |
| **AL** | 1,8% | 26,9% | 72,7% | 37,5% | 22,2% | 3,4% | 12,9 |
| **HR** | 3,9% | 32,6% | 74,2% | 46,6% | 31,9% | 5,3% | 15,6 |
| **ME** | 0,7% | 20,6% | 93,7% | 50,7% | 34,9% | 5,2% | 11,2 |
| **RS** | 1,4% | 28,1% | 71,0% | 42,7% | 30,7% | 5,5% | 16,7 |
| **SI** | 3,7% | 39,0% | 87,5% | 43,8% | 49,4% | 6,5% | 15,4 |
| **TI (AL)** | 3,0% | 43,5% | 80,6% | 28,4% | 29,5% | 4,1% | 12,8 |
| **PO (ME)** | 1,5% | 20,8% | 84,8% | 52,2% | 19,0% | 2,3% | 11,2 |
| **NIŠ (RS)** | 4,3% | 35,0% | 72,5% | 39,1% | 29,8% | 5,4% | 16,3 |
| **KR (SI)** | 5,5% | 41,2% | 77,9% | 40,7% | 45,3% | 5,8% | 16,3 |
| **RI (HR)** | 28,2% | 44,6% | 61,6% | 40,4% | 22,6% | 8,4% | 14,1 |
| **CA (IT)** | 10,5% | 40,5% | 57,2% | 38,5% | 14,8% | 6,4% | 14,1 |
| **RO (IT)** | 17,2% | 45,4% | 47,0% | 35,6% | 13,9% | 9,9% | 15,9 |
| **FI (IT)** | 27,1% | 49,4% | 57,8% | 31,4% | 19,7% | 8,7% | 14,6 |
| **TO (IT)** | 46,7% | 58,3% | 29,5% | 26,8% | 7,1% | 6,8% | 29,4 |
| **NA (IT)** | 50,9% | 63,1% | 34,4% | 22,9% | 14,8% | 9,2% | 18,5 |
| **MI (IT)** | 45,7% | 59,0% | 36,4% | 23,8% | 19,3% | 10,1% | 22,8 |
| **IT** | 6,6% | 46,4% | 61,9% | 33,4% | 21,2% | 6,3% | 16,7 |
| **BLK** | 1,5% | 29,2% | 80,1% | 43,0% | 33,9% | 5,0% | 15,60 |
| **CR** | 1,8% | 29,7% | 75,3% | 43,6% | 31,6% | 5,3% | 15,38 |
| **EURr** | 5,8% | 38,9% | 64,5% | 41,3% | 24,1% | 9,0% | 13,44 |

**SM6.6 Thematic urban ecosystem service air filtration (Deposition of $PM_{10}$ in urban vegetation) account in pilot municipality for the year 2018.** Values are provided only for the total national urban ecosystem accounting area, irrespective of Ecosystem Type Level 1, and for the Ecosystem Type *Settlements and Other Artificial Areas* within national urban ecosystem areas. Reported values include: total $PM_{10}$ deposited (tonnes, rounded to whole numbers); average $PM_{10}$ deposition per unit area (g/m², rounded to two decimals); and average $PM_{10}$ deposition per unit area normalized by $PM_{10}$ concentration (g/m² per µg/m³, rounded to two decimals).

| | **Total Mass of $PM_{10}$ deposited** | | **Average mass of $PM_{10}$ deposited per land unit (g/$m_2$),** | | **Average mass of $PM_{10}$ deposited per land unit (g/$m_2$) normalized by $PM_{10}$ concentration (in µg/m³).** | |
|---|---|---|---|---|---|---|
| | **Overall Urban Ecosystem** | **Settlements and Other Artificial Areas** | **Overall Urban Ecosystem** | **Settlements and Other Artificial Areas** | **Overall Urban Ecosystem** | **Settlements and Other Artificial Areas** |
| **AL** | 6197 | 250 | 0,67 | 0,49 | 0,05 | 0,04 |
| **HR** | 9270 | 589 | 0,86 | 0,57 | 0,06 | 0,04 |
| **ME** | 6973 | 97 | 0,72 | 0,43 | 0,06 | 0,04 |
| **RS** | 43676 | 1134 | 0,93 | 0,61 | 0,06 | 0,04 |
| **SI** | 5451 | 237 | 1,05 | 0,63 | 0,07 | 0,04 |
| **TI (AL)** | 824 | 26 | 0,74 | 0,38 | 0,06 | 0,03 |
| **PO (ME)** | 812 | 35 | 0,56 | 0,39 | 0,05 | 0,04 |
| **NIŠ (RS)** | 524 | 29 | 0,88 | 0,46 | 0,05 | 0,03 |
| **KR (SI)** | 165 | 10 | 1,09 | 0,64 | 0,07 | 0,04 |
| **RI (HR)** | 26 | 12 | 0,60 | 0,45 | 0,04 | 0,03 |
| **CA (IT)** | 35 | 4 | 0,63 | 0,39 | 0,04 | 0,03 |
| **RO (IT)** | 1008 | 202 | 0,78 | 0,49 | 0,05 | 0,03 |
| **FI (IT)** | 54 | 19 | 0,53 | 0,35 | 0,04 | 0,02 |
| **TO (IT)** | 73 | 37 | 0,66 | 0,53 | 0,02 | 0,02 |
| **NA (IT)** | 54 | 31 | 0,46 | 0,35 | 0,02 | 0,02 |
| **MI (IT)** | 120 | 76 | 0,56 | 0,38 | 0,02 | 0,02 |
| **IT** | 95867 | 6623 | 0,79 | 0,52 | 0,05 | 0,03 |
| **BLK** | 110395 | 3013 | 0,81 | 0,55 | 0,05 | 0,04 |
| **CR** | 71567 | 2308 | 0,88 | 0,58 | 0,06 | 0,04 |
| **EUR** | 724615 | 68422 | 0,65 | 0,50 | 0,05 | 0,04 |
